# Commonality and Variability in Functional Networks in Children Under 5 Years Old

**DOI:** 10.1101/2025.09.12.675913

**Authors:** Jiaxin Cindy Tu, Chenyan Lu, Trevor K. M. Day, Robert Hermosillo, Lucille A. Moore, Anxu Wang, Xintian Wang, Donna Dierker, Aidan Latham, Jeanette K. Kenley, Damien A. Fair, Jed T. Elison, Chad M. Sylvester, Barbara B. Warner, Joan L. Luby, Cynthia E. Rogers, Deanna M. Barch, Christopher D. Smyser, Timothy O. Laumann, Evan M. Gordon, Adam T. Eggebrecht, Muriah D. Wheelock

**Author notes:** **Corresponding Author:** Muriah Wheelock, Mallinckrodt Institute of Radiology 4525 Scott Ave, St. Louis, MO 63110.

## Abstract

Functional brain networks support human cognition, yet how individualized network architecture emerges in early childhood remains poorly understood. Averaging across participants can obscure age-specific organization and person-to-person differences, particularly in slowly developing association cortices. We developed an age-appropriate functional reference that captured common structure across toddlers without averaging away individual variability, enabling estimation of each child’s networks from resting-state fMRI.

Across cohorts of 8–60-month-old children, we found individualized network organization—including finer-scale subdivisions and emerging language lateralization—well before age five. Network layouts showed longitudinal stability, with greater consistency in sensory than association regions. Within-network connectivity was stronger and explained age-related variance when networks were defined using individualized rather than group-consensus topography. Left-lateralization of language networks tracked age-normalized verbal ability, linking early functional architecture to emerging cognition. These findings show that behaviorally relevant brain networks arise far earlier than previously recognized, providing a foundation for studying typical development and early biomarkers.

## Introduction

Functional network patterns in young children (< 5 years) have been characterized at the group level (Eggebrecht et al., 2017; Kardan et al., 2022; Myers et al., 2024; Sylvester et al., 2022; Tu et al., 2025a; Wang et al., 2023). However, these group-average pediatric brain networks may not comprehensively represent the organization of the developing brain. Recent studies in adults have demonstrated idiosyncratic yet reliable features in functional network topography across individuals that are qualitatively different from group-average estimates (Bijsterbosch et al., 2023; Braga and Buckner, 2017; Dworetsky et al., 2021; Gordon et al., 2017c, 2017b; Gratton et al., 2018; Laumann et al., 2015). These trait-like topographical features are stable across both sessions (Seitzman et al., 2019) and task states (Kraus et al., 2021), and may be related to individual behavioral characteristics (Cui et al., 2020) or clinical phenotypes (Lynch et al., 2024). Group-averaging procedures not only fail to capture these individual-specific features, but they also tend to obscure networks with highly variable spatial topographies across individuals (Ladwig et al., 2025). Therefore, we reasoned that using an approach that circumvents the group-averaging could potentially reveal finer network details in a pediatric population, which may reveal unique developmental changes and variability.

Reliable identification of individual-specific functional networks with unsupervised clustering/community detection techniques often requires extended data acquisition. For example, with the commonly used Infomap algorithm (Power et al., 2011; Rosvall and Bergstrom, 2008), more than 90-100 minutes of data has been recommended (Gordon et al., 2017c; Laumann et al., 2015). However, starting these analyses de novo for every new individual might not be necessary, given that most resting-state functional network topography is consistent across individuals (Damoiseaux et al., 2006; Dworetsky et al., 2021; Gratton et al., 2018). Consequently, several prior-based methods have been developed to derive individual-specific functional networks in populations with limited data available per individual (Gordon et al., 2017a; Hacker et al., 2013; Hermosillo et al., 2024; Kong et al., 2019; Li et al., 2019; Luckett et al., 2023; Wang et al., 2015), significantly lowering the data requirement (Hermosillo et al., 2024; Kong et al., 2019).

Although the aforementioned prior-based methods have significantly advanced the mapping of individual-specific network organizations, the selection of network priors remains a critical decision that has been frequently overlooked, especially when applying these methods to populations beyond healthy, young adults. For example, previous work has used a set of adult network priors for data acquired in children younger than 5 years old (Moore et al., 2024; Sun et al., 2025), which assumes that the number and spatial topography of functional networks in pediatric cohorts are reasonably similar to the adult population average. However, mounting evidence suggests functional connectivity (FC) and network features in young children consistently differ from older children and adults (Eggebrecht et al., 2017; Fransson et al., 2007; Gao et al., 2015a, 2015b; Kardan et al., 2022; Marrus et al., 2018; Myers et al., 2024; Sylvester et al., 2022; Tu et al., 2025a, 2025b; Wang et al., 2023). Notably, while some studies have proposed deriving infant-specific network templates by averaging BOLD time series at cortical locations defined by adult networks to calculate network-average FC profiles (Moore et al., 2024), it remains unclear whether this approach necessarily yields truly “age-specific” networks. To date, few studies have delineated individual-specific networks in children younger than 5 years old using age-specific group priors (Derman et al., 2025).

Additionally, hemispheric lateralization of functional networks has been consistently observed in prior literature, especially in heteromodal association cortices (Wang et al., 2014, 2015; Perez et al., 2023). For example, the language network exhibits a left-hemisphere bias (Braga et al., 2020; Lipkin et al., 2022; Wang et al., 2015), while the ventral attention network shows a right-hemisphere bias (Bernard et al., 2020; Wang et al., 2015). Furthermore, left lateralization in the language network strengthens from childhood into young adulthood (Holland et al., 2007; Lidzba et al., 2011; Olulade et al., 2020; Sepeta et al., 2016; Szaflarski et al., 2006b, 2006a), with individual differences in network lateralization linked to handedness (Perez et al., 2023; Wang et al., 2015). However, it is unclear how early hemispheric lateralization emerges in young children and whether it is associated with the development of behavioral functions such as language.

Here, we demonstrate functional network organization variations across both individuals and chronological age in children at 1 to 5 years old using two datasets and fine network divisions representative in young children. Specifically, we first identified group-level network templates using high-quality scan sessions with >20 minutes of low-motion data at 2 and 3 years and then created vertex-level individual-specific networks using fMRI data from a single scan session. This approach enabled inclusion of network details not previously described in group data in this age range and provided a prior for noise-robust subject-specific network localization. Our analyses show that using age-specific network priors supports individual-specific network organizations can be determined using 10-20 minutes of low-motion fMRI data in young children. The population consensus assignment from individual network maps produced a cleaner and finer group-level division than networks found on group-average FC data. Furthermore, lateralization in the language network increased with age, with greater lateralization positively associated with age-normalized verbal developmental quotient. Taken together, our results offer insights into commonality and variability in functional network topography in young children before 5 years and provide insights into functional specialization in the developing brain.

## Results

### Identification of age-appropriate functional connectivity patterns using concatenated individual data

Previously, we identified functional networks based in 2-year-old children using group-averaged functional connectivity (FC) computed within an age-specific area parcellation (Tu et al., 2025a). While this approach captures dominant population-level organization, anatomically matched cortical areas can exhibit distinct FC profiles across individuals during early development. As a result, averaging FC across participants may obscure both inter-individual variability and connectivity patterns that are consistently expressed only in subsets of children (Supplementary Figure 1; see Supplementary Text). To mitigate this limitation, we identified age-appropriate functional connectivity patterns by clustering concatenated individual FC profiles, rather than group-averaged connectivity. Using the same area parcellation as in prior work (Tu et al., 2025a), FC was computed between all pairs of parcels within each session, yielding parcel-wise FC profiles in which each parcel served both as a feature (connectivity target) and as a sample, pooled across individuals. Concatenating these parcel-wise FC profiles across participants allows reproducible connectivity patterns to emerge without requiring them to be present in every individual. Using this approach, we clustered FC profiles from 81 fMRI sessions (72 unique children aged 2–3 years, including 9 children with longitudinal data at Y2 and Y3; all sessions >20 minutes of low-motion data; eLABE Dataset 1; Supplementary Figure 2), analogous to prior literature (Yeo et al., 2011; Zhu et al., 2025). A split-half stability analysis revealed a local peak at K = 23 clusters (Supplementary Figure 3). Full methodological details are provided in Materials and Methods.

### Individual-specific network mapping

The resulting 23 resting-state network profiles represent reproducible functional connectivity patterns observed across subjects at 2 to 3 years. The cluster centroids therefore serve as age-appropriate reference FC profiles or “network templates” (example in Figure 1F;Supplementary Figure 4). These reference profiles were subsequently used to delineate individual-specific network organization at finer spatial resolution. Specifically, vertex-level FC profiles (vertex-to-parcel connectivity maps, Figure 1C) were compared with the 23 reference profiles using Pearson’s correlation, and each vertex was assigned to the most similar network using a winner-take-all procedure, following prior work (Gordon et al., 2017a). This step enables detailed vertex-wise network mapping (Figure 1D) while maintaining correspondence with the age-appropriate connectivity patterns identified above.

**Figure 1.**
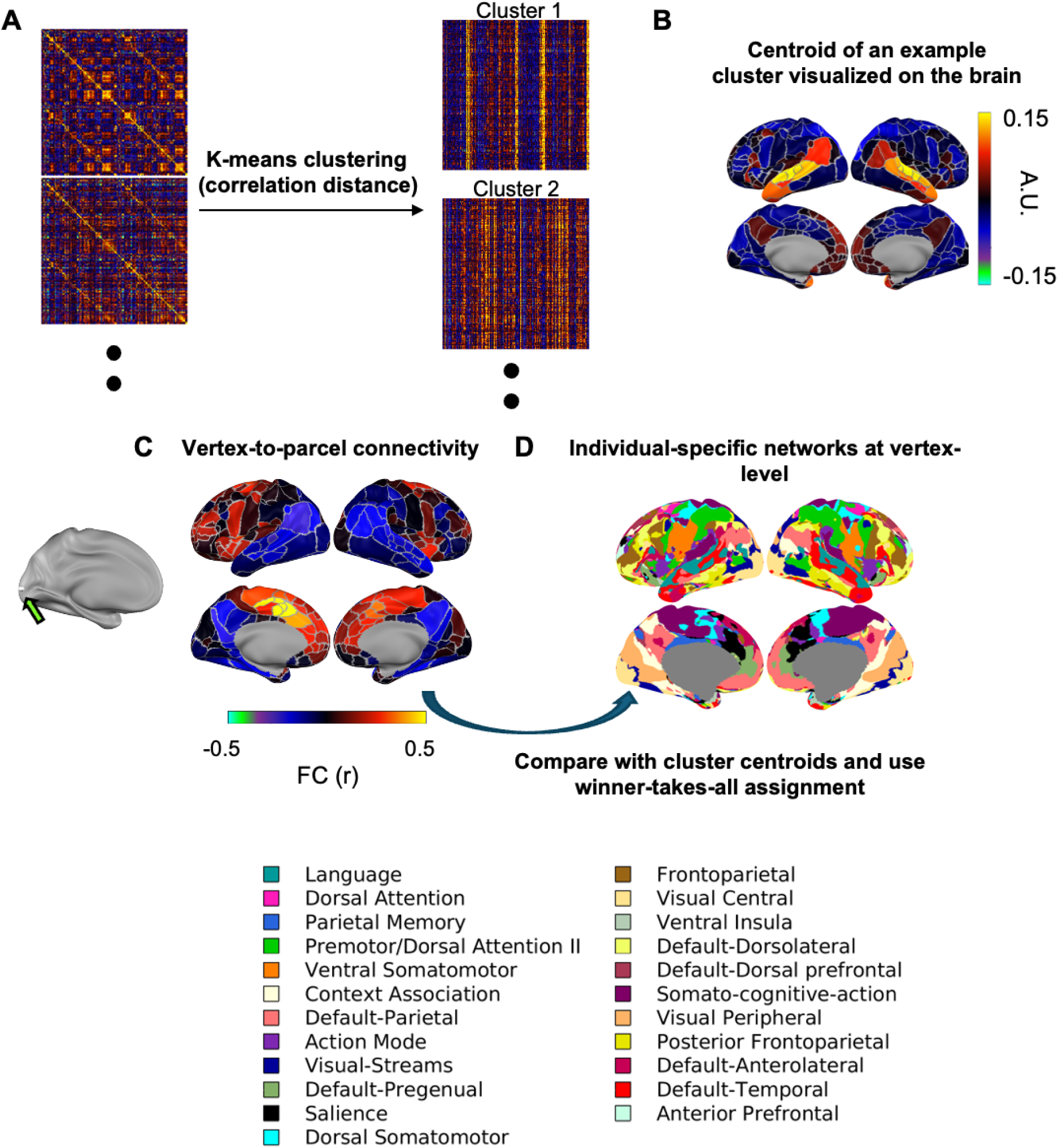
Creation of group-level reference template and individual-specific network mapping. A) Example FC matrices being concatenated and clustered into internally similar clusters of FC profiles. B) Example template FC profiles (centroids) for the Language network. C) Vertex-to-parcel connectivity. D) Individual-specific networks at vertex-level. Network names and colors were chosen with reference to prior literature.

The 23 networks match the fine network divisions in adults and refined our understanding of functional networks in young children.

Network spatial probability plots (Dworetsky et al., 2021; Hermosillo et al., 2024; Lipkin et al., 2022; Lynch et al., 2024) show the most and least common locations of a network across the 81 template sessions (Figure 2). To help put the networks into context and relate them to potential behavioral significance, we further applied the Network Correspondence Toolbox (Kong et al., 2025) and “NiMARE: Neuroimaging Meta-Analysis Research Environment” version 0.5.0 (Salo et al., 2023) to compare the topography of our 23 networks to prior literature (Supplementary Figures 5-16, Supplementary Text). Because task-based fMRI or multimodal data were not available in the present study, functional interpretations of the identified networks are necessarily indirect and based on spatial correspondence with prior reports rather than direct functional validation. Despite the age difference between young children and adults, some of the 23 functional networks have a spatial topography that overlaps with previously identified fine network divisions in adult studies (Du et al., 2024; Gordon et al., 2020; Kong et al., 2019; Lynch et al., 2024). Additionally, the location of less common network occurrences were also consistent with results using highly sampled resting-state fMRI data. For example, we found that the salience network occasionally extends to regions normally assigned to the parietal memory network in some individuals, consistent with a prior report in adults (Kwon et al., 2025). The interdigitation of the Salience (black) and Default-Pregenual (pistachio green) networks at the frontal operculum and the medial prefrontal cortex also resembled the interdigitation of the Salience (black) and Default Mode (red) networks in resting-state fMRI from highly-sampled adult individuals (Braga and Buckner, 2017; Gordon et al., 2017c; Lynch et al., 2024).

**Figure 2.**
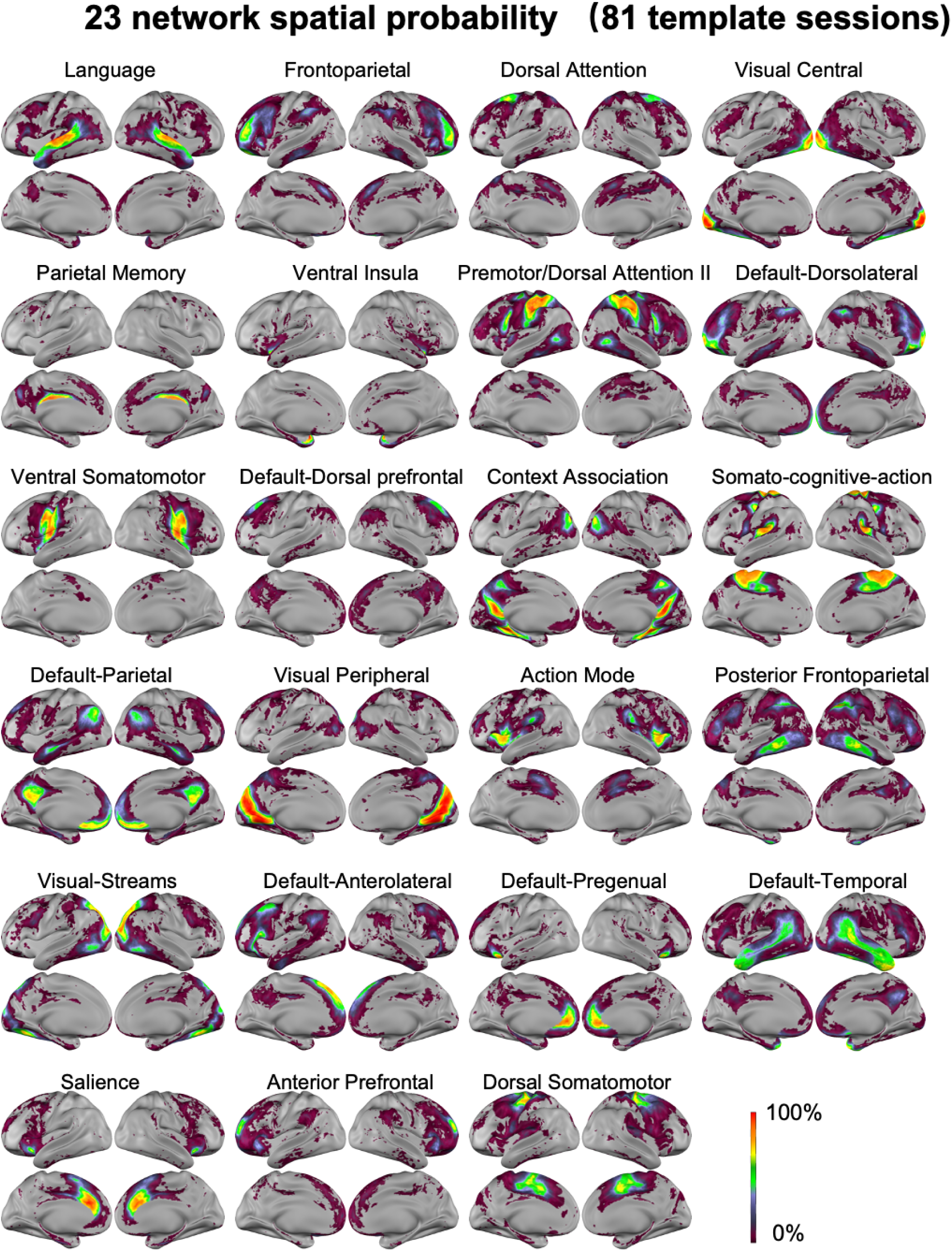
Functional networks spatial probability. For each vertex, the spatial probability of the network is quantified as the percentage of sessions within the 81 template sessions with that matching network assignment.

To further compare and group the 23 fine network divisions identified here with prior reports in adults, we quantified the similarity between each network to the 20 finely divided adult networks from 45 highly sampled adult individuals (Lynch et al., 2024) based on both network spatial probability maps (Figure 2) as well as the average network FC obtained from averaging FC from all vertices assigned to that network in each subject, and then averaged across subjects (Supplementary Figure 17). The main reason these finely divided adult networks were chosen was that the spatial probability map and average FC in Lynch et al. 2024 was created from highly sampled precision fMRI data with a finer division than other group-based network identifications, allowing us to test whether our fine divisions correspond to prior adult literature or they represent children-specific networks. Although most of the 23 networks had clear correspondence with the 20 adult networks, some of them (e.g., “Anterior Prefrontal”, “Default-Dorsal prefrontal” and “Default-Temporal”) did not, potentially suggesting some networks that are present in childhood are later incorporated into other networks in adulthood (Supplementary Figure 18-19). Additionally, there is a much smaller Action Mode Network in young children compared to adults, especially around the dorsal anterior cingulate cortex. Parts of the ventral insula and dorsal somatomotor network in young children might later be incorporated into the Action Mode Network (Supplementary Figure 19). Furthermore, while the frontoparietal network was divided into more cohesive pieces in other studies (Du et al., 2024; Yeo et al., 2011), there was less anterior-posterior bias in the divisions than those reported here. To aid interpretation of the function of these networks and put our networks in the context of existing findings, we made a table summary of previous nomenclature applied to similar network topography in the literature (Table 1), with the caveat that this is only a guideline and some networks may not perfectly overlap with what we described here. This table was made both with the quantitative spatial and connectivity correspondence measures mentioned above, and with expert visual assessment of spatial correspondence, used to complement the quantitative measures.

**Table 1.**
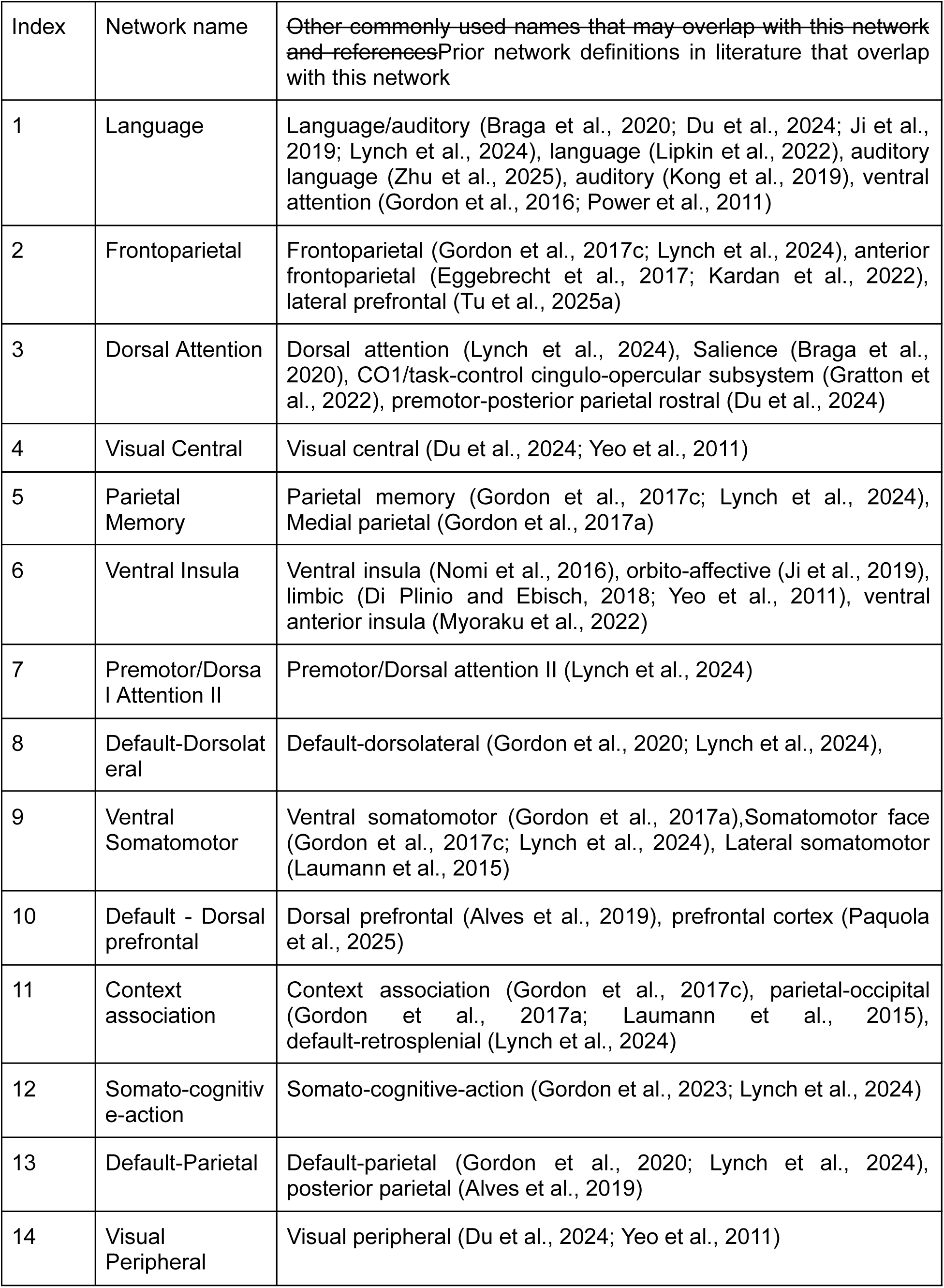
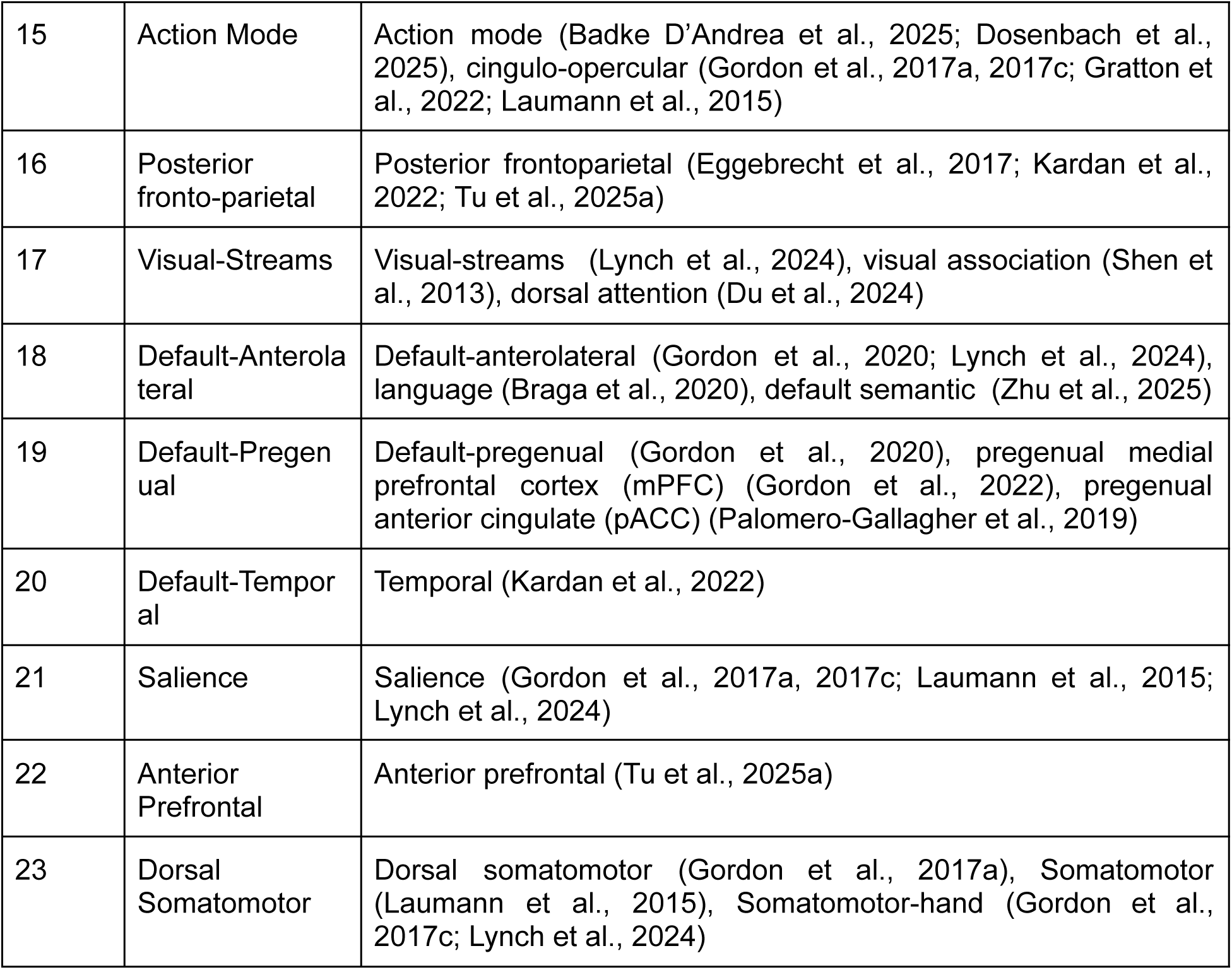
Network names and prior mentions.

### Individual-specific functional networks in young children are stable within sessions and across longitudinal sessions one year apart

To examine the individual-specificity of these functional networks, we obtained the individual functional network assignments using the subset of 49 individuals in Dataset 1 with two longitudinal sessions at Y2 and Y3 time points (2.08 ± 0.12 and 3.16 ± 0.24 years old, respectively) with the same procedure described above (Figure 1). First, we derived functional networks from the first half and second half of each fMRI session. The consistency of network assignment across the split halves (measured with normalized mutual information, NMI) was significantly higher (two-sample t-test p<0.001) within subject (Y2: NMI = 0.4822 ± 0.0669, Y3: NMI = 0.4919 ± 0.0686) than between subjects (Y2: NMI = 0.3274 ± 0.0383, Y3: NMI = 0.3250 ± 0.0379) (Figure 3A, 3C-F), with NMI higher within subject in almost all subjects (Y2: 49, Y3: 47 out of 49), suggesting that the network assignment contain individual-unique features. As a sanity check, the NMI between the network assignments in the first half and the randomly permuted network assignments in the second half is much lower, at approximately 0.001: for Figure 3C, the 95% confidence interval of the permutation null is [0.0013, 0.0016]. To address the potential inflation of NMI by including frames from the same run in the two split halves, we repeated the analysis without overlapping runs and the within-individual NMI was still significantly higher (Supplementary Figure 20, Supplementary Text). To enhance the robustness of our results, we conducted the split-half network reliability analysis in an additional dataset of 8-60 months’ children from the Baby Connectome Project (Dataset 2, Supplementary Figure 21). Qualitatively similar results with higher within-subject NMI were obtained (Supplementary Figure 22A-C). Within-subject NMI was positively correlated with the amount (in minutes) of low-motion data in each split-half (r = 0.6447, p < 0.001), with the total number of minutes of scan time (r = 0.5781, p < 0.001), and was negatively correlated with the mean framewise displacement of the scan session (r = −0.1336, p = 0.0204) but was not significantly correlated with age (r = −0.0553, p = 0.3390;Supplementary Figure 22D). This suggests that the ability to retrieve reliable networks was somewhat limited by the data amount and quality.

**Figure 3.**
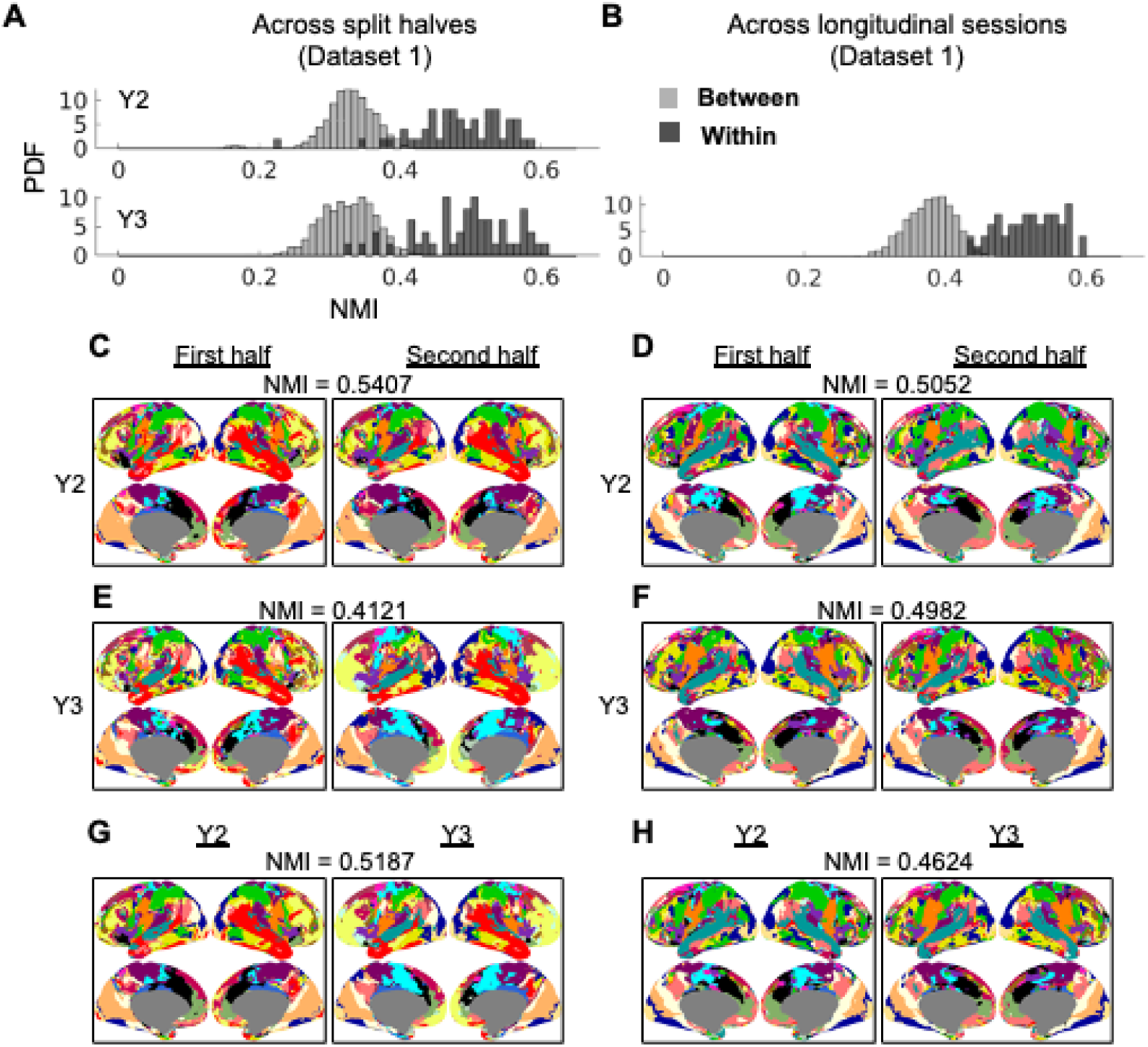
Individual-specific network topography for 23 networks (Dataset 1, 49 individuals with longitudinal sessions). A) Within-versus between-individual NMI (across split halves) at Y2 and Y3 time points. PDF: probability density function. B) Within-versus between-individual NMI (across longitudinal sessions). C-D) Network assignments for example individuals at Y2. E-F) Same as C-E but at Y3. G-H) Network assignments for example individuals across the Y2 and Y3 time points.

To assess longitudinal stability of the individual-specific networks, we also derived individual-specific functional networks from the whole fMRI session for Y2 and Y3 time points of the data collection. Because of the relationship between NMI and amount of data reported above, we decided to use the whole session (Y2: 19.0 ± 4.1 minutes low-motion data, Y3: 18.7 ± 3.7 minutes low-motion data) for this analysis, unlike the prior split-half analysis which used about half of the data amount to generate the networks. The consistency of network assignment across the two longitudinal sessions was significantly higher (two-sample t-test p<0.001) within subject (NMI = 0.5190 ± 0.0444) than between subjects (NMI = 0.3788 ± 0.0330) (Figure 3B, G-H), and the NMI was higher within subject for all 49 subjects, indicating the individual-specific networks were stable from 2 to 3 years old. While the within-subject NMI values are relatively modest, they are comparable to results from a similar template matching method—albeit using different templates and a different similarity metric—applied to the MSC dataset (a high-quality adult dataset with 5 hours of resting-state fMRI). In that study (Hermosillo et al., 2024), within-subject NMI ranged from 0.527 to 0.648 using all available low-motion data after frame censoring, and was around 0.5 using only 15 minutes of contiguous low-motion data. To further explore the amount of data required to achieve reliability and individual-specificity of the networks, we conducted additional analysis with different lengths of contiguous low-motion data (minimum 2.5 minutes and maximum of 20 minutes in steps of 2.5 minutes; Supplementary Figure 23, Supplementary Text). We found that the within-subject NMI was higher than between-subject NMI with as little as 2.5 minutes of low-motion data *t*(2399) = 14.30, *p* < 0.0001 (Supplementary Table 1), and the two diverged by 7.5 minutes based on confidence interval (Supplementary Figure 23E, Supplementary Text). Taken together, we found that relatively stable individual-specific functional networks can be observed within a scan session and also across longitudinal sessions in children under 5 years old.

### Population consensus of individual-specific network assignments demonstrated high variability in the lateral frontal cortex, temporal-parietal junction, and the occipital-temporal cortex

Population consensus (i.e. the most frequent network at each vertex across the population) across all available subjects at Y2 (N = 113, Figure 4A) and at Y3 (N = 86, Figure 4B) of Dataset 1 appears to be similar to each other. Consistent with previous results from adults (Gordon et al., 2017b; Gratton et al., 2018; Kong et al., 2019; Li et al., 2019; Mueller et al., 2013), we found low variability at unimodal processing regions, such as the auditory, visual, and somatosensory and somatomotor cortices, and high variability at the lateral frontal cortex, temporal-parietal junction (TPJ), and the occipital-temporal cortex based on the percentage of subjects with the most frequent assignment (Figure 4C-D). Since this percentage only considered the most frequent assignment but not the frequency of other assignments, we also calculated a complementary “versatility” (Shinn et al., 2017) measure. Vertices that switch assignment among multiple networks across individuals would have a higher versatility than vertices that switch between two networks, even when the most frequent affiliation is the same.

**Figure 4.**
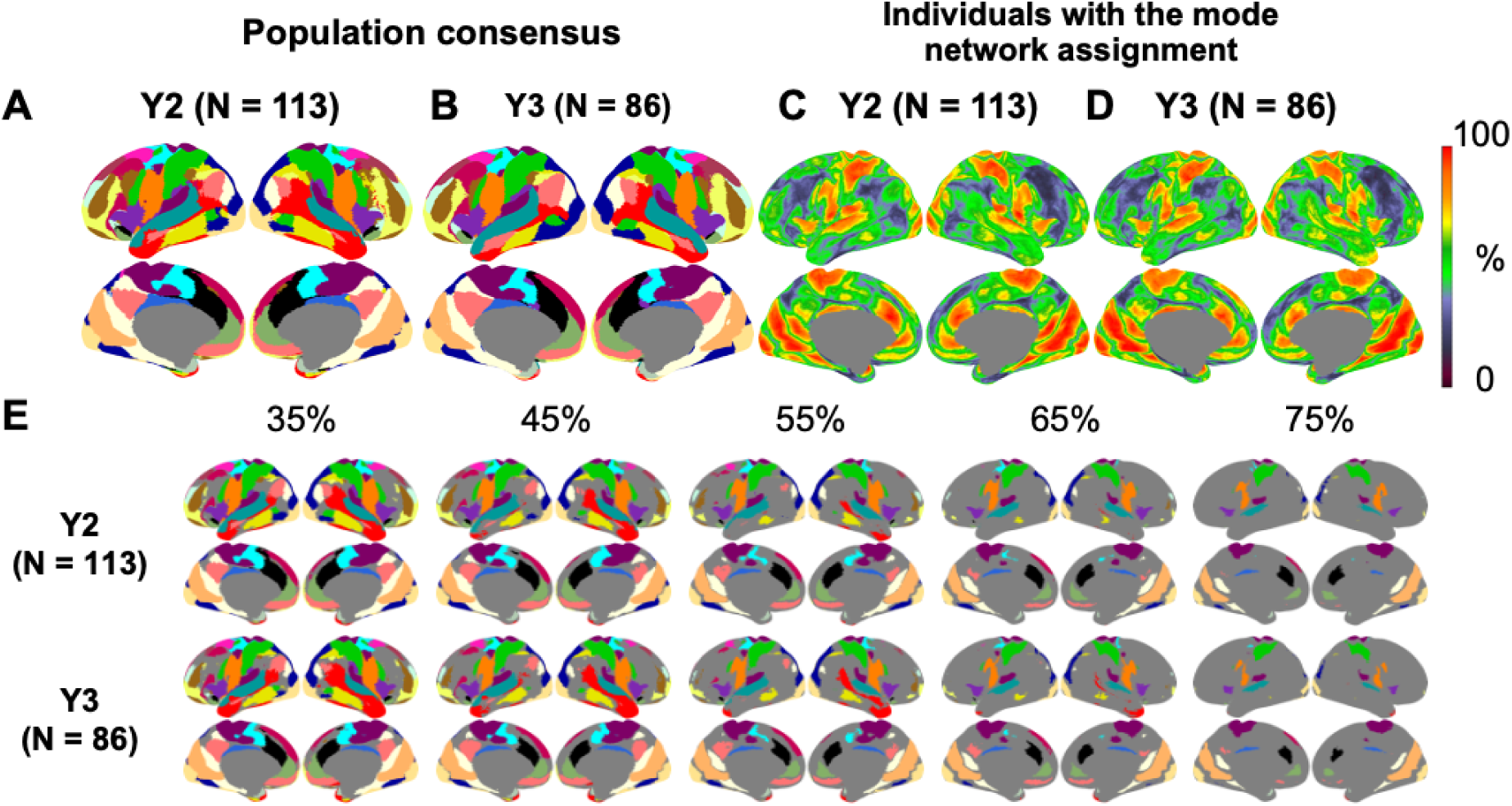
Population consensus network patterns (most frequent assignment) across subjects (Dataset 1). A) Population consensus (Y2, 113 subjects). B) Population consensus (Y3, 86 subjects). C-D) Percentage of subjects with the same assignment as the most frequent assignment. E) Population consensus of network identity for only the locations with consistent assignment among at least 35%, 45%, 55%, 65% and 75% subjects.

The resultant versatility spatial distribution (Supplementary Figure 24) is qualitatively similar to Figure 4C-D. To facilitate comparison to the spatial distribution of the most consistent network assignment reported in older 9 to 10-year-old children (Hermosillo et al., 2024), we also applied a threshold of at least 35%, 45%, 55%, 65% and 75% of individuals on the population consensus network assignments in Dataset 1 (Figure 4E). The locations with the least agreement among participants aligned with the boundaries between network patches, consistent with previous findings in adults where differences in borders among individual participants contributed to the majority of network variability (Dworetsky et al., 2024; Gordon et al., 2017b).

### Within-network connectivity is higher and reflects more age-related variance for individualized network topography compared to the group consensus network topography

So far, we have demonstrated the individual-specificity of the individualized network and the locations of variability. It remains unclear whether the individualized networks were superior to the group-level networks in capturing truly meaningful biologically relevant signals in individuals. If a network is correctly delineated, regions assigned to it should be more strongly connected to each other. Therefore, we compared the within-network FC for networks defined by individualized network topography or the group consensus network topography (from the whole dataset to avoid leakage of age, Y2 and Y3, N = 199, Figure 5A). We found that within-network FC was significantly higher for the individualized network topography compared with the group consensus network topography for 18 out of 23 networks (FDR-corrected p < 0.05, paired t-test). Dorsal Attention, Visual Central and Default-Parietal networks did not demonstrate a significant difference, while Visual Peripheral and Default-Pregenual had a significant lower within-network FC for individualized network topography compared with the group consensus network topography (FDR-corrected p < 0.05). The Visual Peripheral network result is potentially due to the group consensus network capturing the most strongly connected region within the network, which was highly consistent across individuals, but individualized networks sometimes include weaker FC at the boundaries.

**Figure 5.**
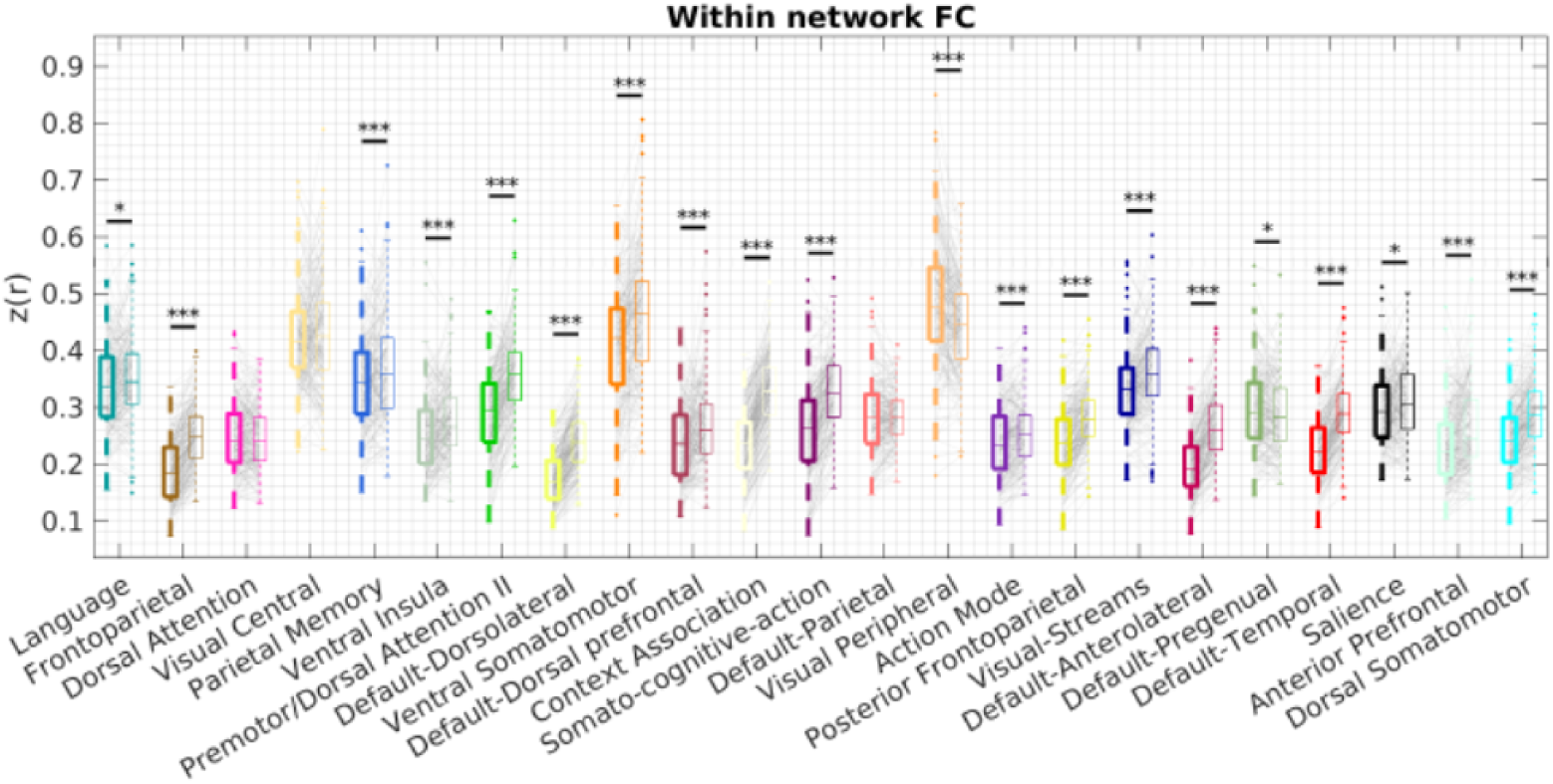
Within-network FC using network topography defined by group consensus or individualized network. A) Within-network FC with network topography defined by group consensus (bolded, left) or individualized network (right). * p<0.05, ** p<0.01, *** p<0.001. FDR-corrected.

To assess the developmental information captured by within-network FC beyond motion, we compared models including mean framewise displacement (FD) alone to models additionally incorporating within-network connectivity features. Mean FD alone explained negligible variance in age (R² ≈ 0). Adding the 23 within-network FC measures defined by individualized topographies explained substantially more age-related variance than adding the 23 within-network FC measures defined by group-consensus topographies (ΔR² = 0.35 vs. 0.21; adjusted R² = 0.26 vs. 0.11), despite identical model complexity. To assess subject-level generalization, we performed 5-fold cross-validation grouped by subject using ridge regression, ensuring the different sessions of the same subject fall in the same fold, with mean FD and within-network FC from all 23 networks as predictors. Prediction accuracy was evaluated by aggregating out-of-sample predictions across folds and computing the correlation between predicted and true age. Models using individualized network topographies achieved substantially higher prediction accuracy (r = 0.435, R² = 0.190) than models using group-consensus network topographies (r = 0.276, R² = 0.076), indicating that individualized network representations capture developmentally relevant information beyond group-level network definitions.

### Developmental variability in hemispheric lateralization is reflected in the individual-specific functional network topography

Given findings from prior literature about network lateralization, especially in the language networks, we sought to test the hypothesis that language-related networks increase in lateralization in young children and that the lateralization relates to the individual child’s language ability. We focused on two networks that contain regions commonly reported to be involved in language functions (Braga et al., 2020; Lipkin et al., 2022; Zhu et al., 2025): the language network (Network 1) and the default-anterolateral network (Network 18).

We started with Dataset 2, which is mixed cross-sectional with more subjects and sessions and covered a relatively large developmental window. Across children scanned at 8 to 60 months (Dataset 2), the language network primarily includes the superior temporal gyrus (STG), which supports auditory language processing. However, it sometimes incorporates the inferior frontal gyrus (IFG) and middle frontal gyrus (MFG), key components of the adult language network (Braga et al., 2020; Lipkin et al., 2022) that additionally contribute to domain-general processing (Fedorenko and Blank, 2020; Koyama et al., 2017) (Figure 6A, red arrows). Most frequently, the IFG and MFG were assigned to the default-anterolateral network (Figure 6B, red arrows). Compared to participants aged 8–10 months, children at 30–36 months had higher probability of left-hemisphere involvement and lower probability in the right hemisphere involvement for both networks, resulting in increased network lateralization across age (Figure 6C-H).

**Figure 6.**
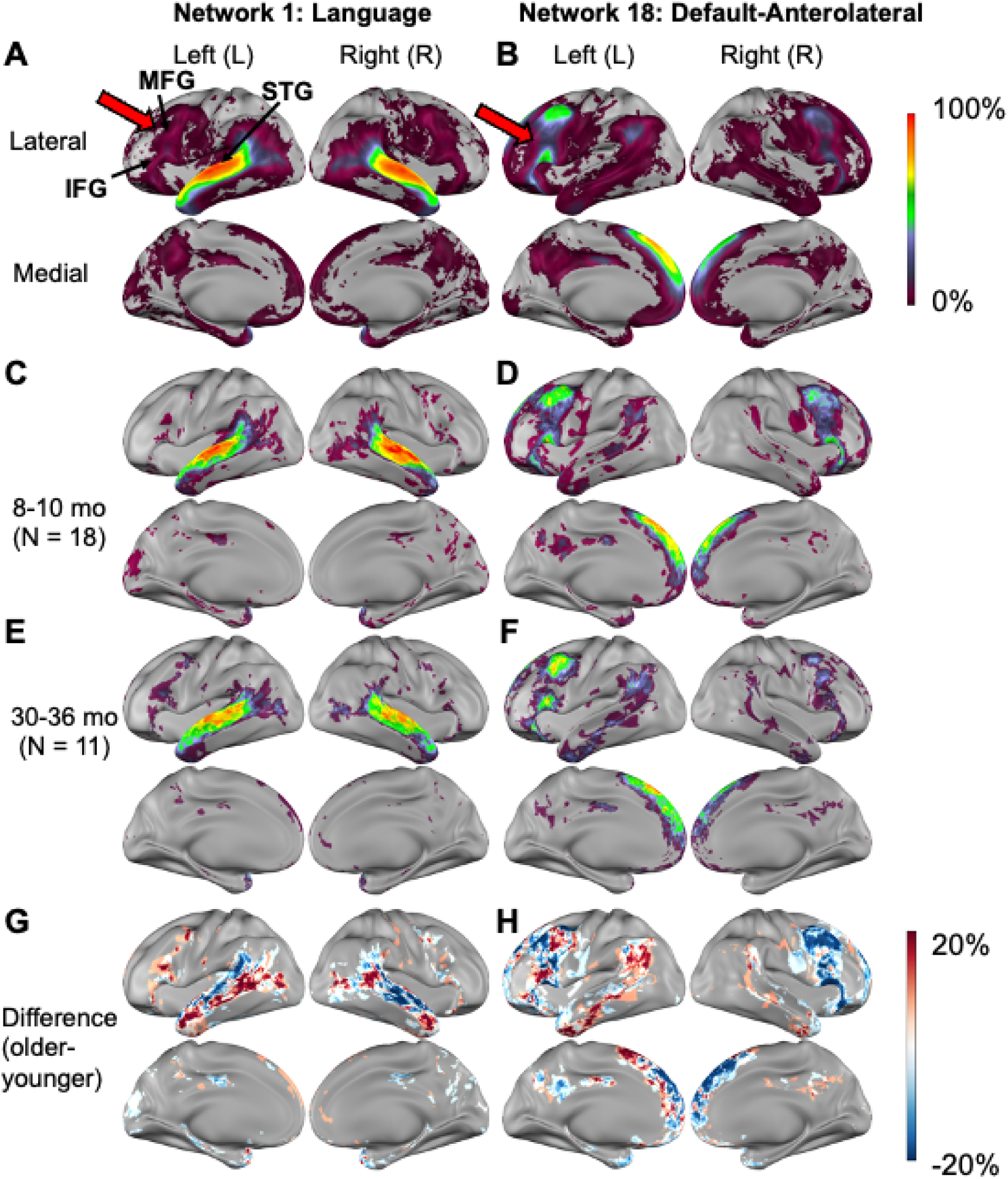
Spatial probability of Language and Default-Anterolateral networks. A) Language network probability and B) Default-Anterolateral network probability in children at 8-60 months (Dataset 2). C) Language network probability and D) Default-Anterolateral network probability at 8-10 months. E) Language network probability and F) Default-Anterolateral network probability at 30-36 months. G-H) Difference between the older and younger groups. MFG: Middle Frontal Gyrus, IFG: Inferior Frontal Gyrus, STG: Superior Temporal Gyrus.

To quantify network lateralization, we calculated a “laterality index” (LI) defined as the difference in the network surface area between the left and right hemispheres, expressed as a percentage of the total network surface area. In children at 8–60 months (Dataset 2), the Language network showed weak left-lateralization (LI = 0.07 ± 0.13; one-sample t-test, t (300) = 9.2858, p < 0.001), whereas the Default-Anterolateral network showed strong left-lateralization (LI = 0.40 ± 0.19; t (300) = 37.1876, p < 0.001). Using linear mixed-effects models with a random intercept for subject, laterality of the Language network increased with age (β = 0.025 per year, SE = 0.011, t = 2.29, df = 299, p = 0.023, Figure 7A). Laterality of the Default-Anterolateral network also increased with age (β = 0.059 per year, SE = 0.015, t = 3.82, df = 299, p < 0.001, Figure 7B). These associations were robust to the addition of mean framewise displacement of the scan session.

**Figure 7.**
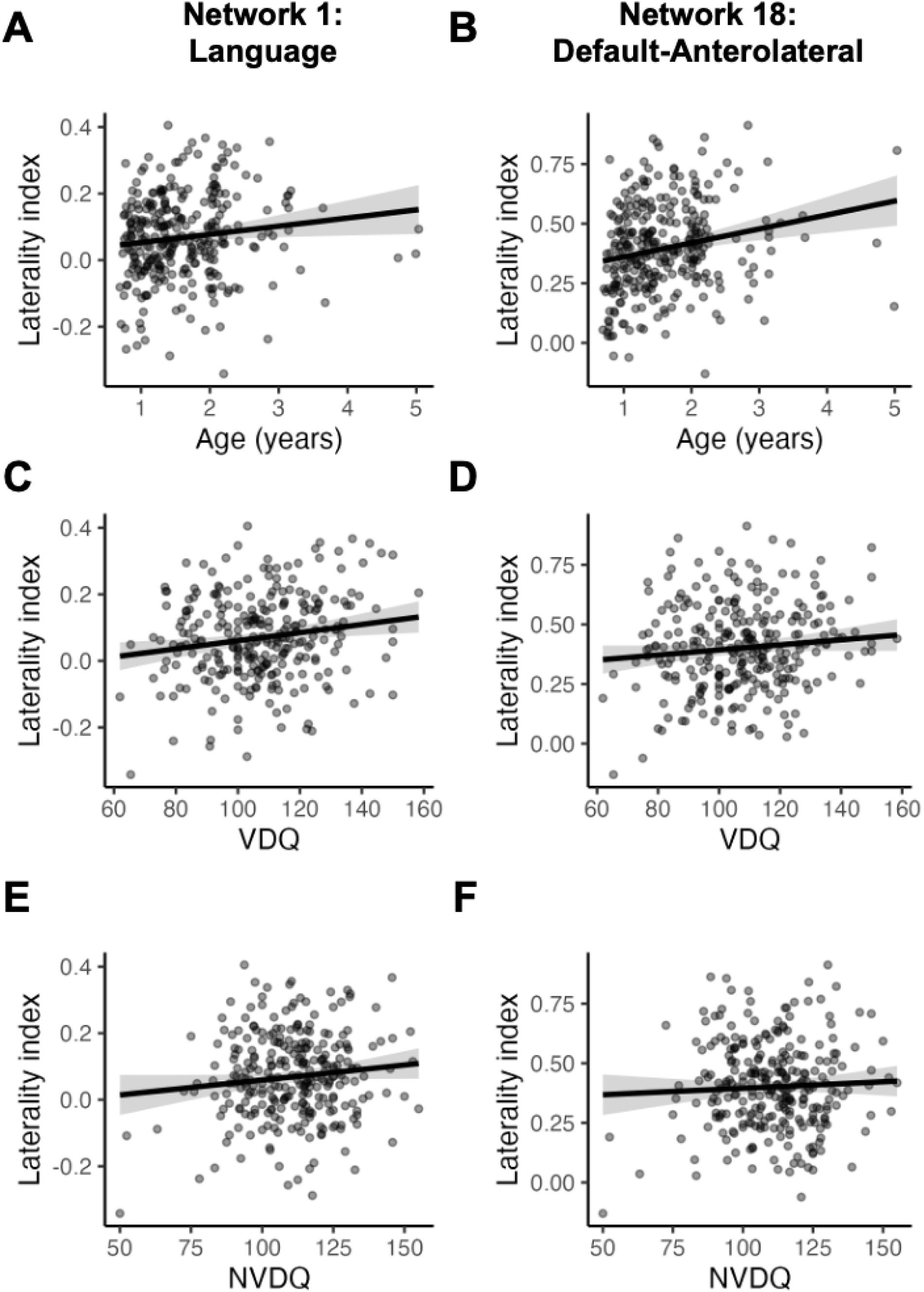
The relationship between functional network laterality and age or Mullen Scales of Early Learning developmental quotients. Laterality index against A-B) age, C-D) verbal developmental quotient (VDQ), missing 21 scores, E-F) non-verbal developmental quotient (NVDQ), missing 19 scores. Shaded band = 95% CI of fixed effect (age was held constant in C-F).

Associations between network laterality and behavioral measures were assessed using linear mixed-effects models including age and behavioral scores as fixed effects and a random intercept for subject. Verbal and non-verbal abilities were quantified using the Mullen Verbal Developmental Quotient and Non-verbal Developmental Quotient (VDQ and NVDQ; age-normalized scores from the Mullen Scales of Early Learning (Mullen, 1995)), excluding sessions without available scores (21 without VDQ and 19 without NVDQ). Language network laterality was positively associated with verbal ability beyond the effect of age (β = 0.00122 per VDQ point, SE = 0.00044, t = 2.79, df = 271.6, p = 0.0057, Figure 7C). In contrast, laterality of the Default-Anterolateral network did not significantly associate with VDQ (β = 0.00107 per VDQ point, SE = 0.00062, t = 1.72, df = 269.1, p = 0.087, Figure 7D). As a control analysis, Language network laterality did not significantly associate with NVDQ (β = 0.00090 per NVDQ point, SE = 0.00049, t = 1.85, df = 254.0, p = 0.066, Figure 7E), nor did the Default-Anterolateral network (β = 0.00055 per NVDQ point, SE = 0.00069, t = 0.79, df = 247.2, p = 0.430, Figure 7F).

Associations in the same direction were observed in the longitudinal dataset (Dataset 1), but statistical significance was reduced potentially due to a smaller sample size, a more restricted age range centered around 2–3 years, and the use of the Bayley Scales of Infant and Toddler Development language composite as a proxy for verbal ability (Supplementary Materials, Supplementary Figure 26).

## Discussion

### A fine network division for developmental detail

Many existing studies have demonstrated residual differences in functional network organization across adult individuals despite anatomical alignment (Bijsterbosch et al., 2023; Braga and Buckner, 2017; Dworetsky et al., 2021; Gordon et al., 2017c, 2017b; Gratton et al., 2018). With young children, there may be even more variability in functional template localization given the substantial amount of developmental changes in the brain during this period (Bethlehem et al., 2022). Smaller and more variable networks risk being obscured by the averaging of anatomically matched locations, and templates created from averaging within a generic network mask will be contaminated by signals from neighboring networks in individual subjects.

By design, individual networks generated from template matching and their population consensus resemble the network priors used in the creation of templates. Although some studies have tried to derive infant-specific network templates (Moore et al., 2024), they were still using a coarse adult network atlas to derive the templates. Moreover, a considerable amount of evidence suggests that there are finer network divisions in individuals than are present in many commonly used adult network atlases (Badke D’Andrea et al., 2025; Braga and Buckner, 2017; Gordon et al., 2023, 2020; Gratton et al., 2022; Lynch et al., 2024). Additional network detail may help capture subtle changes in development.

Here, we created templates directly on concatenated data instead of group-average data. By avoiding averaging FC at matching anatomical locations across individuals, we uncovered functional network patterns that are more representative of the individual FC profiles assigned to that network. These templates also produced better separation of neighboring networks. Our findings improved upon the current standard practice of averaging FC data across individuals even when the goal is to find group-level functional networks. For example, the population consensus division of the anterior temporal lobe reported here is more consistent with previous observations in adults (Akiki and Abdallah, 2019; Andrews-Hanna et al., 2010; Braga and Buckner, 2017; Du et al., 2024; Gordon et al., 2017c, 2016; Ji et al., 2019; Yeo et al., 2011), compared to the divisions identified directly in a group-average FC (Tu et al., 2025a). Even though we did not have secondary data like task fMRI and DTI to directly verify the biological validity of our functional network divisions, we did observe alignment between our networks and biologically relevant areas that increased activation in some task-based experiments in adults (Table 1).

### Precision functional mapping with a moderate scan length for scalability

Approximately 10 minutes of data can yield a moderate reliability of NMI = 0.46 in individual networks created from longitudinal sessions at 2 and 3 years old, but a longer scan than 10 minutes further increases the reliability of individual-specific networks up to and likely beyond 20 minutes (Supplementary Figure 23). Our results suggest that scan time per individual and data quality persist as a challenge in precision brain mapping (Du et al., 2024; Lynch et al., 2024), echoing claims from one recent study advocating for longer scan times per participant (Ooi et al., 2025). We reason that while high-quality data samples with many hours of fMRI data can be useful for data-driven exploration of individual-specific networks (Gordon et al., 2017c; Kwon et al., 2025; Laumann et al., 2015; Lynch et al., 2024) and within-subject study designs (Huth et al., 2016; Newbold et al., 2020), datasets with more moderate amount of data (e.g. 30 minutes to an hour, following recommendations in Ooi et al., 2025) per participant in relatively large number of participants may be used to increase the quality of group prior-based mapping of individual-specific networks. In line with this idea, recent papers found that a more extensive data collection is required to reach FC reliability in pediatric samples compared to adults, with diminishing returns after 40–50 minutes of post-censoring data (Moser et al., 2025; Rai et al., 2025). Since precise characterization of individual functional networks has been demonstrated to have clinical significance such as improving the efficacy of targeted brain stimulation (Cash et al., 2021; Lynch et al., 2022; Nahas et al., 2025), fast precision functional mapping with a moderate scan length using a network template offers a scalable solution for unique populations, broadening its potential applications. Further validation is required to assess the utility and data required of prior-based network mapping in those applications.

### Individual-specific functional network topography bridges brain specialization with behavioral development

We observed a link between functional network lateralization in the language network and language behavioral performance, beyond the effect of age. In contrast to the strong association between Language network lateralization and the verbal developmental quotient, the association for the Default-Anterolateral network is relatively weaker and statistically insignificant. This finding is consistent with prior studies in school-age children. In particular, greater resting-state laterality in the temporal gyrus but not the inferior frontal gyrus, has been positively correlated with linguistic skills (Day et al., 2024). This observation might be explained by the additional domain-general processing function of the IFG and MFG (Default-Anterolateral network) beyond language processing (Fedorenko and Blank, 2020; Koyama et al., 2017). While these findings alluded to the behavioral significance of the individual-specific functional networks and their potential utility in clinical applications in young children, such as presurgical or brain stimulation site planning, further validation is required to assess the amount of data needed for a reliable application of the individual-specific networks. Additionally, task-based activation is required to confirm the correspondence between the resting-state network topography and the functional co-localization.

### Functional network assignments are more consistent at sensory cortices than association cortices

Consistent with prior results in infants/toddlers (Gao et al., 2014; Molloy and Saygin, 2022; Xu et al., 2019), school-age children (Cui et al., 2020; Hermosillo et al., 2024), and adults (Du et al., 2024; Dworetsky et al., 2021; Gordon et al., 2017b; Gratton et al., 2018; Kong et al., 2019; Langs et al., 2016; Laumann et al., 2015; Li et al., 2019; Miranda-Dominguez et al., 2014; Mueller et al., 2013), we observed higher inter-individual variability of functional network topography in the association cortices than the sensory cortices in young children (Figure 2). As described in prior works (Hill et al., 2010; Tu et al., 2025b), this spatial bias in inter-individual variability may have a developmental origin: early maturation in some areas limits future plasticity. Additionally, the early-developing sensorimotor areas may be more evolutionarily primitive (Hill et al., 2010) and therefore genetically determined to be more consistent across human individuals. One recent paper examining the cell-type abundance in resting-state network divisions in post-mortem brain tissues seemed to support both ideas: the authors observed that sensorimotor network affiliations were better predicted by cell-type than association network affiliations, suggesting more stereotypical cell-type profiles in the sensorimotor networks (Zhang et al., 2024). The contribution of each factor to this observation is awaiting future exploration.

### Limitations and future directions

One large difference between extant neuroimaging data in young children and other populations is that most young children were scanned during natural sleep. A large amount of literature illustrates the differences in FC during sleep and those estimated during resting wakefulness (Horovitz et al., 2009; Larson-Prior et al., 2011, 2009; Mitra et al., 2015; Tagliazucchi et al., 2013), and even across different sleep stages (Tagliazucchi et al., 2012; Yang et al., 2024). Babies sleep much longer than adults, with up to 16 hours of sleep per day in neonates and 12 hours in 2-to-3-year-olds. Additionally, the sleep pattern is also different in neonates under 2 months of age, with up to 50% of sleep time spent in the REM/active sleep stage, compared to around 20% in adults (Roffwarg et al., 1966). Therefore, some of the developmental differences described here are potentially attributable to differences in the sleep-stage composition of fMRI data across different developmental stages.

While our choice of clustering on FC profiles with area parcels as seeds increased the computational efficiency substantially, it may have averaged out some fine spatial detail, limiting the number of functional networks that can be reliably identified across the population. Additionally, more extensive validation with task fMRI (Braga et al., 2020; Du et al., 2024; Nielsen et al., 2023) and behavioral phenotype prediction (Cui et al., 2020; Kong et al., 2019; Lynch et al., 2024) in larger cohorts with richer clinical representation will be important for further assessing the behavioral and clinical relevance of the proposed individual-specific functional networks. Such analyses may help determine whether individualized network representations can reliably distinguish clinical populations or identify early biomarkers of atypical brain development. Direct evidence from diffusion tensor imaging and cytoarchitecture would also be useful in validating the biological relevance of the identified networks. While we assign a single network label to each vertex for simplicity, network boundaries tend to exhibit more ambiguous assignments, and others have described this organization in terms of overlapping networks (Bijsterbosch et al., 2023).

Even though we only assigned cortical vertices to functional networks in the current study, the conceptual template matching framework is transferable to subcortical network assignments (Ji et al., 2019; Seitzman et al., 2020). Moreover, functional profiles based on task/movie responses (Haxby et al., 2020; Meschke et al., 2023; Zhi et al., 2022) could be utilized to further increase the reliability of the mapped network.

## Materials and Methods

### Datasets

We used two high-quality infant/toddler datasets spanning approximately 1-5 years (total number of participants = 328) sampling distinct socioeconomic backgrounds for the current study to ensure robustness of the findings. An overview is provided in Table 2, and details of the demographics, acquisition, and processing of the data were reported in other studies with slight differences in the selection of final samples to include (Tooley et al., 2024; Tu et al., 2025a) and are briefly recapitulated below. Age and sex distributions are provided in Supplementary Figures 2 and 21.

**Table 2.**
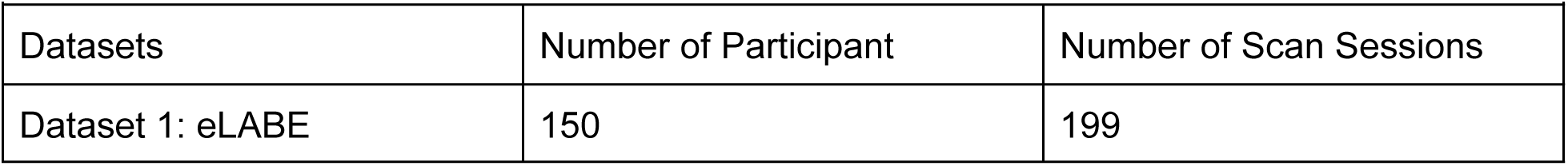
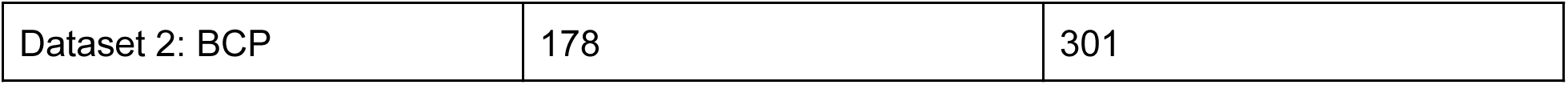
Dataset summary.

### Dataset 1: Early Life Adversity, Biological Embedding (eLABE)

#### Demographics

We used the Y2 (approximately 2 years old) and Y3 time (approximately 3 years old) point data from the eLABE study with a longitudinal data collection design as our discovery dataset, with basic demographic information summarized in Supplementary Figure 2. This study was approved by the Human Studies Committees at Washington University in St. Louis, and informed consent was obtained from a parent of all participants. Recruitment oversampled mother-infant pairs facing adversity (e.g., poverty and stress). At each time point, all participants with at least 8 minutes of usable low-motion fMRI data and no major quality issue in the structural or functional MRI were included, resulting in a total of 150 participants and 199 sessions, of which 49 individuals had both Y2 and Y3 sessions. A subset of 81 sessions (40 from Y2 and 41 from Y3) from healthy term-born toddlers had more than 20 min of usable low-motion data were used to generate the network templates, and vertexwise assignments of individualized networks were created for all sessions. Since we wanted to sample a diverse demographic for our results, and for increasing statistical power, we did not exclude the 20 preterm-born subjects.

#### MRI data acquisition

MRI data were acquired during natural sleep without sedating medications using a 3T Prisma scanner (Siemens Corp.) and 64-channel head coil loosely following HCP-style acquisition parameters. During each scan session, a T1-weighted image was collected (sagittal, 208 slices, 0.8-mm isotropic resolution, repetition time = 2400 ms, echo time = 2.22 ms). Functional imaging data (fMRI) were collected using a blood oxygen level–dependent (BOLD) gradient-recalled echo-planar multiband sequence (72 slices, 2.0-mm isotropic resolution, echo time = 37 ms, repetition time = 800 ms, multiband factor = 8, 420 volumes) for a total of 2-9 runs (5.6 minutes each). Spin-echo field maps (at least 1 anterior–posterior and 1 posterior-anterior) were obtained during each session with the same parameters. Framewise Integrated Real-time MRI Monitoring (FIRMM) software suite was used to monitor motion summary in realtime (Dosenbach et al., 2017).

#### MRI data preprocessing

Anatomical scan processing and segmentation were conducted using Freesurfer 7.2. Field distortion correction was performed using the FSL TOPUP toolbox. Functional data preprocessing followed established procedures (Power et al., 2014), which includes the basic correction of intensity differences attributable to interleaved acquisition, bias field correction, intensity normalization of each run to whole-brain mode value of 1000, linear realignment within and across runs to compensate for rigid body motion, and linear registration of BOLD images to the adult Talairach isotropic atlas performed in one single step that includes the BOLD to individual T1 to group-average T1 from this cohort to 711-2N Talairach atlas. Subsequently, volumetric resting-state BOLD time series were projected to fs_LR-32k aligned surfaces for each individual using spherical registration and smoothed with geodesic 2D Gaussian kernels (σ = 2.25 mm) (Van Essen et al., 2012). Additional fMRI preprocessing to mitigate the effect of motion and other non-neuronal sources of signal was conducted with (i) demean and detrend, (ii) nuisance regression including white matter, ventricles, extra-axial cerebrospinal fluid, and the whole brain mean signals and the 24-parameter Friston expansion regressors derived from the 6 rigid body realignment parameters (x, y, z, pitch, yaw, roll), (iii) bandpass filtering to 0.005-0.1 Hz (Power et al., 2014) after the BOLD data was mapped to surface. Framewise displacement (FD) was calculated to index the amount of motion within each session and for the creation of a motion censoring time mask. Specifically, the 6 rigid body realignment parameters were first filtered with an age-specific respiratory notch filter (0.27-0.5 Hz) to reduce the impact of artifactual motion introduced by B0 field perturbation (Fair, 2020; Kaplan et al., 2022). FD was then calculated as the sum of the absolute values (L1-norm) of the differentiated notch-filtered 6 rigid body realignment parameters (by backwards differences) at every time point, with the FD of the first frame set to 0 by convention. fMRI data were then censored based on a threshold of FD_filt_ > 0.2 mm, with the additional restriction that only epochs of at least 3 consecutive frames FD_filt_ < 0.2 mm were included, and the first 5 frames were also excluded. Lastly, due to imperfect registration of the medial wall, a small subset of vertices were missing data at the hippocampal regions, and were imputed by the mean value of the nearest neighbor using a custom MATLAB script.

### Dataset 2: Baby Connectome Project (BCP)

#### Demographics

Full-term (gestational age of 37-42 weeks) infants free of any major pregnancy and delivery complications were recruited as part of the Baby Connectome Project (Howell et al., 2019). All procedures were approved by the University of North Carolina at Chapel Hill and the University of Minnesota Institutional Review Boards. Informed consent was obtained from the parents of all participants. In the final cohort used following fMRI data quality control, we retained 301 sleeping fMRI sessions from 178 individuals acquired during natural sleep.

#### MRI data acquisition

All MRI images were acquired on a Siemens 3T Prisma scanner with a 32-channel head coil at the University of Minnesota and the University of North Carolina at Chapel Hill during natural sleep without the use of sedating medications. T1-weighted (TR=2400 ms, TE=2.24 ms, 0.8 mm isotropic; flip angle = 8°), T2-weighted images (TR=3200 ms, TE=564 ms, 0.8 mm isotropic), spin echo field maps (SEFM) (TR=8000 ms, TE=66 ms, 2 mm isotropic, MB=1), and fMRI data (TR=800 ms, TE=37 ms, 2 mm isotropic, MB=8) were collected. A mixture of Anterior→Posterior (AP) and Posterior→Anterior (PA) phase encoding directions was used for fMRI acquisition in each session, but they were concatenated into one time series. A subset of data (88 sleeping sessions) had a 720-ms TR instead of an 800-ms TR.

#### MRI data preprocessing

Data processing used the DCAN-Labs infant-abcd-bids-pipeline (v0.0.22), which largely follows the HCP processing (Glasser et al., 2013) and the steps described previously for the ABCD dataset (Feczko et al., 2021). The processing steps were similar to the adult data except for the use of infant-specific MNI templates at ages 08-11/11-14/14-17/17-21/21-27/27-33/33-44/44-60 months (Fonov et al., 2009) to better register the structural data. In addition, segmentation of the brain structures was conducted with Joint Label Fusion (JLF). The toddler-specific brain mask and segmentation were substituted for the Freesurfer (Fischl, 2012) pipeline to refine the white matter segmentation and guide the FreeSurfer surface delineation for each individual scan session of each participant. The native surface data were then deformed to the fs_LR-32k template via a spherical registration.

Functional data processing was largely the same as before, except for the following minor differences: (i) across-run intensity normalization to a whole-brain mode value of 10,000 instead of 1000. (ii) An infant MNI template was used for registration. (iii) fMRI BOLD volumes were sampled to native surfaces using a ribbon-constrained sampling procedure available in Connectome Workbench and then deformed and resampled from the individual’s “native” surface to the fs_LR-32k surface with minimal smoothing at σ = 0.85 mm. Additional steps to mitigate the non-neuronal sources of artifact was conducted on the surface data similar to Dataset 1 with a few minor differences: (i) the mean signal of extra-axial cerebrospinal fluid was not included in the regression, and the mean signals of the 91k grayordinates substituted the whole-brain signal in the nuisance regression, the first-order derivative of the white matter, ventricular CSF and been grayordinate signals were also included for nuisance regression (ii) a respiratory notch filter (0.28-0.48 Hz) (Fair, 2020; Kaplan et al., 2022) was applied to the motion parameters estimates before both FD calculation and the construction of the 24-parameter Friston expansion regressor. (iii) fMRI data were then censored based on a threshold of FD_filt_ > 0.2 mm, and the first 7 frames were also excluded. Subsequently, outlier frames whose across-grayordinate standard deviation was more than 3 median absolute deviations from the median of all frames included, and individual runs were further excluded from a very small subset of sessions due to failure of the visual quality check. A further smoothing with a geodesic 2D Gaussian kernel (σ = 2.40 mm) was applied to give a final effective smoothing of σ = 2.55 mm. We then focused on only the cortical vertices on the fs_LR-32k surface (59412 after excluding medial masks) for the subsequent analyses.

### Functional connectivity representations

Analyses were conducted using an age-specific cortical parcellation previously demonstrated to fit this developmental period (Tu et al., 2025a). Parcels refer to anatomically defined cortical regions from this parcellation. Vertices refer to surface mesh points on the cortical surface. Two types of functional connectivity (FC) representations were used, each serving a distinct analytical role. Parcel-to-parcel FC was used to identify reproducible connectivity patterns across individuals. Vertex-to-parcel FC was used to assign network identities to individual cortical vertices based on data-driven network templates. In both cases, FC was defined as the Pearson correlation between BOLD time series and computed across the neocortex in both hemispheres.

### Identification of age-appropriate functional connectivity patterns

For each fMRI session, a parcel-by-parcel FC matrix was computed. Each parcel’s connectivity profile to all other parcels (i.e., one row of the parcel-to-parcel FC matrix) was treated as a single parcel-wise FC profile. In this representation, parcels serve both as samples (rows) and as features (connectivity targets). Parcel-wise FC profiles from all sessions were concatenated across individuals prior to clustering. The analysis included 81 fMRI sessions from 72 unique children aged 2–3 years (9 children contributed longitudinal data at Y2 and Y3; all sessions contained >20 minutes of low-motion data; eLABE Dataset 1). Clustering was performed using K-means with correlation distance. The resulting cluster centroids represent reproducible parcel-wise FC patterns and were used as age-appropriate reference network profiles. Initial centroid position was determined by the K-means++ algorithm (default). Because the K-means solution is susceptible to the initial centroid location, we set the number of times to repeat the clustering to 1000 to minimize the possibility of stopping at a local minimum. The output K cluster centroids were subsequently used as network templates for template matching.

### Selection of the number of clusters

The number of clusters (K) was selected using a split-half stability analysis. The 81 sessions were randomly divided into two halves 20 times. For each split, parcel-wise FC profiles were concatenated separately and clustered using k-means with K ranging from 1 to 40. Cluster stability was quantified as the normalized mutual information (NMI) between cluster assignments predicted across split halves. A local maximum in stability was observed at K = 23, which was selected for all subsequent analyses. This choice favors sensitivity to finer-grained connectivity patterns, including patterns that may be consistently expressed in only a subset of individuals.

### Individual-specific network mapping

To derive individual-specific network organization at finer spatial resolution, reference network profiles were mapped onto the cortical surface at the vertex level. For each session, vertex-to-parcel FC was computed by correlating the BOLD time series of each cortical vertex with the mean BOLD time series of each parcel, yielding a vertex-wise FC profile for every vertex. Each vertex-wise FC profile was compared with the parcel-level reference network profiles using Pearson’s correlation. Vertices were assigned to the network with the highest correlation using a winner-take-all procedure, similar to prior studies (Gordon et al., 2017a). No magnitude thresholding or distance-based exclusion was applied to the FC profiles. We retained all connections, including weak and anticorrelated values, as these may contribute to network differentiation, particularly in early development.

### Reliability of individual-specific functional networks

One way to measure reliability is to compare the individual-specific functional networks generated using different data samples from the same session. We split the number of low-motion frames into the first half and second half of the session and calculated two functional connectomes. Then, we generated the individual-specific functional networks from the two functional connectomes and compared them using the normalized mutual information (NMI). Similarly, we generated the individual-specific functional networks from the two functional connectomes generated using the fMRI data in Y2 and Y3 time points for the same individual to examine longitudinal reliability.

### Nodal versatility

Nodal versatility (V) (Shinn et al., 2017) measures whether a node is consistently assigned (V ≈ 0) to a specific community, or it is inconsistently assigned to different communities (V >> 0) across different partitions (in our case, different individual networks).

### Lateralization of individual-specific functional networks

Since the fsLR-32k surface template does not sample the gyral and sulcal surfaces equally, to get an accurate measure of the individual-specific functional network size, we calculated the surface area of each vertex using the Connectome Workbench (version 1.2.3) command “wb_command -surface-vertex-areas” using the participant’s midthickness surface obtained from the structural data in the same scan session. The sizes of a given network in each of the left and right hemispheres were calculated as the sum of the surface area of all vertices assigned to that network in the individual in the corresponding hemisphere. A lateralization index was then calculated as the difference in network sizes in the left hemisphere and right hemisphere, divided by the total network size.

### Behavioral measures: Mullen Scales of Early Learning (MSEL)

Mullen Scales of Early Learning (Mullen, 1995) was a standardized assessment of language, motor, and perceptual abilities for children through 5 years of age. There are 5 subdomains: 1) gross motor (GM), 2) fine motor (FM), 3) visual reception (VR), 4) receptive language (RL), and 5) expressive language (EL). The assessment takes 20-45 minutes and was implemented at every behavioral visit between 3 and 60 months of age (Howell et al., 2019). We calculated verbal and non-verbal developmental quotient scores (VDQ and NVDQ) as defined by the formula below with the age-equivalent (AE) scores from the subdomains (Stephens et al., 2018):

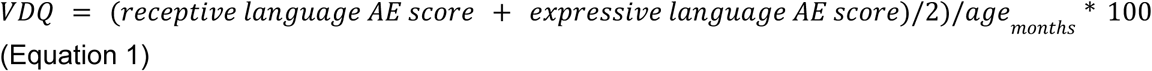

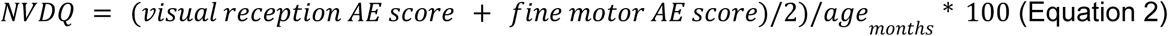

### Calculating similarity between networks with spatial probability and FC

To put our 23 functional networks in the context of prior literature and determine the hierarchy in the functional networks, we adapted a method from a prior paper to calculate a similarity measure using both the spatial overlap and FC (Lynch et al., 2024). Specifically, the overall similarity is calculated as the product of FC similarity and spatial overlap. In the original code base, binarized infomap community topography was used so the spatial overlap was calculated as the average spatial probability covered by that community. In our study, our network topography was specified as continuous (probability, ranging from 0 to 1) rather than binarized value, so the spatial overlap of each of our 23 networks and the Lynch network priors was modified to be calculated as the summed product of the spatial probability of that network and the Lynch network priors, divided by the sum of the spatial probability of that network, similar to the conditional probability formula. The final script “pfm_identify_networks_precalculated.m” is available at https://github.com/cindyhfls/PFM, adapted from https://github.com/cjl2007/PFM-Depression.

### Quality of separation of the FC profiles

To measure the quality of separations into clusters defined by the network assignments from the group-average FC and the K-means optimized assignments, we use the silhouette index (SI) metric (Rousseeuw, 1987) for each individual FC profile defined as:

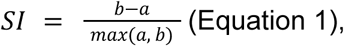

where a is the mean correlation distance (i.e. 1-correlation) to all other FC profiles within the assigned cluster, and b is the mean correlation distance to all FC profiles within the best alternative cluster. The SI ranges from +1 to −1 with the sign indicating whether the area has a more similar FC profile within the assigned cluster (+) or to FC profiles in an alternative cluster (-). The magnitude indicates the degree of separation between the clusters.

### Statistics and Reproducibility

Associations between network laterality and age or behavioral measures were assessed using linear mixed-effects models implemented in R (lme4), with fixed effects for predictors of interest and a random intercept for subjects to account for repeated scans. Age was included as a covariate in all models due to its developmental relevance; mean framewise displacement was included in secondary analyses to assess robustness to motion. Model diagnostics of “outliers” included inspection of standardized residuals and subject-level influence using Cook’s distance (Supplementary materials - “outlier analysis”); no outcome-based exclusions were applied. Analyses were hypothesis-driven and focused on predefined language-related networks and measures, and p-values are therefore reported without correction for multiple comparisons.

## Code Availability

The analysis code has been made publicly available to ensure reproducibility of the results: https://github.com/cindyhfls/Tu-2026-IndividualBabyNetworks.

## Data Availability

The fMRI and phenotypic data supporting this study are available from the corresponding datasets. Data from the Baby Connectome Project can be accessed via the NIH Data Archive (https://nda.nih.gov/edit_collection.html?id=2848). Data from the eLABE study are available upon request at https://eedp.wustl.edu/research/elabe-study/

## Acknowledgments

The authors thank Dr. Julia Moser for their helpful discussions on the template matching procedures, and Dr. Matthew Glasser for suggestions on network nomenclature and Dr. Samuel R. Krimmel for providing the surface-to-volume transformation script. The current study is funded by the NIH (R00EB029343 and R01HD115540 to MDW; R01MH122389, R01MH134966, and R01MH131584 to CMS). The Early Life Adversity, Biological Embedding (eLABE) study was supported by NIH/NIMH R01 MH113883. The Baby Connectome Project was supported by NIH/NIMH R01 MH104324 and NIH/NIMH U01 MH110274.

## Author contribution

Conceptualization - JCT. Formal analysis/visualization - JCT and CL. Data curation - JCT, TKMD, LAM, XW, DD, AL, JKK. Methodology/software - EMG, RH, LAM, AW. Funding acquisition - MDW, JTE, DMB, BBW, JLL and CDS. Writing-original draft - JCT and MDW. Writing - review and editing of the final manuscript - all authors.

## Declaration of generative AI and AI-assisted technologies in the writing process

During the preparation of this work the author(s) used ChatGPT and Deepseek in order to help with writing style and to improve the clarity of the sentences. After using this tool/service, the author(s) reviewed and edited the content as needed and take(s) full responsibility for the content of the published article.

