## Supplemental Materials for "Commonality and Variability in Functional Networks in Children Under 5 Years Old"

### Supplementary Materials

#### Supplementary Text

##### Group-averaged FC may obscure finer individual-specific features present in individual participants' data

To illustrate this, consider each row of the parcel-by-parcel functional connectome as an FC profile—the connectivity pattern between one parcel and all others (Supplementary Figure 1A). These profiles can also be mapped onto a standard brain surface for spatial visualization (Supplementary Figure 1A). Averaging FC matrices from 81 template sessions (examples in Supplementary Figure 1A–B) yields a group-level FC (Supplementary Figure 1C). However, individual variability is evident in the distribution of pairwise correlations between FC profiles, such as lateral prefrontal cortex parcels across sessions (Supplementary Figure 1D). Some correlations are low, near or below zero (Supplementary Figure 1D, with an example pair in Supplementary Figure 1E–F), indicating weak consistency across individuals. Consequently, the group-averaged FC (Supplementary Figure 1C/G) may obscure finer individual-specific features in individual participants' data.

##### Replicating the split-half reliability analysis of individual-specific functional networks with no overlapping runs across split halves

For the split-half reliability analysis of individual-specific functional networks, we evenly divided the low-motion frames across all fMRI runs in a single scan session to maximize the amount of data per split half. However, due to temporal autocorrelations and shared preprocessing, having neighboring frames within a run in the two split halves could artificially inflate the NMI. Therefore, we repeated the same analysis but calculated the split-halves, ensuring no overlapping runs in the two split halves (e.g., the middle of 5 runs is dropped). The overall NMI is lower, potentially due to both less data per split half and reduced temporal dependence between split halves, but the conclusions remain the same (Supplementary Figure 20). The consistency of network assignment across the split halves (measured with normalized mutual information, NMI) was still significantly higher (two-sample t-test $p<0.001$) within individual (Y2: NMI = 0.4416 ± 0.0934, Y3: NMI = 0.4710 ± 0.0882) than between individuals (Y2: NMI = 0.3037 ± 0.0517, Y3: NMI = 0.3123 ± 0.0409).

##### Effect of data amount on longitudinal network stability

Because within-subject NMI is higher with more data, we conducted additional analysis to assess the effect of the data amount on longitudinal network stability in the 49 subjects with longitudinal sessions by sampling parts of the data from the whole session. Notably, due to the temporal autocorrelation in BOLD data, random sampling of the same number of fMRI volumes (TRs) within a long window tends to increase data reliability than contiguous data sampling (Hermosillo et al., 2024; Laumann et al., 2015). Therefore, the use of the term "minutes" to refer to random sampling can be misleading and not informative for data collection. We sampled contiguous data for the number of frames that approximately match the amount of target data length based on a TR of 0.8 seconds in the fMRI data to better simulate actual data collection.

For example, we sampled the first 375 frames of low-motion data to obtain a target length of 5 minutes. Because of the frame censoring procedure, the acquired data length is often slightly greater than 5 minutes and may be variable across subjects, but we omit that detail for simplicity. Since there is a sharp drop-off in available subjects at a threshold of 20 minutes' low-motion data (Supplementary Figure 23A), we sampled data segments with lengths of 2.5 to 20 minutes at a step of 2.5 minutes for both Y2 and Y3 data. The number of available subjects went down with longer data length, with only 11 subjects in the most sparse bin (20 minutes in Y2 and 20 minutes in Y3). When comparing the networks obtained from an increasing amount of data from 5 minutes to 20 minutes in Y3 to networks obtained from a fixed 20 minutes of data in Y2, both within and between network NMI increases (Supplementary Figure 23B), but within-subject NMI is always higher than between-subject NMI. We also calculated the mean within-subject (Supplementary Figure 23C) and between-subject (Supplementary Figure 23D) NMI for each data-length pair, and found that increasing the data amount in both Y2 and Y3 increased the within-subject and between-subject NMI, but the within-subject NMI is always higher than the respective between-subject NMI. The difference between within- and between-subject NMI is significant even when both Y2 and Y3 have 2.5 minutes of low-motion data $t(2399) = 14.30$, $p < 0.0001$ (Supplementary Table 1), and the 83.4% confidence interval started to have no overlap before 7.5 minutes. An 83.4% confidence interval was deemed more suitable for evaluating the difference between two means rather than comparing a mean to a fixed value, as this approach maintains a Type I error rate close to $\alpha = 0.05$ when the standard errors of the samples are comparable (Knol et al., 2011; Payton et al., 2003).

Supplementary Table 1. NMI between longitudinal sessions with different amounts of low-motion minutes.

| | | | | | |
|---|---|---|---|---|---|
| 2.5 | 0. 2415 ± 0.0475 | 0. 1790 ± 0.0208 | 2399 | 14.30 | <0.0001 |
| 5 | 0. 3511 ± 0.0467 | 0. 2604 ± 0.0308 | 2399 | 20.17 | <0.0001 |
| 7.5 | 0. 4145 ± 0.0425 | 0. 3054 ± 0.0297 | 2399 | 25.19 | <0.0001 |
| 10 | 0. 4564 ± 0.0392 | 0. 3339 ± 0.0299 | 2350 | 27.91 | <0.0001 |
| 12.5 | 0. 4839 ± 0.0369 | 0. 3519 ± 0.0306 | 2062 | 27.54 | <0.0001 |
| 15 | 0. 5038 ± 0.0376 | 0. 3643 ± 0.0316 | 1888 | 27.13 | <0.0001 |
| 17.5 | 0. 5250 ± 0.0379 | 0. 3744 ± 0.0333 | 804 | 18.94 | <0.0001 |
| 20 | 0. 5371 ± 0.0460 | 0. 3848 ± 0.0343 | 570 | 14.47 | <0.0001 |
| *Statistics from a two-sample t-test (two-tailed). | | | | | |

Finding the correspondence of networks to literature mentions

We used two major toolboxes to put our networks in context with prior literature mentions. Specifically, we assessed the spatial correspondence between our 23 functional networks and canonical networks from prior literature using the “Network Correspondence Toolbox” version 0.2.0 (Kong et al., 2025), and we reported the most frequently reported labels to our network topography using the “Neurosynth discrete decoding” in “NiMARE: Neuroimaging Meta-Analysis Research Environment” version 0.5.0 (Salo et al., 2023). Both are open-source Python-based software for neuroimaging meta-analyses.

For the analysis with the Network Correspondence Toolbox, nine representative atlases from different research groups were selected as references: “Du2024-15” (Du et al., 2024), “Woodward2024-Task-12”(Percival and Woodward, 2025), “Yan2023-400+Kong2019-17”(Kong et al., 2019; Yan et al., 2023), “Gordon2024-5"(Gordon et al., 2023), “Gordon2017-17”(Gordon et al., 2017), “Shen2013-268-8"(Shen et al., 2013) and “Glasser2016-360+Ji2019-12”(Glasser et al., 2016; Ji et al., 2019), “Tu2025-19”(Tu et al., 2025), “Kardan2022-10”(Kardan et al., 2022). All reference atlases were projected to the fs_LR-32k surface space to match the input data using the toolbox. A binarized map with network locations that are present with probability greater than 0.35 in the 81 template sessions was compared with the references using a Dice similarity coefficient. To determine statistical significance, we applied a spin test with 1000 surface-based rotations of each input map, generating a null distribution of Dice coefficients. The empirical p-value was defined as the proportion of rotated Dice values exceeding the observed Dice coefficient. Reference networks with $p < 0.05$ were considered to exhibit significant spatial similarity. The significantly overlapping networks in each atlas and the corresponding dice coefficient are visualized in circular bar chart plots (Supplementary Figures 5-16), and the raw dice coefficients and p-value statistics of all networks in all atlases are available in csv files at https://github.com/cindyhfls/Tu-2026-IndividualBabyNetworks/tree/main/Supplementary/NetworkCorrespondenceToolboxResults. For all but one atlas (a.k.a. “Woodward2024-Task-12”), the network acronyms are intuitive and have been used widely in the systems neuroscience literature. However, the task-based network acronyms are less commonly encountered, so we provide a key here for reference:

1RESP – Response One-Handed
2RESP – Response Two-Handed
AAR – Auditory Attention Response
AUD – Auditory Primary Sensory
DMNA – DMN Novel
DMNB – DMN Traditional
FoVF – Focus on Visual Features
INIT – Initiation
LN – Language
MAIN – Maintaining
MDN – Multiple Demand
RE-EV – Re-evaluation

For the analysis with the "NiMARE toolbox, we first created a binarized map with network locations that are present with probability > 0.35 and transformed it to the MNI space using the Connectome Workbench command "metric-to-volume-mapping" with the nearest vertex interpolation method and filled in the holes with "-volume-fill-holes". Subsequently, we employed the "ROIAssociationDecoder" tool to perform a region-based discrete decoding by correlating each binarized map with term-specific meta-analytic maps. These maps were derived from Neurosynth v7, a large-scale database of over 14,000 fMRI studies (Yarkoni et al., 2011). Following standard Neurosynth procedures, we used annotations based on a vocabulary of 3,229 curated cognitive and functional terms. These terms were selected from study abstracts using a Term Frequency–Inverse Document Frequency (TF-IDF) approach, which prioritizes words that are frequent in specific documents but uncommon across the full corpus. This selection process ensures that the vocabulary contains meaning and discriminative concepts in neuroimaging literature. For each of the 23 binarized maps, NiMARE computed the Pearson correlation coefficient with every term's meta-analytic activation map, which quantifies their spatial alignment. From the resulting associations, the 15 top-ranked terms per map were selected to represent the dominant cognitive functions associated with the network. To facilitate interpretation, we excluded general terms such as "network", "cortex", "area", "lobe", "region", "system", "activation", "brain", "functional", "related", "associated", "healthy", "task", "mode" and "connectivity". Morphological variants of the same word, such as past-tense verbs and plural nouns, were removed through lemmatization using the Natural Language Toolkit Python package (Bird and Loper, 2004) version 3.9.1. These selected terms were visualized as word clouds, where font size was scaled by the strength of the correlation to the meta-analytic activation maps (N.B. same scales were used for all networks), to highlight the most relevant labels for each network.

#### Lateralization of the functional networks in Dataset 1

The left lateralization of networks were replicated in Dataset 1 (longitudinal data at 22-47 months), both the Language network (LI = 0.13 ± 0.16, one-sample t-test, t (198) = 11.2807 , $p < 0.001$) and the Default Anterolateral network (LI = 0.44 ± 0.17, one-sample t-test, t (198) = 37.0012 , $p < 0.001$) were left lateralized. Laterality of the Language network increased significantly with age (β = 0.055 per year, SE = 0.016, t = 3.50, df = 91.9, p = 0.0007, Supplementary Figure 25A), but the laterality of the Default-Anterolateral network did not (β = 0.017 per year, SE = 0.020, t = 0.86, df = 140.6, p = 0.39, Supplementary Figure 25B). Verbal ability was measured with Bayley Scales of Infant and Toddler Development age-normalized language composite scores, which was calculated from 49 items in the receptive and 48 items in the expressive domain (Balasundaram and Avulakunta, 2025). There is a weak positive association between laterality of the Language network (β = 0.0016 per composite score point, SE = 0.0008, t = 0.86, df = 175.2, p = 0.052, Supplementary Figure 25C) and the Bayley language composite scores that did not reach statistical significance. Similarly, the association between the Default-Anterolateral network laterality and the Bayley language composite scores did not reach significance (β = 0.0012 per composite score point, SE = 0.0008, t = 0.86, df = 154.7, p = 0.144, Supplementary Figure 25D).

Outlier analysis

To assess whether associations between network laterality and age or behavioral measures were driven by extreme observations or influential participants, we conducted residual and influence diagnostics for all linear mixed-effects models. Specifically, we computed the proportion of observations with standardized residuals exceeding |3| and calculated subject-level Cook's distance, which quantifies the influence of each participant on fixed-effect estimates. For each association, the proportion of extreme residuals and the maximum Cook's distance are reported in Supplementary Table 2. Across all analyses, the proportion of observations exceeding |3| standardized residuals was small (≤0.3%), and Cook's distance values were modest, with no participant exerting disproportionate influence on model estimates. Accordingly, no outcome-based exclusions were applied.

Supplementary Table 2. Outlier analysis

| Dependent variable: Laterality of Network 1 ("Language"), Dataset 2 | | | |
|---|---|---|---|
| Independent variable of interest | Outlier proportion | Max. Cook's distance | Corresponding figure |
| age | 0.003 | 0.064 | Figure 7A |
| VDQ | 0 | 0.102 | Figure 7C |
| NVDQ | 0 | 0.200 | Figure 7E |
| Dependent variable: Laterality of Network 18 ("Default-Anterolateral"), Dataset 2 | | | |
| Independent variable of interest | Outlier proportion | Max. Cook's distance | Corresponding figure |
| age | 0 | 0.274 | Figure 7B |
| VDQ | 0 | 0.164 | Figure 7D |
| NVDQ | 0 | 0.216 | Figure 7F |
| Dependent variable: Laterality of Network 1 ("Language"), Dataset 1 | | | |
| Independent variable of interest | Outlier proportion | Max. Cook's distance | Corresponding figure |
| age | 0 | 0.060 | Supplementary Figure 25A |
| Language composite | 0 | 0.057 | Supplementary Figure 25C |
| Dependent variable: Laterality of Network 18 ("Default-Anterolateral"), Dataset 1 | | | |

| Independent variable of interest | Outlier proportion | Max. Cook's distance | Corresponding figure |
|---|---|---|---|
| age | 0 | 0.037 | Supplementary Figure 25B |
| Language composite | 0 | 0.069 | Supplementary Figure 25D |

Non-parametric permutation test for brain-behavioral relationship

To validate the statistical significance of associations between functional network laterality and behavioral measures, we performed nonparametric permutation testing in addition to the parametric linear mixed-effects analyses reported in the main text. Because behavioral measures varied across scan sessions and some children contributed multiple scans, permutation testing was implemented using a subject-level residual sign-flipping procedure derived from a reduced mixed-effects model (Winkler et al., 2014). Specifically, we first fit a reduced model including age as a fixed effect and a random intercept for subject (LI ~ age + (1 | subject)) to obtain fitted values and residuals under the null hypothesis of no behavioral effect. Residuals were then randomly sign-flipped at the subject level, preserving the within-subject dependence structure and the effects of age. For each permutation, a permuted outcome was reconstructed by adding the sign-flipped residuals to the fitted values, and the full mixed-effects model including the behavioral predictor was refit. Repeating this procedure across 1000 permutations yielded a null distribution of the behavioral regression coefficient, against which the observed coefficient was compared to obtain an empirical two-sided permutation p-value (minimum 0.001).

Permutation testing confirmed that the observed association between Language network laterality and VDQ remained statistically significant ($p_{perm} = 0.012$), consistent with the parametric mixed-effects result reported in the main text. In contrast, associations involving NVDQ did not reach significance under permutation testing ($p_{perm} = 0.150$), in line with parametric estimates. Default Anterolateral network and VDQ ($p_{perm} = 0.116$) or NVDQ ($p_{perm} = 0.533$) associations were also not significant. Together, these results indicate that the reported language-specific laterality effects are robust to distributional assumptions and are not driven by violations of model assumptions or repeated-measures structure.

Comparison with alternative templates from a generic group-average network atlas

An alternative to a data-driven clustered centroid on FC profiles of all individuals for the network templates (Figure 1) is to use the networks assignments from group-average FC (Tu-19 networks (Tu et al., 2025), Supplementary Figure 26). As predicted, the quality of separation of the FC profiles was much lower when the network assignments from group-average FC were used (average silhouette index = 0.0337, Supplementary Figure 27A-B), as opposed to the optimized clusters across the population (average silhouette index = 0.1551, Supplementary Figure 27C-D). The distribution of correlation values between the individual FC profiles to their cluster centroids are higher and less skewed in the optimized clusters than the cluster assignments from the Tu-19 networks based on group-average FC (Supplementary Figure 28), indicating that the FC separation is better defined using the 23-network solution.

We repeated the template matching procedure using the group-average prior (Tu-19 networks). For most networks that appear in both the data-driven and group-average priors, their spatial topographies were largely similar. However, the data-driven prior more finely differentiated certain regions (Supplementary Figure 29). For example, it separates the “Dorsal Somatomotor” found in the group-average prior into two distinct networks: a refined “Dorsal Somatomotor” network, encompassing hand and foot effector regions, and a “Somatic-Cognitive Action” network for integrating goals, physiology and body movement (Gordon et al., 2023). Additionally, when using the group-average prior, we observed broader regions of low template correlation around network boundaries (Supplementary Figure 30). This likely reflects the fact that the group-average templates are spatially blurred and may contain mixed signals from adjacent networks, leading to low-confidence assignments at the edges under the winner-takes-all matching approach.

#### Supplementary Figures

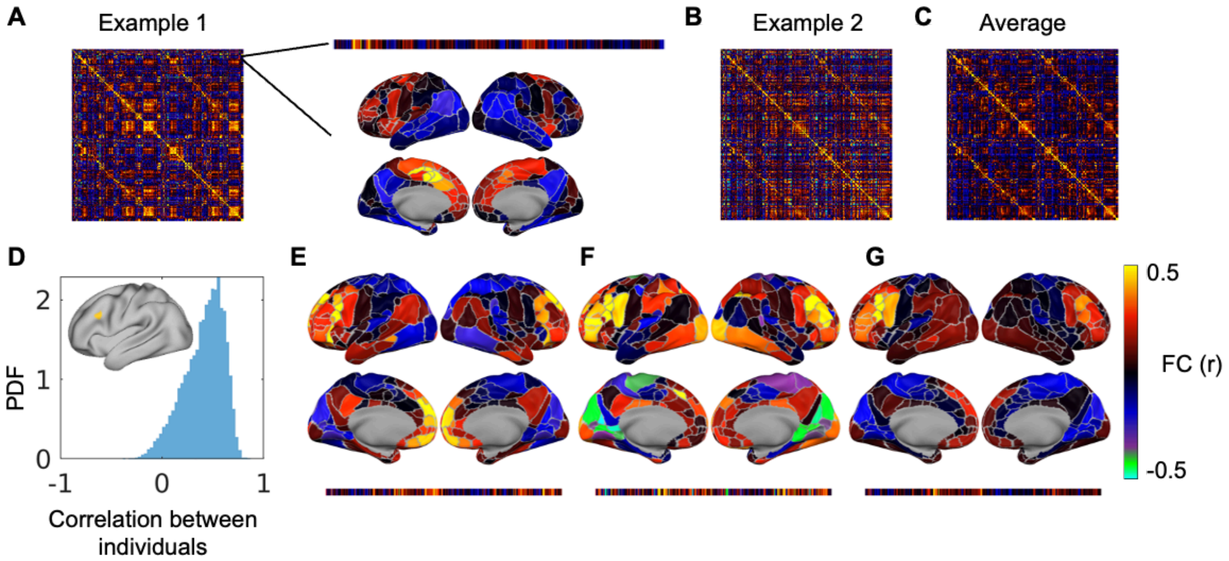

**Supplementary Figure 1**. *Functional connectivity (FC) variability across individuals.* A) Parcel-to-parcel FC matrix from one example session. The individual rows can be seen as the FC from one example parcel to all cortical parcels, which can also be visualized on a standard brain surface. B) FC from another session. C) Average FC across all 81 template sessions. D) The correlation between pairs of FC profiles (out of 81 template sessions) of one example parcel at the lateral prefrontal cortex. E-F) Example pairs of FC profiles of the example parcel. G) Average FC of the example parcel across 81 template sessions.

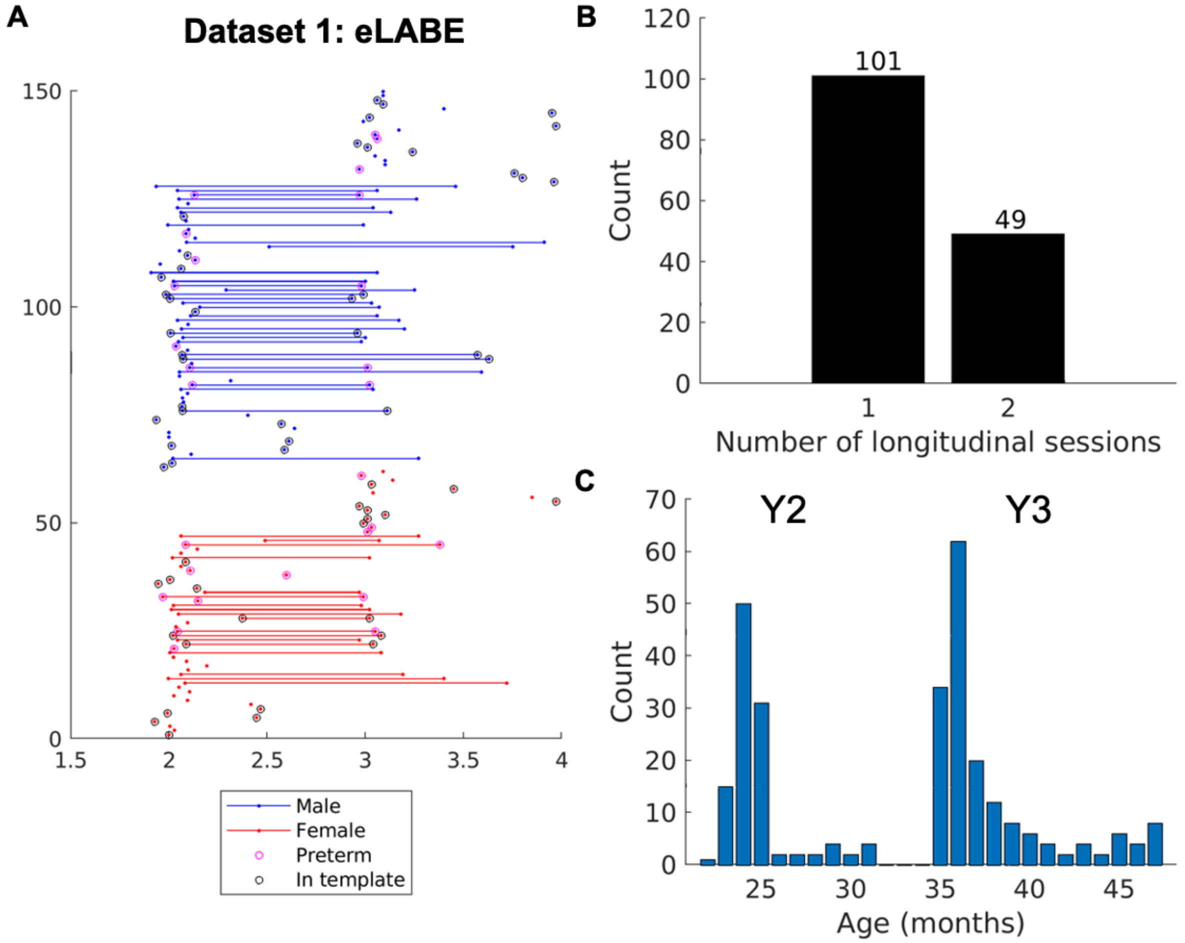


**Supplementary Figure 2**. *Dataset demographics (eLABE).* A) The age at fMRI session for each participant, with participants sorted by Male (blue) and Female (red), with preterm participants marked with magenta circles and sessions used in creating the template marked with black circles. B) Number of participants with 1-6 longitudinal sessions. C) Number of sessions each month.

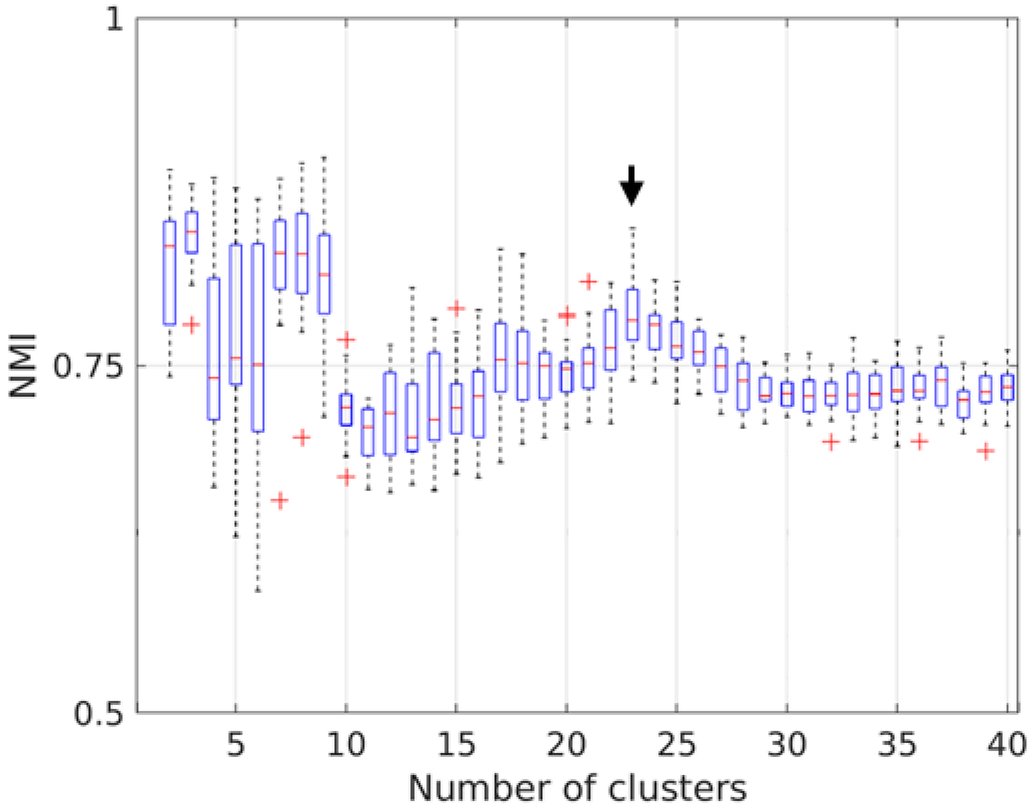

**Supplementary Figure 3**. *K-means clustering stability across 20 split-half samples of sessions.*

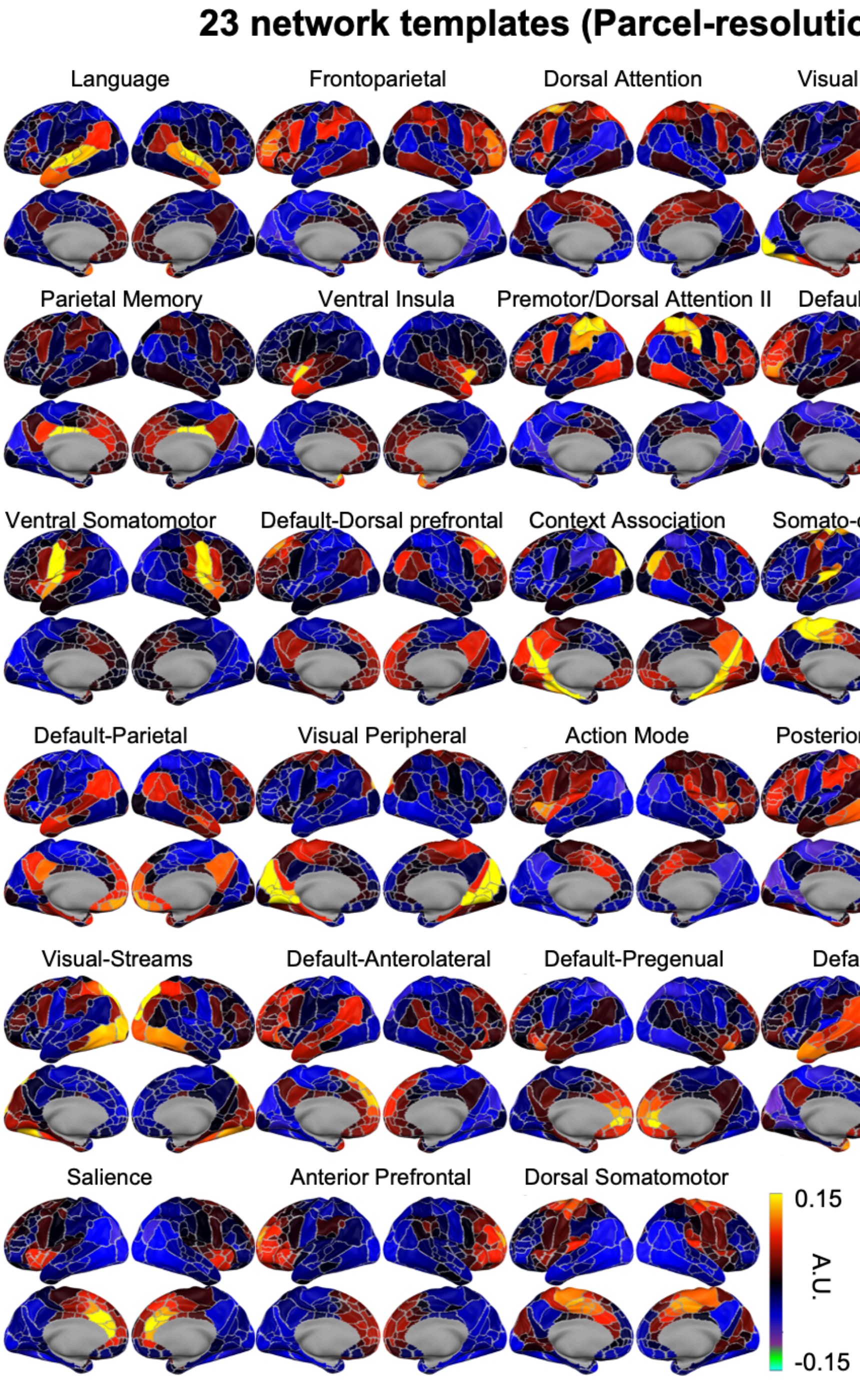


**Supplementary Figure 4**. *23 network templates at the resolution of parcels.* The values represent the mean of normalized vectors (a.k.a. cluster centroids using correlational distance) in arbitrary units (A.U.).

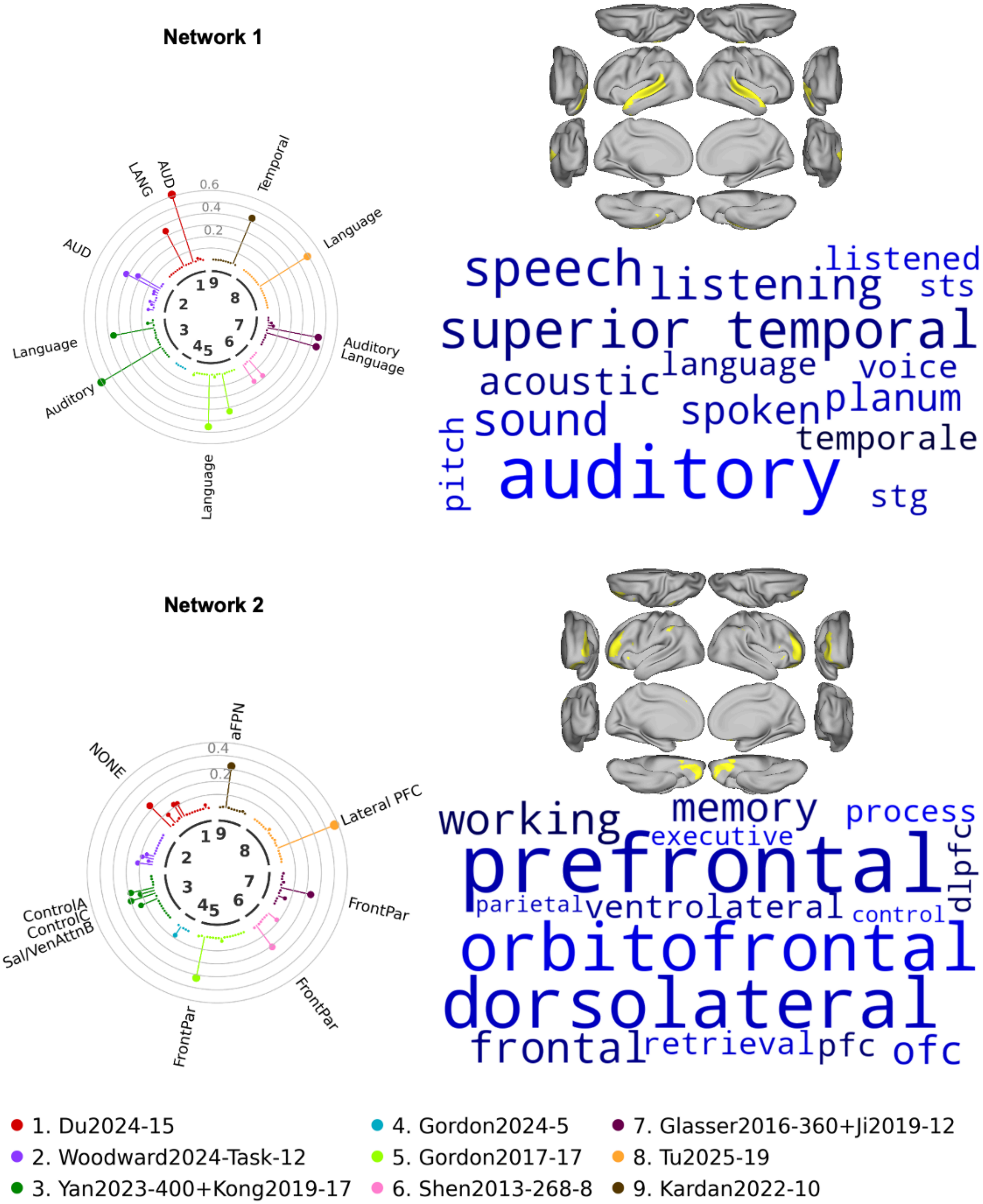


**Supplementary Figure 5**. *Correspondence between the 23 networks and previously reported functional networks/decoded terms (Networks 1-2)*. Circular bar chart plot from the network correspondence toolbox for spatial correspondence with the network locations with >0.35 probability and 7 atlases from the literature. Circles illustrate the magnitude of the dice coefficient in intervals of 0.1. Only significant networks from the spatial permutation test ($p<0.05$) were named. Top right: the network locations with >0.35 probability visualized on a template brain surface. Top bottom: terms with highly correlated meta-analytic activation maps to the network topography.

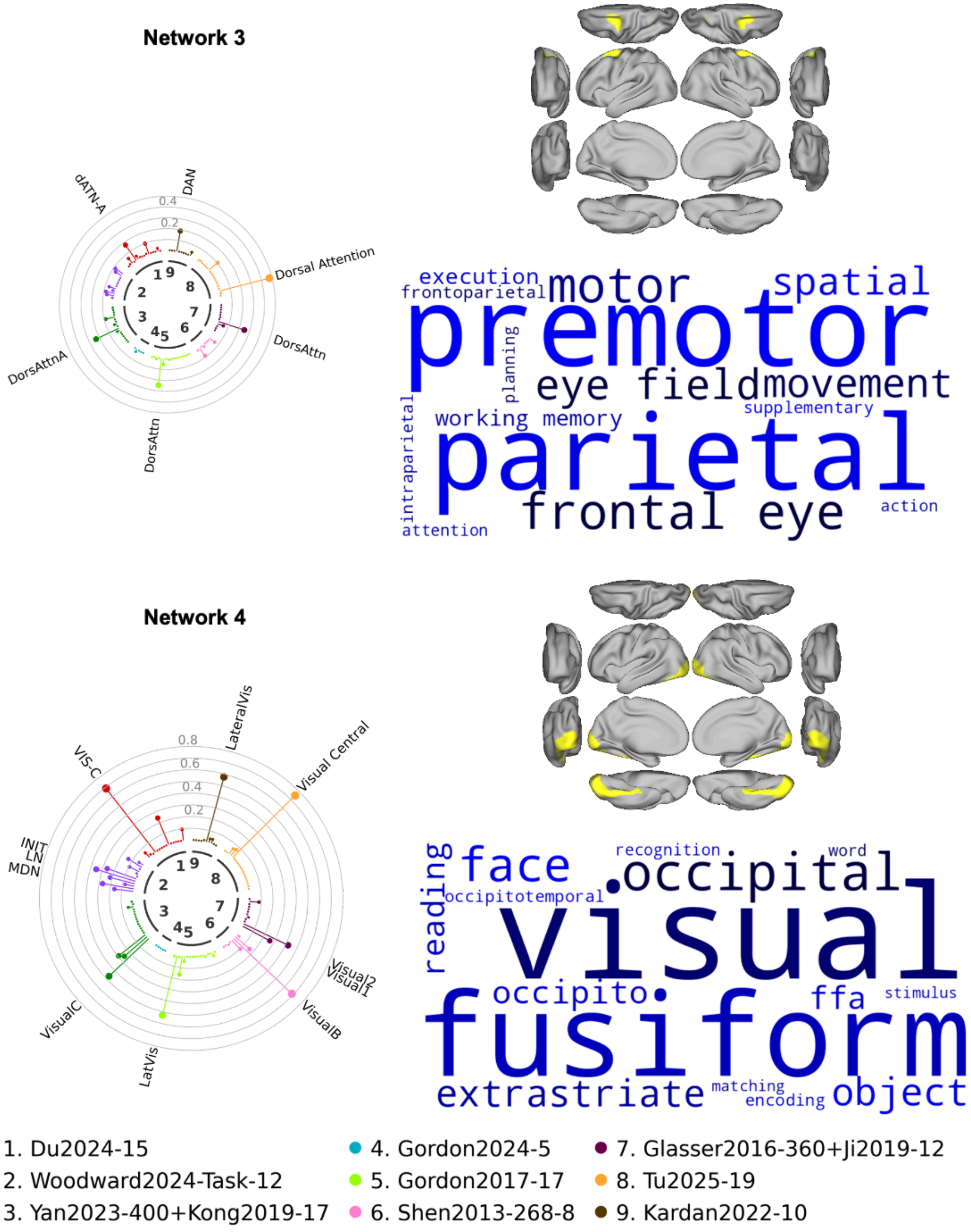


**Supplementary Figure 6**. *Correspondence between the 23 networks and previously reported functional networks/decoded terms (Networks 3-4)*.Left: Circular bar chart plot from the network correspondence toolbox for spatial correspondence with the network locations with >0.35 probability and 7 atlases from the literature. Circles illustrate the magnitude of the dice coefficient in intervals of 0.1. Only significant networks from the spatial permutation test ($p<0.05$) were named. Top right: the network locations with >0.35 probability visualized on a template brain surface. Top bottom: terms with highly correlated meta-analytic activation maps to the network topography.

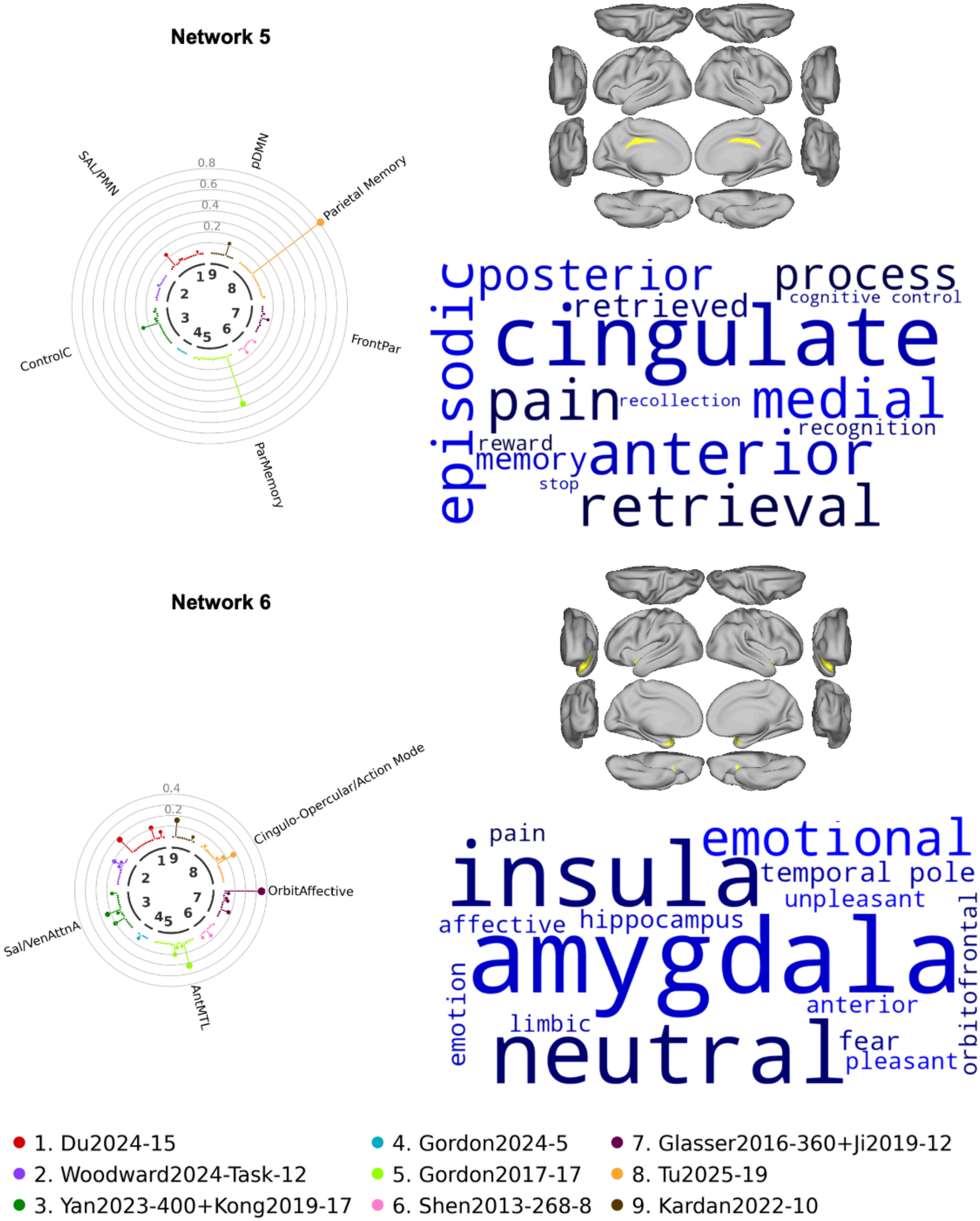
Network 5
SAL/PMN
pDMN
Parietal Memory
FrontPar
ParMemory
ControlC
0.8
0.6
0.4
0.2
posterior
process
cognitive control
retrieved
cingulate
episodic
pain
recollection
medial
recognition
reward
memory
anterior
stop
retrieval
Network 6
Cingulo-Opercular/Action Mode
OrbitAffective
Sal/VenAttnA
AntMTL
0.4
0.2
pain
emotional
insula
temporal pole
unpleasant
affective
hippocampus
orbitofrontal
amygdala
emotion
anterior
limbic
fear
pleasant
neutral
1. Du2024-15
2. Woodward2024-Task-12
3. Yan2023-400+Kong2019-17
4. Gordon2024-5
5. Gordon2017-17
6. Shen2013-268-8
7. Glasser2016-360+Ji2019-12
8. Tu2025-19
9. Kardan2022-10

**Supplementary Figure 7**. *Correspondence between the 23 networks and previously reported functional networks/decoded terms (Networks 5-6)*. Left: Circular bar chart plot from the network correspondence toolbox for spatial correspondence with the network locations with >0.35 probability and 7 atlases from the literature. Circles illustrate the magnitude of the dice coefficient in intervals of 0.1. Only significant networks from the spatial permutation test ($p<0.05$) were named. Top right: the network locations with >0.35 probability visualized on a template brain surface. Top bottom: terms with highly correlated meta-analytic activation maps to the network topography.

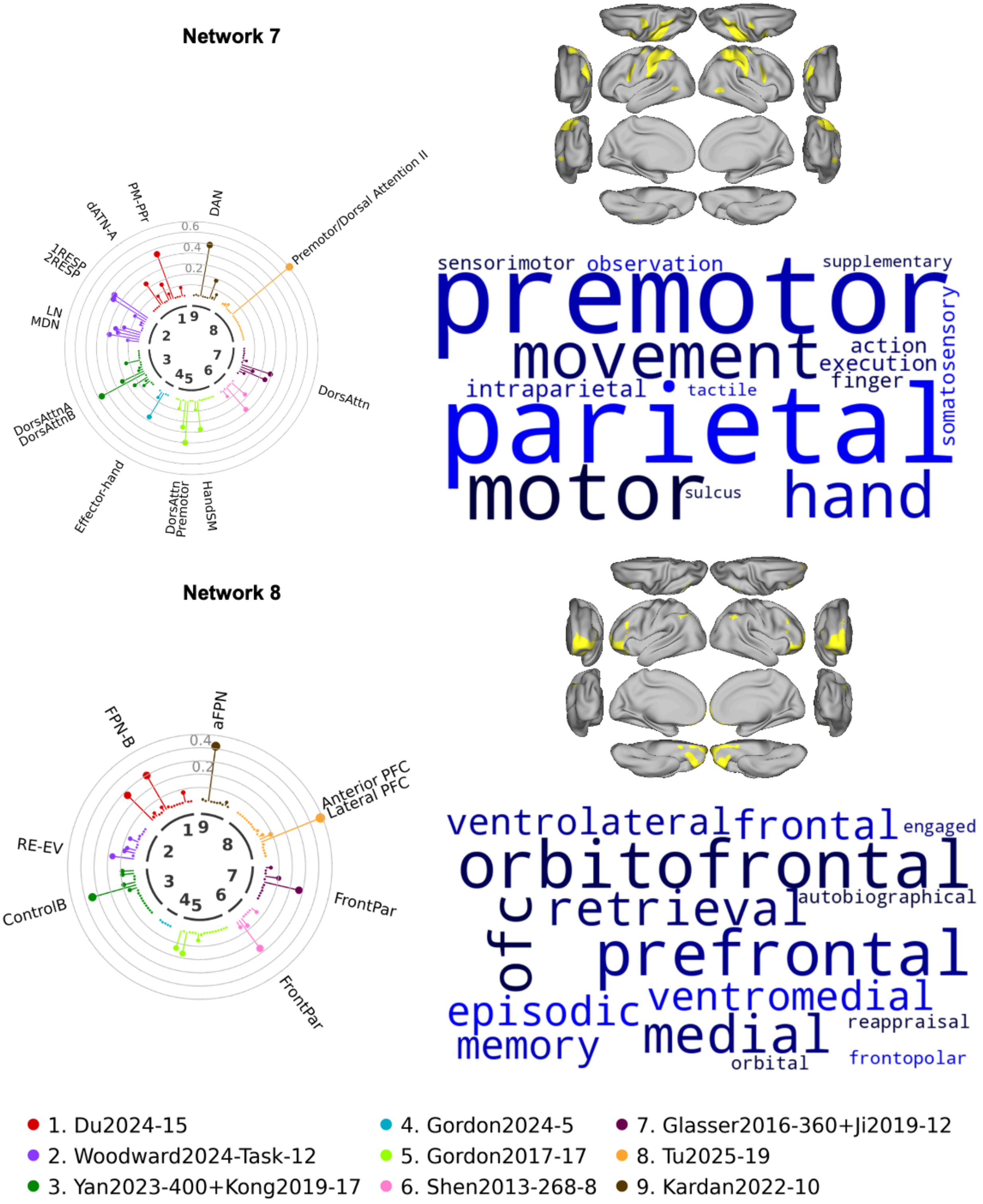

Network 7
DAN
PM-PPr
dATN-A
1RESP
2RESP
LN
MDN
DorsAttnA
DorsAttnB
Effector-hand
DorsAttn
Premotor
HandSM
DorsAttn
Premotor/Dorsal Attention II
0.6
0.4
0.2
premotor
parietal
motor
movement
hand
sensorimotor
observation
supplementary
somatosensory
action
execution
finger
intraparietal
tactile
sulcus
Network 8
aFPN
FPN-B
RE-EV
ControlB
FrontPar
FrontPar
Anterior PFC
Lateral PFC
0.4
0.2
orbitofrontal
prefrontal
ventrolateral
frontal
engaged
retrieval
autobiographical
ofc
ventromedial
episodic
medial
memory
reappraisal
orbital
frontopolar
1. Du2024-15
2. Woodward2024-Task-12
3. Yan2023-400+Kong2019-17
4. Gordon2024-5
5. Gordon2017-17
6. Shen2013-268-8
7. Glasser2016-360+Ji2019-12
8. Tu2025-19
9. Kardan2022-10

**Supplementary Figure 8**. *Correspondence between the 23 networks and previously reported functional networks/decoded terms (Networks 7-8)*. Left: Circular bar chart plot from the network correspondence toolbox for spatial correspondence with the network locations with >0.35 probability and 7 atlases from the literature. Circles illustrate the magnitude of the dice coefficient in intervals of 0.1. Only significant networks from the spatial permutation test ($p<0.05$) were named. Top right: the network locations with >0.35 probability visualized on a template brain surface. Top bottom: terms with highly correlated meta-analytic activation maps to the network topography.

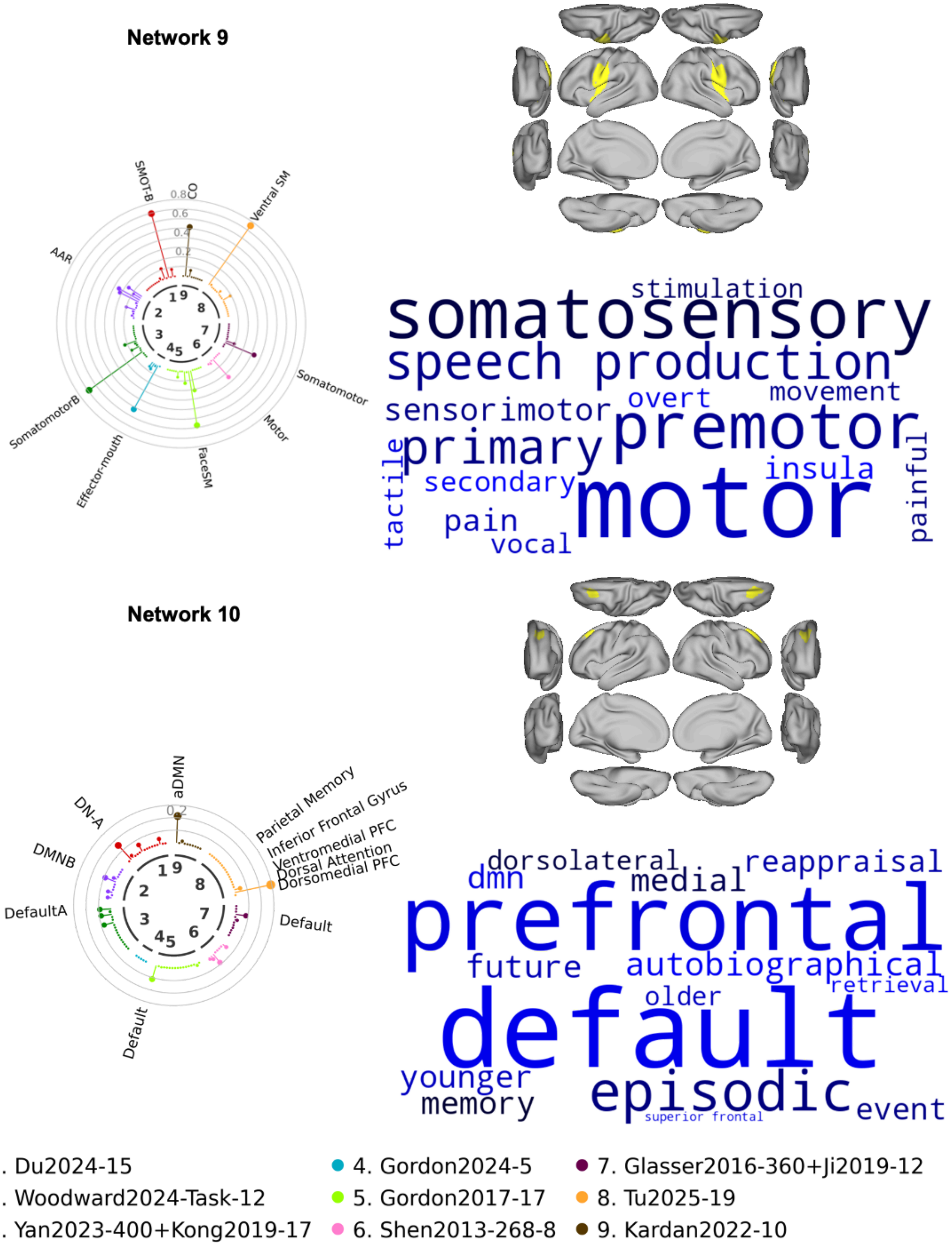


**Supplementary Figure 9**. *Correspondence between the 23 networks and previously reported functional networks/decoded terms (Networks 9-10)*. Left: Circular bar chart plot from the network correspondence toolbox for spatial correspondence with the network locations with

>0.35 probability and 7 atlases from the literature. Circles illustrate the magnitude of the dice coefficient in intervals of 0.1. Only significant networks from the spatial permutation test ($p<0.05$) were named. Top right: the network locations with >0.35 probability visualized on a template brain surface. Top bottom: terms with highly correlated meta-analytic activation maps to the network topography.

**Network 11**

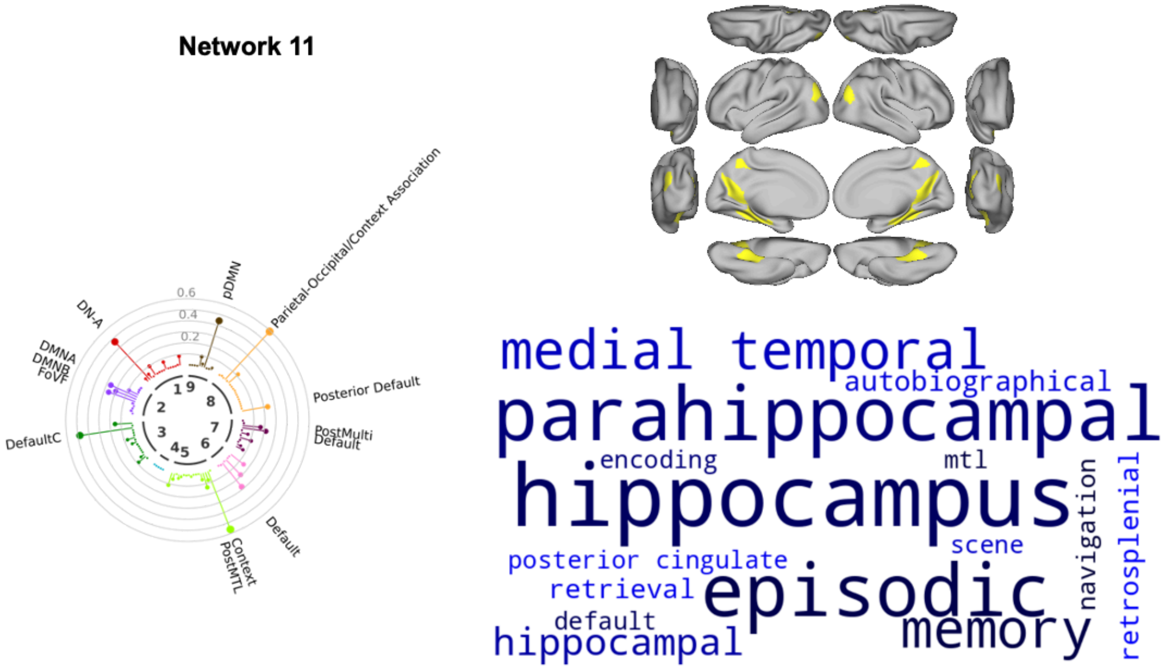


**Network 12**

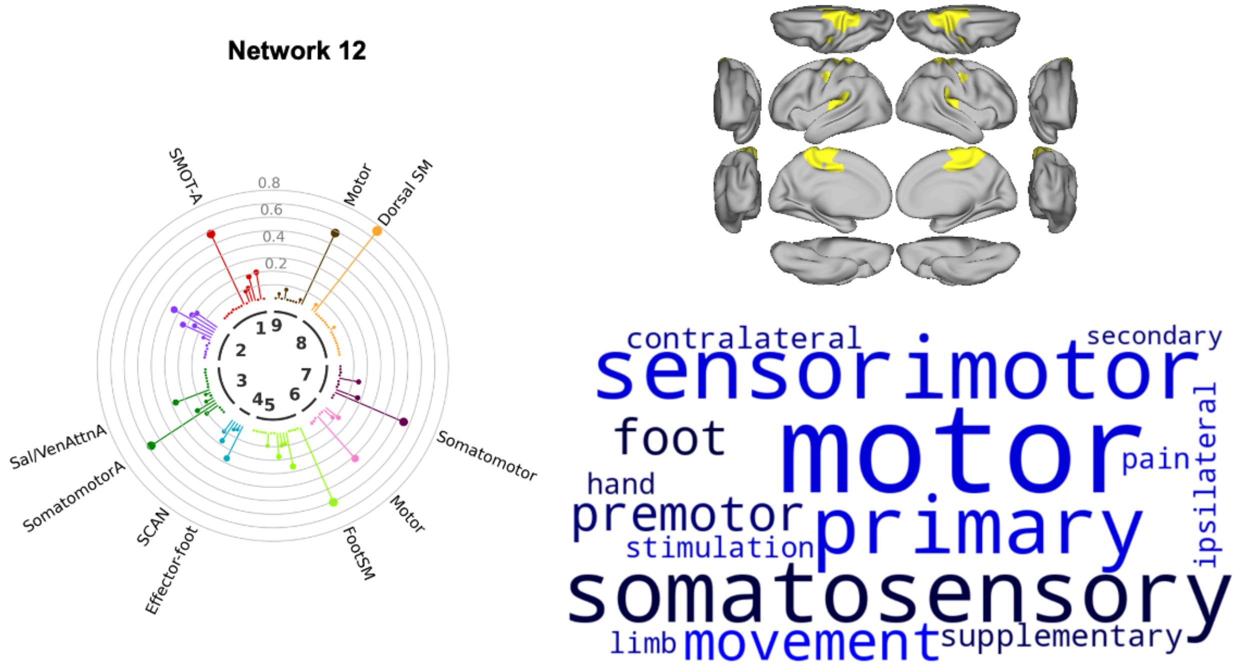


- 1. Du2024-15
- 2. Woodward2024-Task-12
- 3. Yan2023-400+Kong2019-17
- 4. Gordon2024-5
- 5. Gordon2017-17
- 6. Shen2013-268-8
- 7. Glasser2016-360+Ji2019-12
- 8. Tu2025-19
- 9. Kardan2022-10

**Supplementary Figure 10**. *Correspondence between the 23 networks and previously reported functional networks/decoded terms (Networks 11-12)*. Left: Circular bar chart plot from the network correspondence toolbox for spatial correspondence with the network locations with >0.35 probability and 7 atlases from the literature. Circles illustrate the magnitude of the dice coefficient in intervals of 0.1. Only significant networks from the spatial permutation test ($p<0.05$) were named. Top right: the network locations with >0.35 probability visualized on a template brain surface. Top bottom: terms with highly correlated meta-analytic activation maps to the network topography.

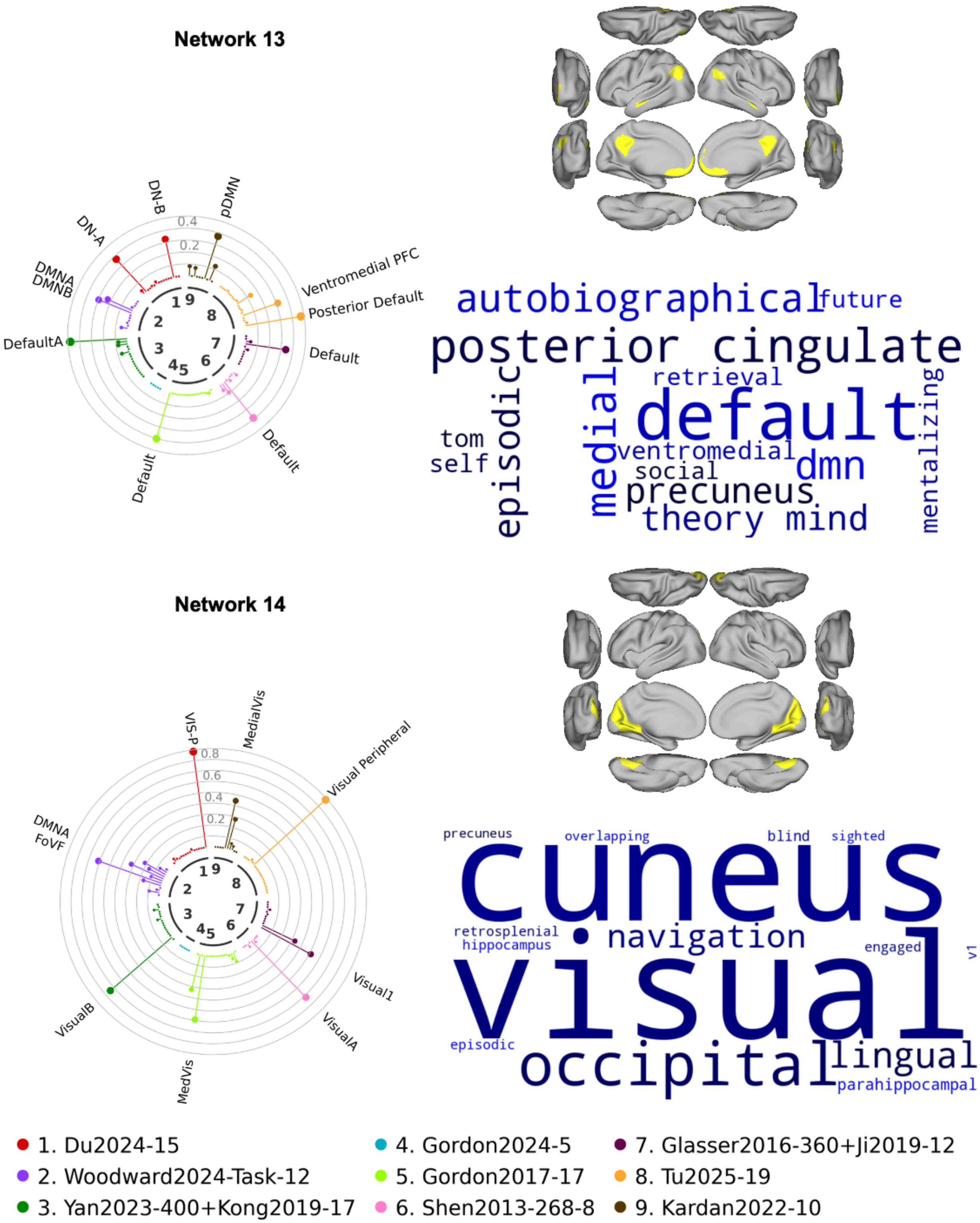

Network 13
DN-B
pDMN
DN-A
DMNA
DMNB
Ventromedial PFC
Posterior Default
DefaultA
Default
Default
Default
0.4
0.2
autobiographical
future
posterior cingulate
retrieval
episodic
medial
default
mentalizing
tom
self
ventromedial
social
dmn
precuneus
theory mind
Network 14
VIS-P
MedialVis
Visual Peripheral
DMNA
FoVF
Visual1
VisualA
VisualB
MedVis
0.8
0.6
0.4
0.2
precuneus
overlapping
blind
sighted
cuneus
retrosplenial
hippocampus
navigation
engaged
v1
visual
episodic
occipital
lingual
parahippocampal
1. Du2024-15
2. Woodward2024-Task-12
3. Yan2023-400+Kong2019-17
4. Gordon2024-5
5. Gordon2017-17
6. Shen2013-268-8
7. Glasser2016-360+Ji2019-12
8. Tu2025-19
9. Kardan2022-10

**Supplementary Figure 11**. *Correspondence between the 23 networks and previously reported functional networks/decoded terms (Networks 13-14)*. Left: Circular bar chart plot from the network correspondence toolbox for spatial correspondence with the network locations with >0.35 probability and 7 atlases from the literature. Circles illustrate the magnitude of the dice coefficient in intervals of 0.1. Only significant networks from the spatial permutation test ($p<0.05$) were named. Top right: the network locations with >0.35 probability visualized on a template brain surface. Top bottom: terms with highly correlated meta-analytic activation maps to the network topography.

**Network 15**

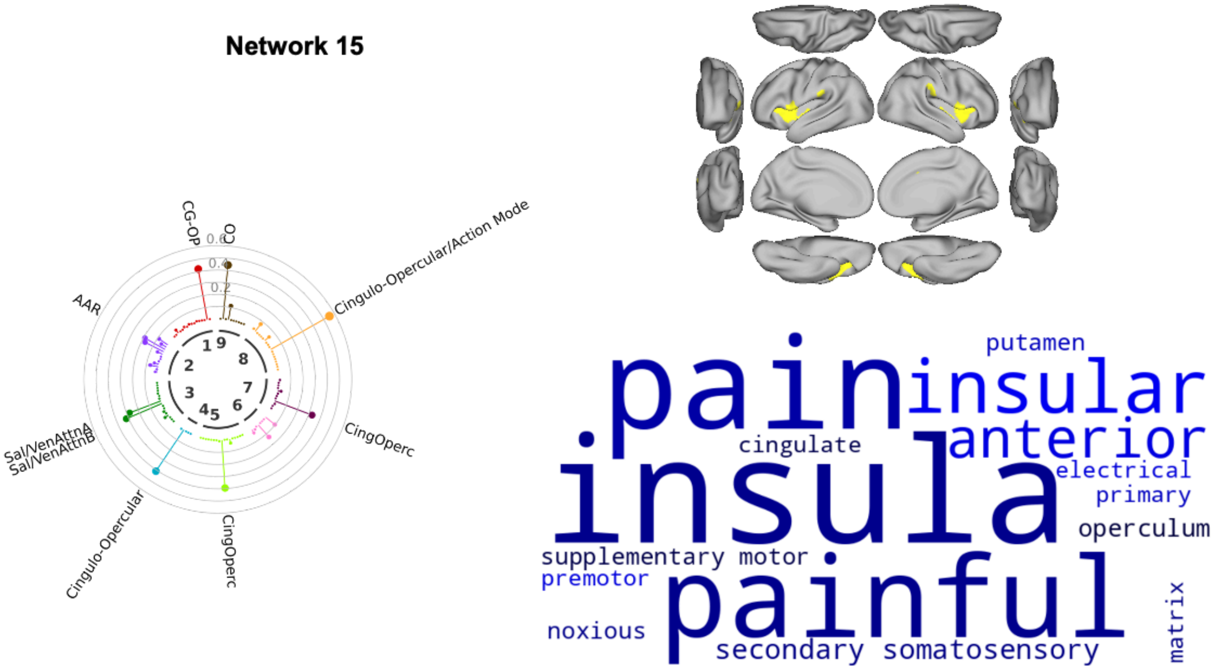


**Network 16**

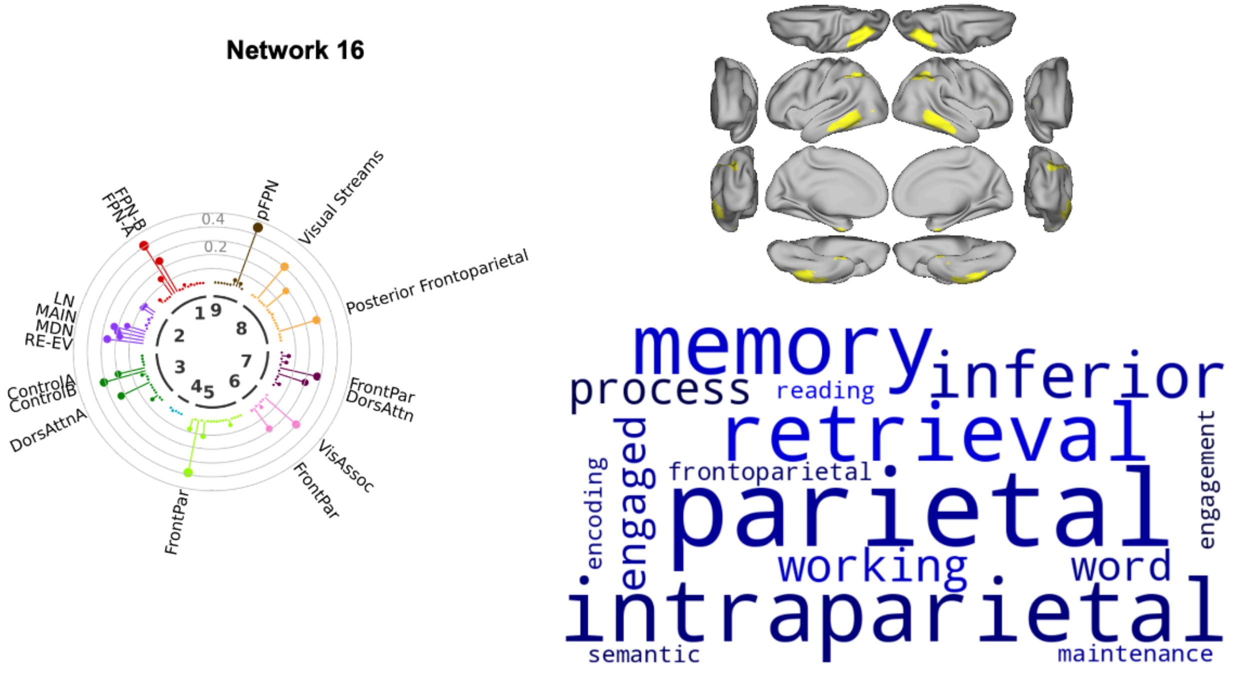


- 1. Du2024-15
- 2. Woodward2024-Task-12
- 3. Yan2023-400+Kong2019-17
- 4. Gordon2024-5
- 5. Gordon2017-17
- 6. Shen2013-268-8
- 7. Glasser2016-360+Ji2019-12
- 8. Tu2025-19
- 9. Kardan2022-10

**Supplementary Figure 12**. *Correspondence between the 23 networks and previously reported functional networks/decoded terms (Networks 15-16)*. Left: Circular bar chart plot from the network correspondence toolbox for spatial correspondence with the network locations with >0.35 probability and 7 atlases from the literature. Circles illustrate the magnitude of the dice coefficient in intervals of 0.1. Only significant networks from the spatial permutation test ($p<0.05$) were named. Top right: the network locations with >0.35 probability visualized on a template brain surface. Top bottom: terms with highly correlated meta-analytic activation maps to the network topography.

**Network 17**

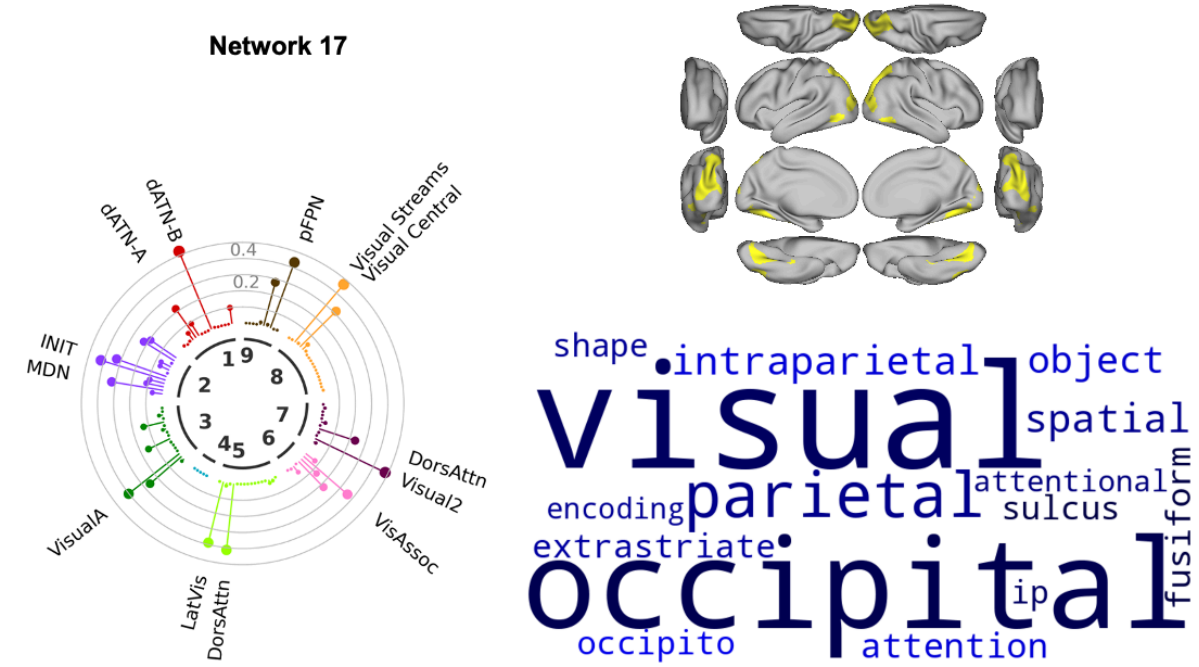


**Network 18**

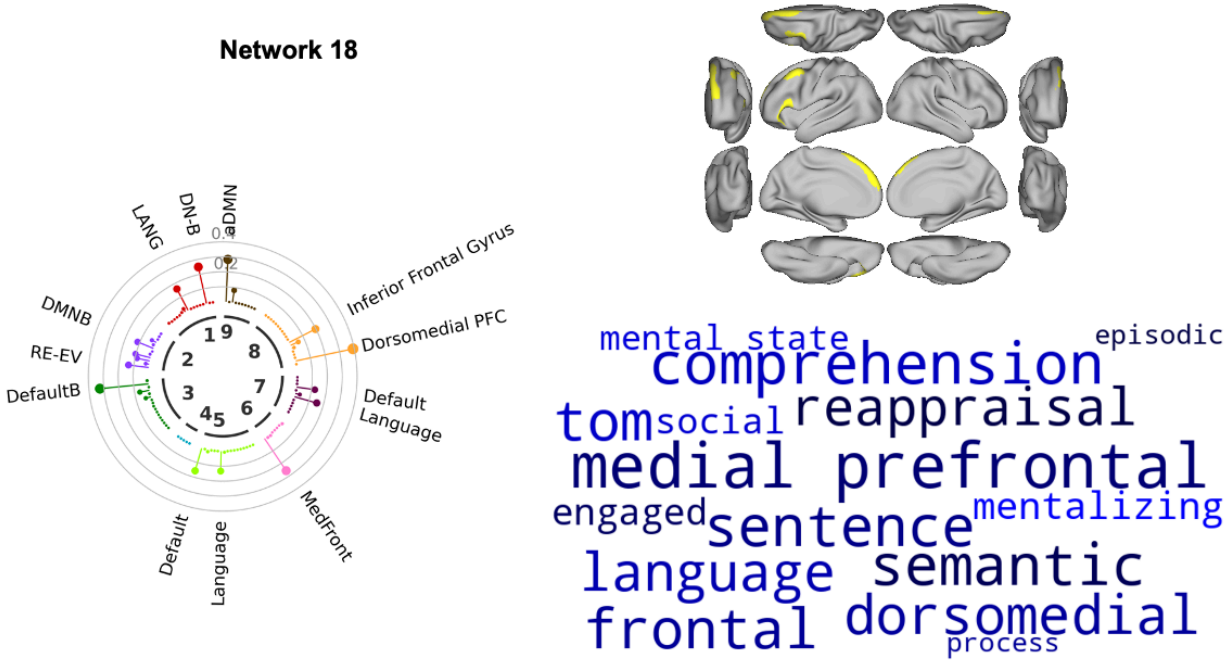


- 1. Du2024-15
- 2. Woodward2024-Task-12
- 3. Yan2023-400+Kong2019-17
- 4. Gordon2024-5
- 5. Gordon2017-17
- 6. Shen2013-268-8
- 7. Glasser2016-360+Ji2019-12
- 8. Tu2025-19
- 9. Kardan2022-10

**Supplementary Figure 13**. *Correspondence between the 23 networks and previously reported functional networks/decoded terms (Networks 17-18)*. Left: Circular bar chart plot from the network correspondence toolbox for spatial correspondence with the network locations with >0.35 probability and 7 atlases from the literature. Circles illustrate the magnitude of the dice coefficient in intervals of 0.1. Only significant networks from the spatial permutation test ($p<0.05$) were named. Top right: the network locations with >0.35 probability visualized on a template brain surface. Top bottom: terms with highly correlated meta-analytic activation maps to the network topography.

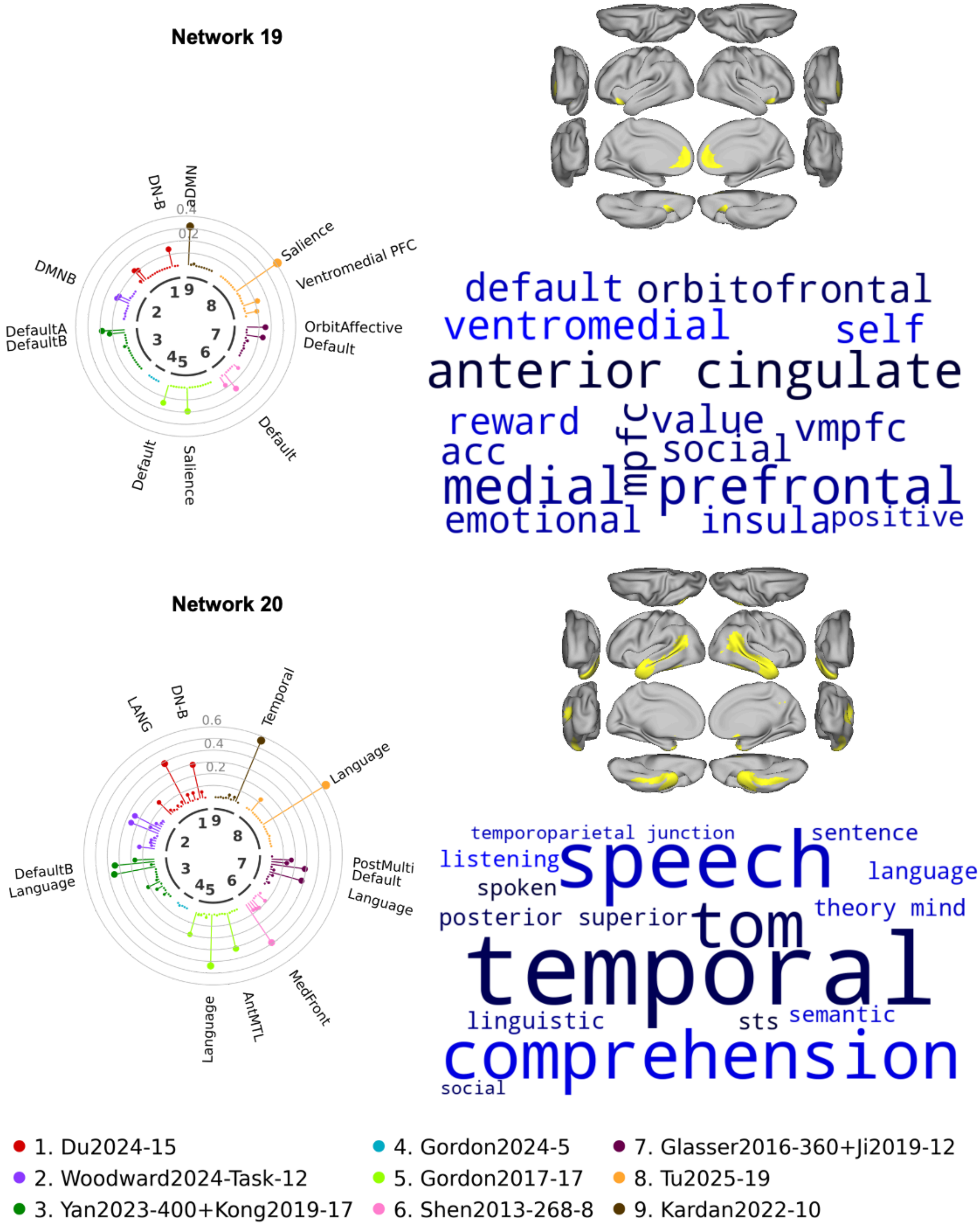
Network 19
aDMN
DN-B
DMNB
DefaultA
DefaultB
Salience
Ventromedial PFC
OrbitAffective
Default
Default
Default
Salience
0.4
0.2
default orbitofrontal
ventromedial self
anterior cingulate
reward value vmpfc
acc mpfc social
medial prefrontal
emotional insula positive
Network 20
LANG
DN-B
Temporal
Language
PostMulti
Default
Language
MedFront
AntMTL
Language
DefaultB
Language
0.6
0.4
0.2
temporoparietal junction
sentence
listening
speech
language
spoken
posterior superior
tom
theory mind
temporal
linguistic
sts
semantic
comprehension
social
1. Du2024-15
2. Woodward2024-Task-12
3. Yan2023-400+Kong2019-17
4. Gordon2024-5
5. Gordon2017-17
6. Shen2013-268-8
7. Glasser2016-360+Ji2019-12
8. Tu2025-19
9. Kardan2022-10

**Supplementary Figure 14**. *Correspondence between the 23 networks and previously reported functional networks/decoded terms (Networks 19-20)*. Left: Circular bar chart plot from the network correspondence toolbox for spatial correspondence with the network locations with >0.35 probability and 7 atlases from the literature. Circles illustrate the magnitude of the dice coefficient in intervals of 0.1. Only significant networks from the spatial permutation test ($p<0.05$) were named. Top right: the network locations with >0.35 probability visualized on a template brain surface. Top bottom: terms with highly correlated meta-analytic activation maps to the network topography.

**Network 21**

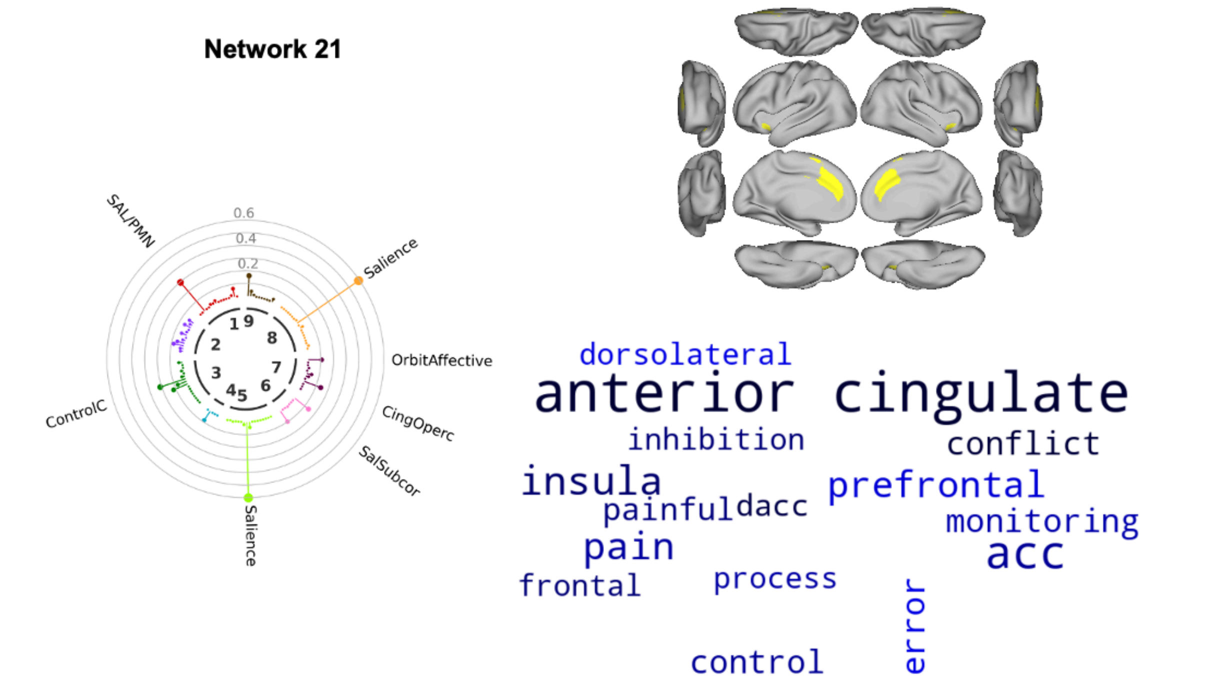


**Network 22**

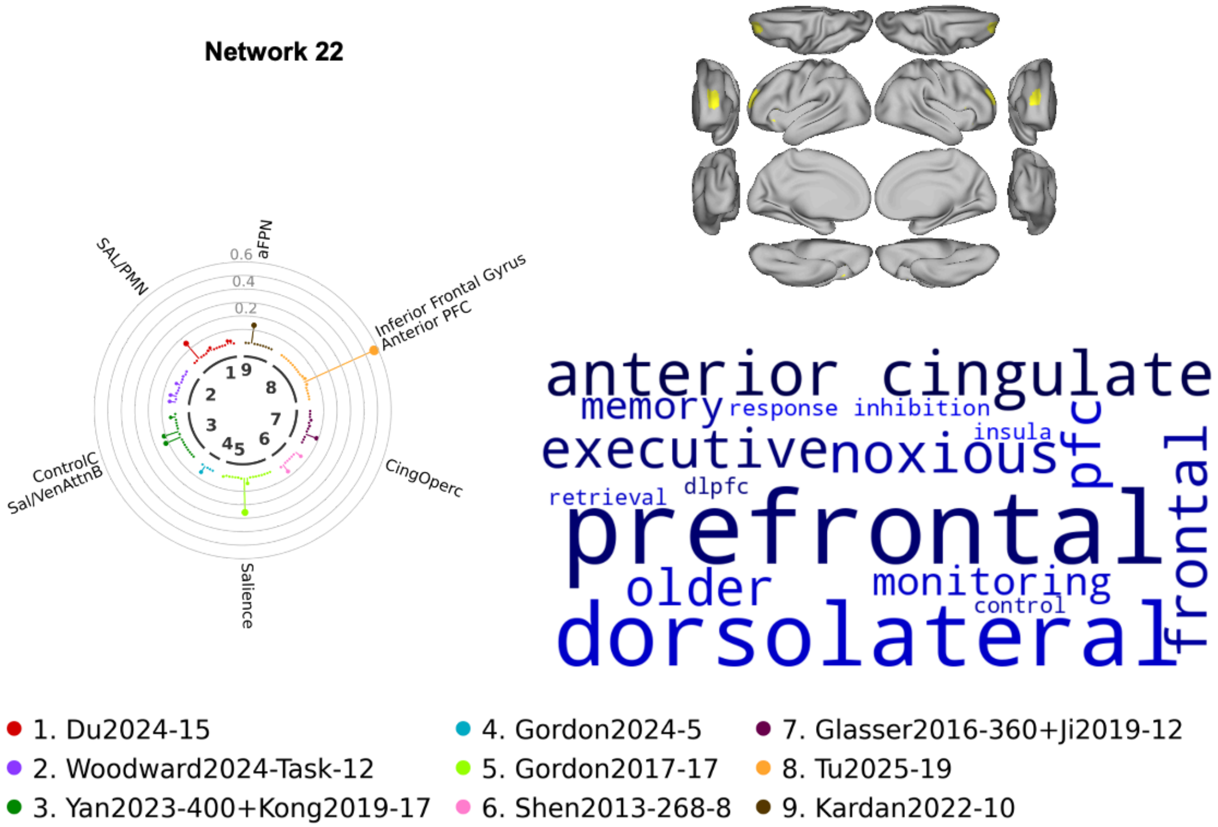

**Supplementary Figure 15**. *Correspondence between the 23 networks and previously reported functional networks/decoded terms (Networks 21-22)*. Left: Circular bar chart plot from the network correspondence toolbox for spatial correspondence with the network locations with >0.35 probability and 7 atlases from the literature. Circles illustrate the magnitude of the dice coefficient in intervals of 0.1. Only significant networks from the spatial permutation test (p<0.05) were named. Top right: the network locations with >0.35 probability visualized on a template brain surface. Top bottom: terms with highly correlated meta-analytic activation maps to the network topography.

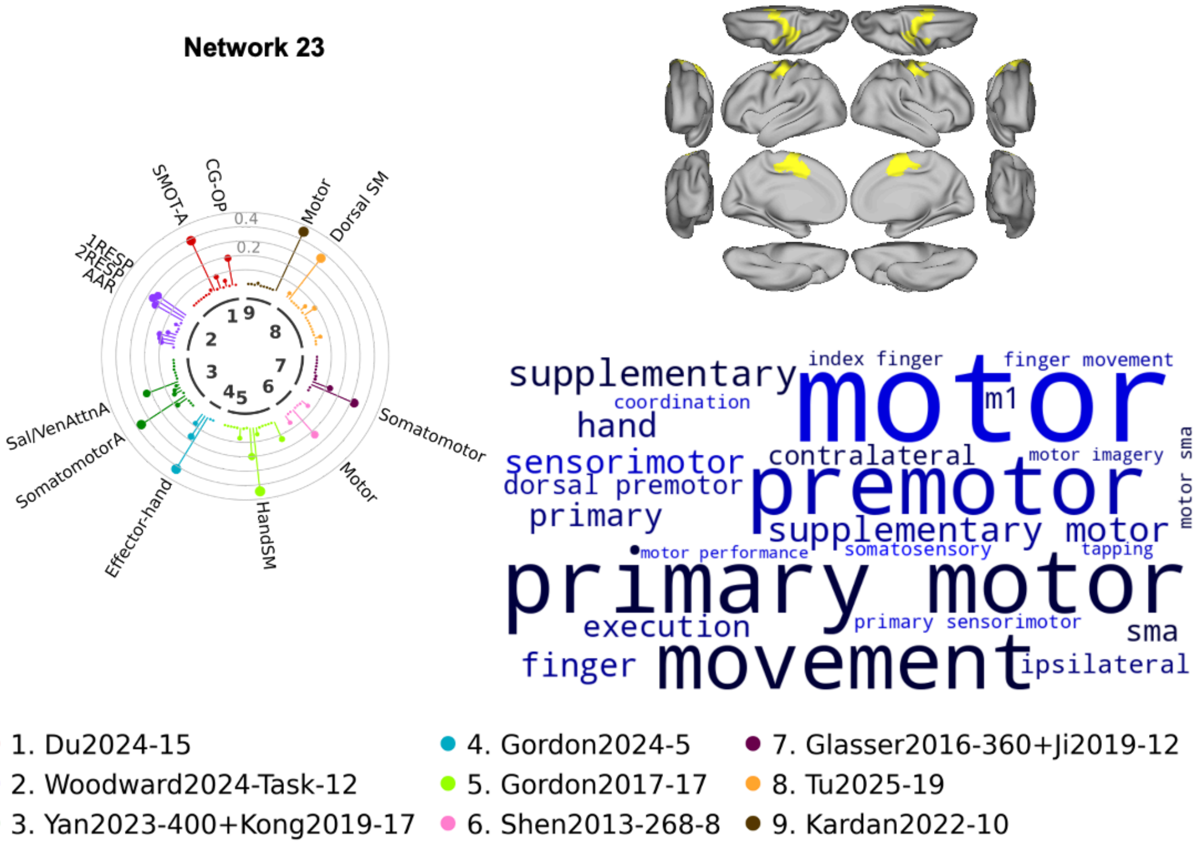

**Supplementary Figure 16**. *Correspondence between the 23 networks and previously reported functional networks/decoded terms (Networks 23)*. Left: Circular bar chart plot from the network correspondence toolbox for spatial correspondence with the network locations with >0.35 probability and 7 atlases from the literature. Circles illustrate the magnitude of the dice coefficient in intervals of 0.1. Only significant networks from the spatial permutation test ($p<0.05$) were named. Top right: the network locations with >0.35 probability visualized on a template brain surface. Top bottom: terms with highly correlated meta-analytic activation maps to the network topography.

#### 23 network population-average FC (81 template sessions)

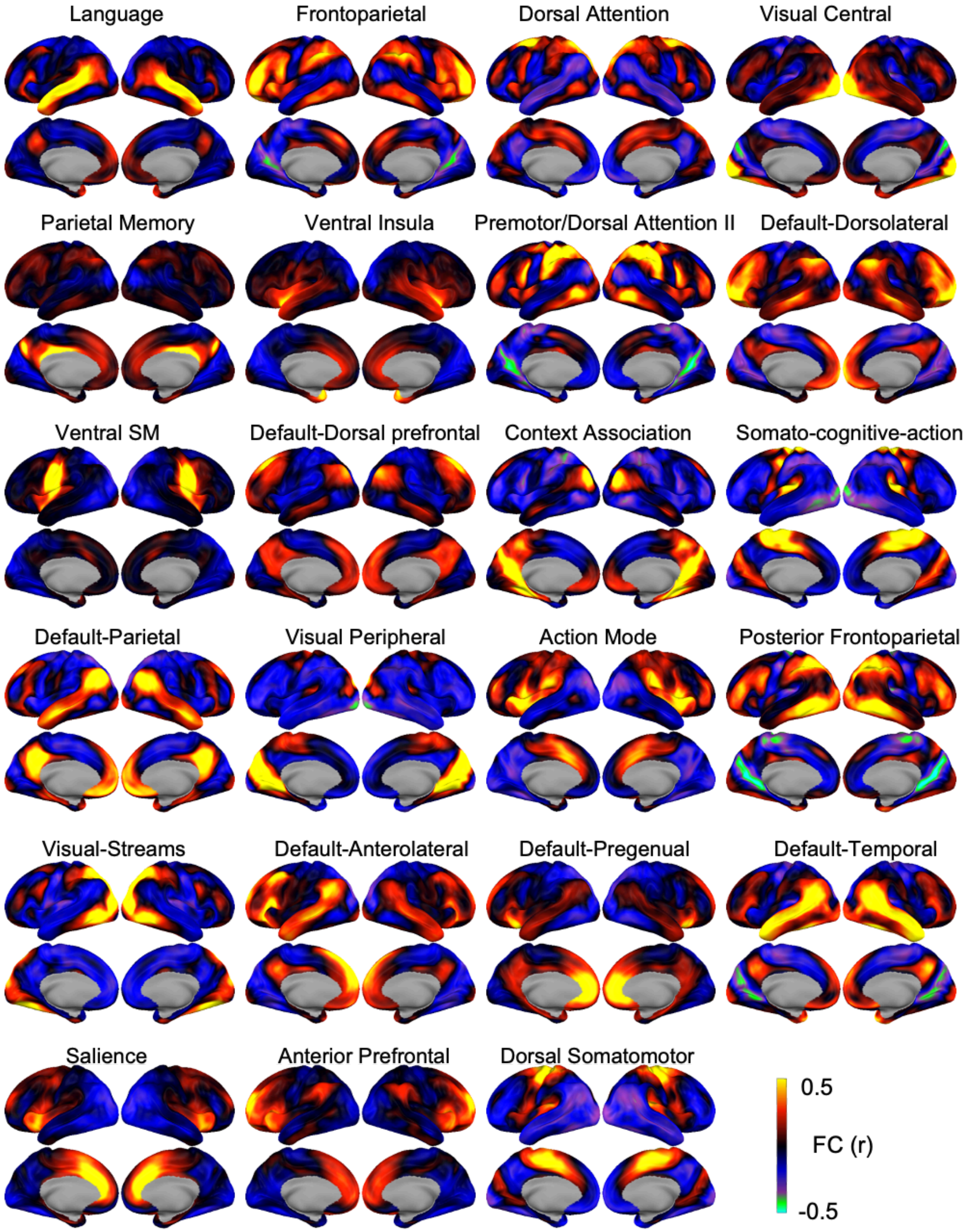

**Supplementary Figure 17**. *Population-average FC from the averaging of the network functional connectivity in each subject.*

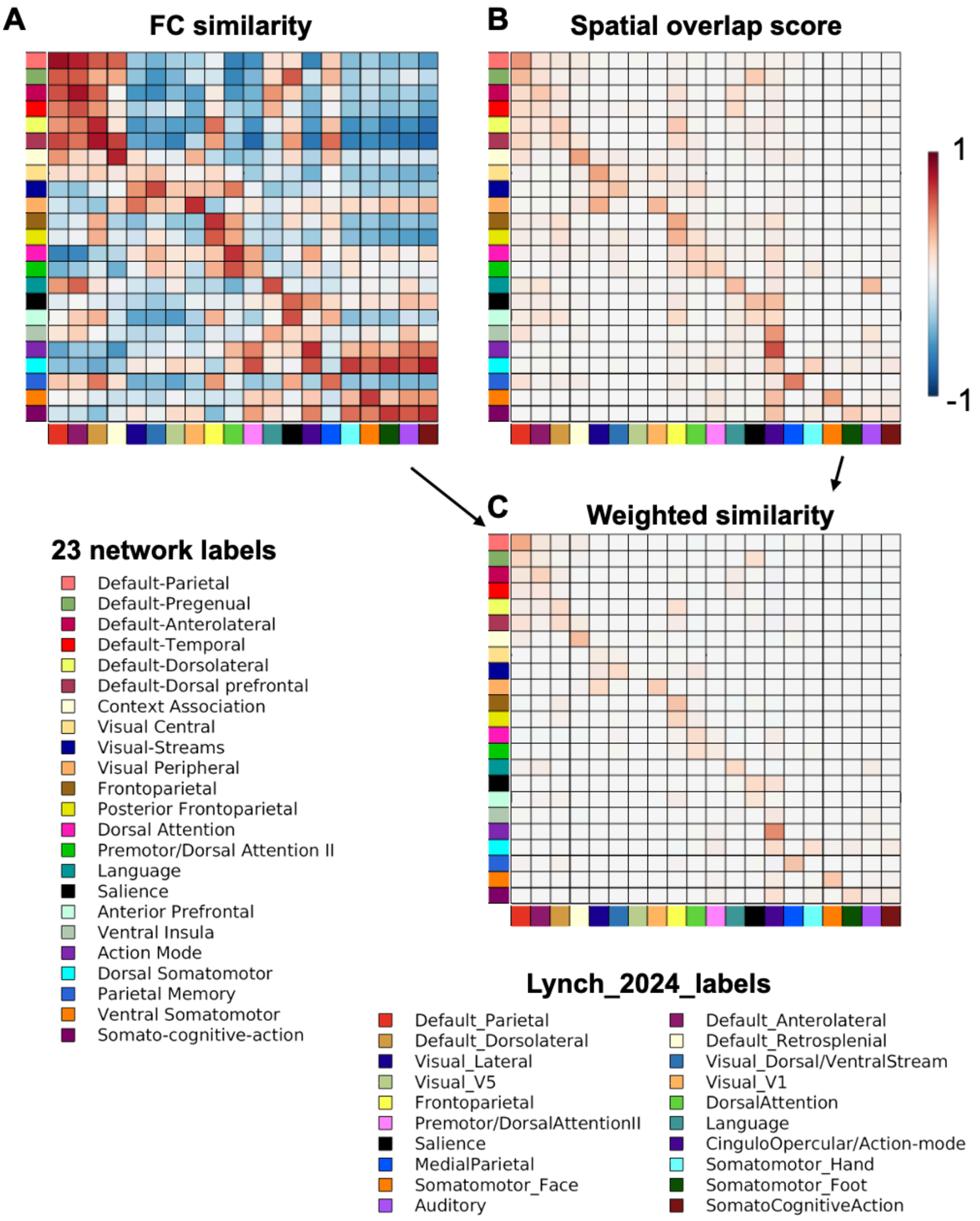

**Supplementary Figure 18**. *Similarity between networks based on functional connectivity and spatial overlap*- Matching the 23 networks (rows) to the 20 networks from Lynch 2024 (columns). A) The functional connectivity (FC) similarity. B) The spatial overlap score. C) Weighted similarity.

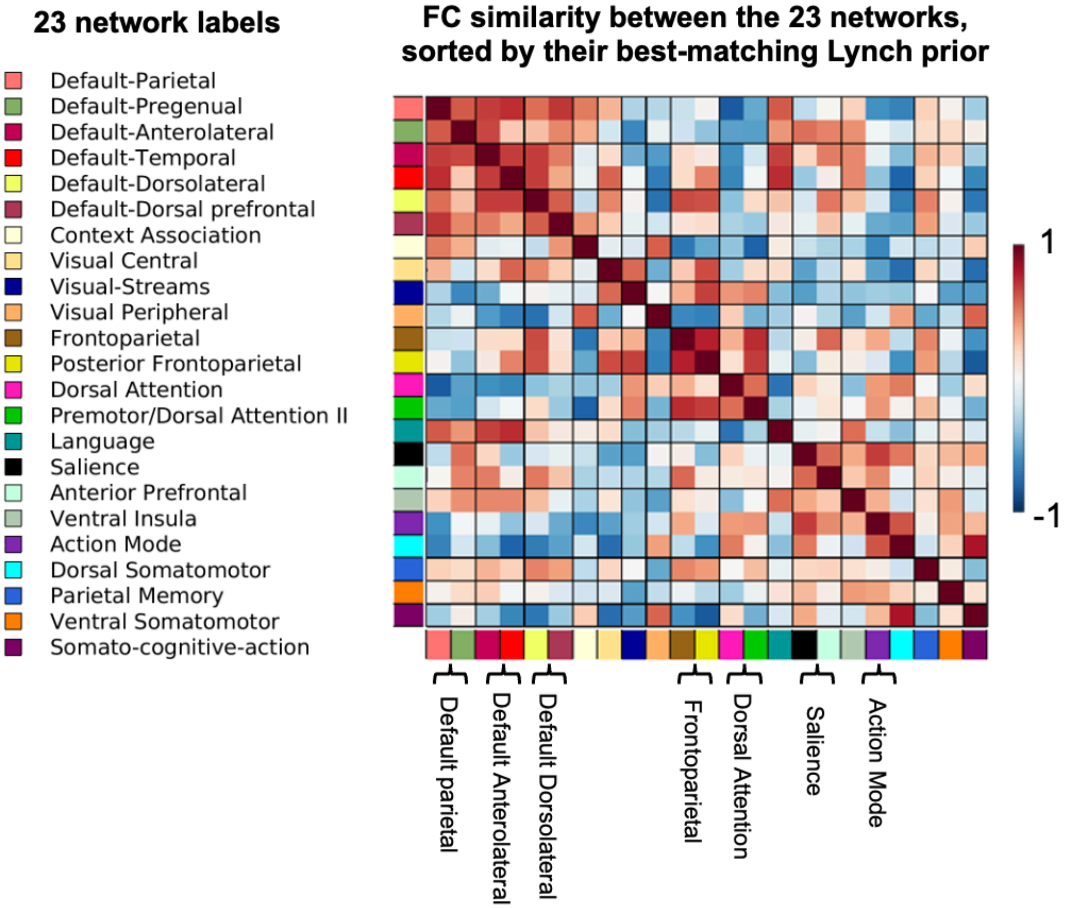


**Supplementary Figure 19**. *FC similarity between the 23 networks, sorted by their best-matching Lynch prior. The blocks show that some of the 23 networks match to the same network in Lynch 2024.*

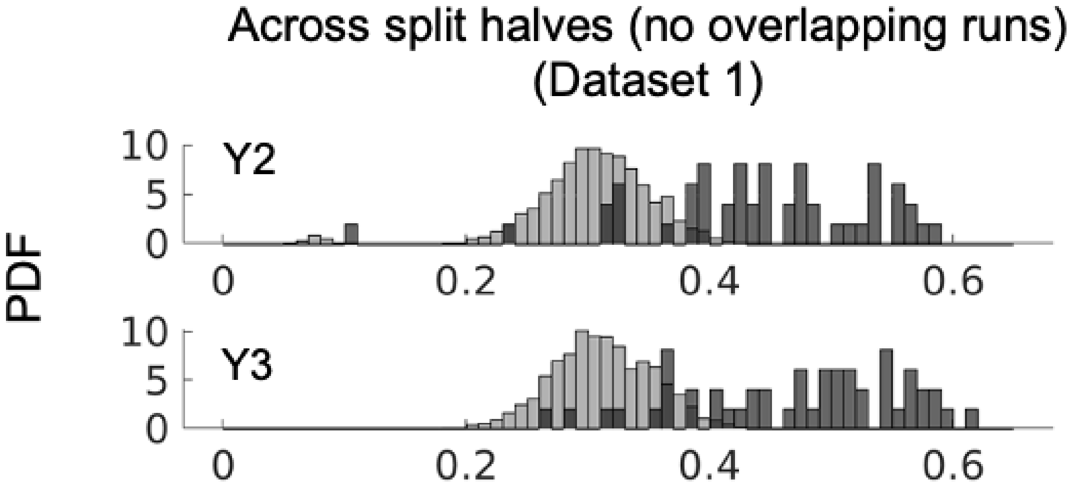


**Supplementary Figure 20**. Within versus between individual NMI across split halves (Dataset 1) using split halves with no overlaps in runs.

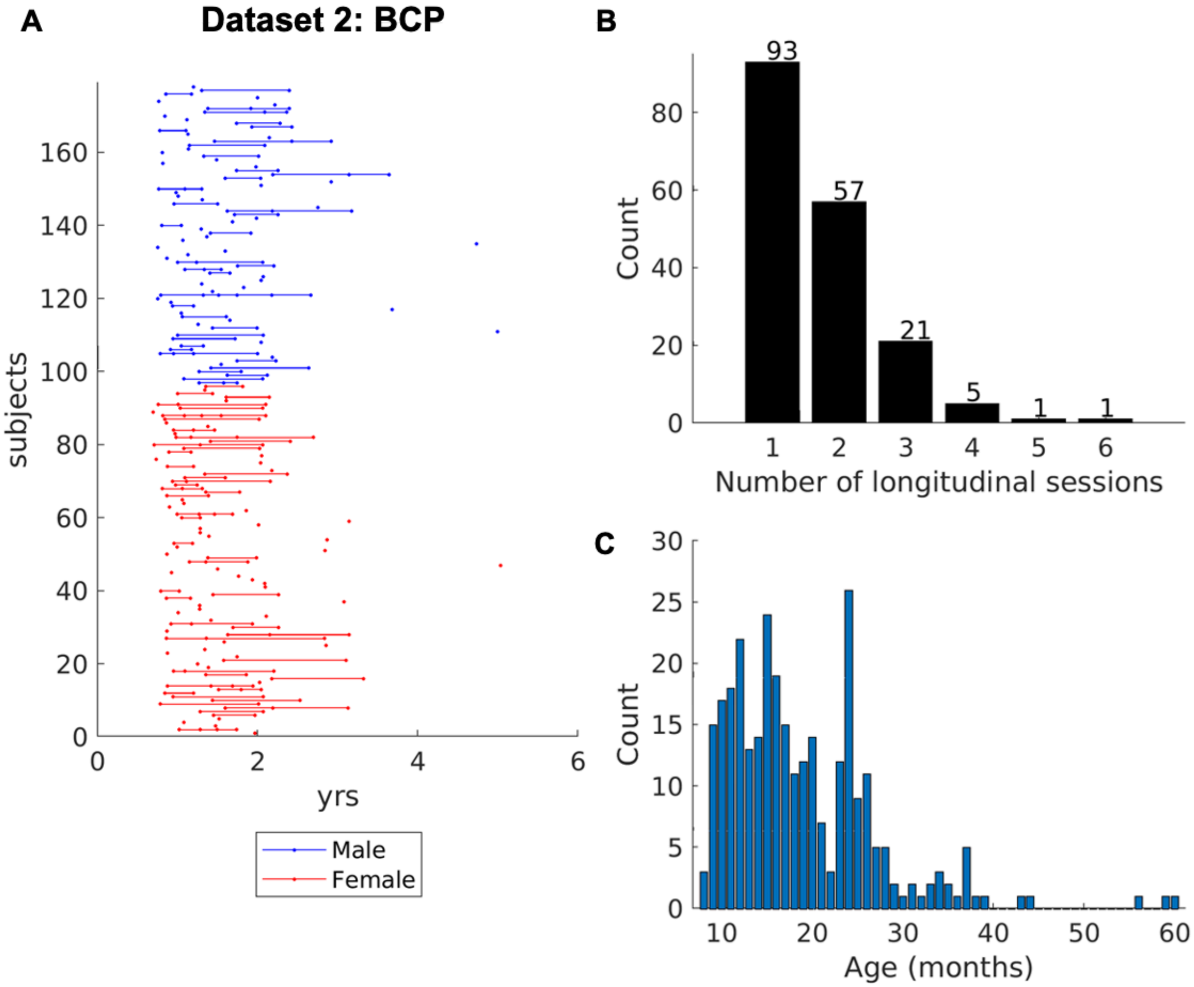

**Supplementary Figure 21**. *Dataset demographics (BCP).* A) The age at fMRI session for each participant, with participants sorted by Male (blue) and Female (red). B) Number of participants with 1-6 longitudinal sessions. C) Number of sessions each month.

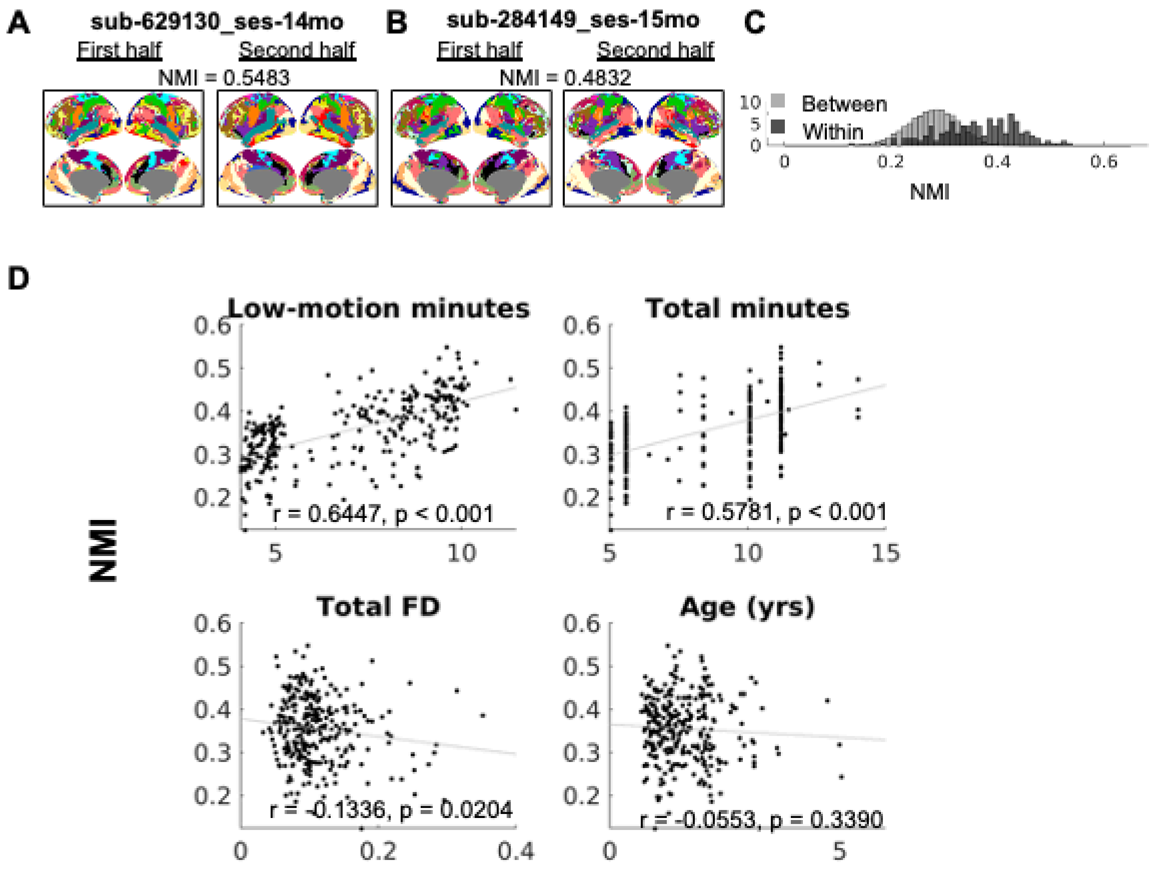


**Supplementary Figure 22**. *Individual-specific network topography for 23 networks (Dataset 2).* A-B) Individual-specific functional networks using fMRI data from two non-overlapping splits of two example participants. C) Within-session NMI compared to between-session NMI for 178 sessions from 178 unique participants. The session with the largest amount of usable, low-motion data was chosen for each individual when multiple longitudinal sessions exist for a participant. D) The relationship between NMI and the amount of low-motion data, total scan time, total framewise displacement, and age.

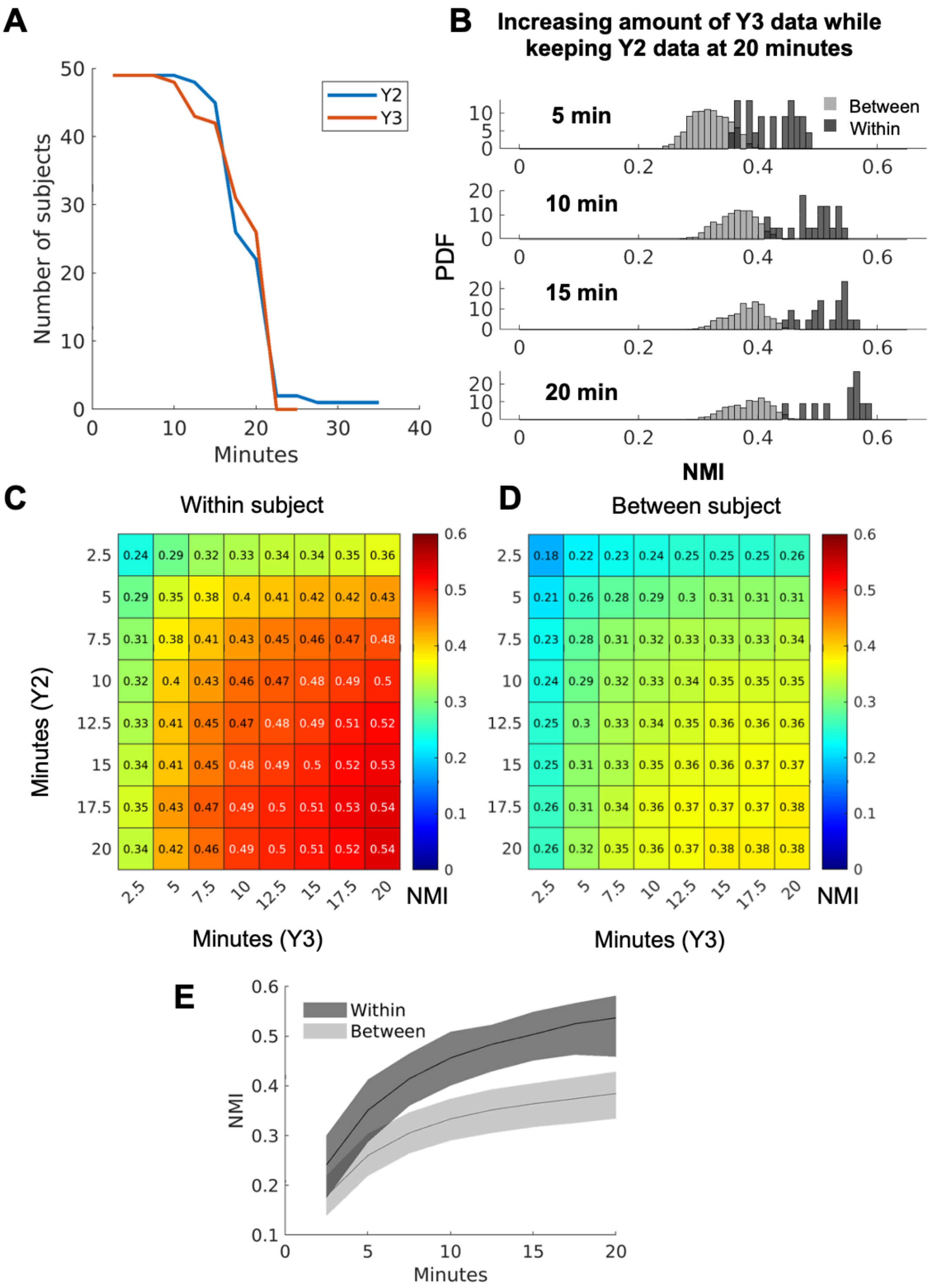
A
Number of subjects
Minutes
Y2
Y3
B
Increasing amount of Y3 data while keeping Y2 data at 20 minutes
5 min
10 min
15 min
20 min
Between
Within
PDF
NMI
C
Within subject
Minutes (Y2)
Minutes (Y3)
NMI
2.5 0.24 0.29 0.32 0.33 0.34 0.34 0.35 0.36
5 0.29 0.35 0.38 0.4 0.41 0.42 0.42 0.43
7.5 0.31 0.38 0.41 0.43 0.45 0.46 0.47 0.48
10 0.32 0.4 0.43 0.46 0.47 0.48 0.49 0.5
12.5 0.33 0.41 0.45 0.47 0.48 0.49 0.51 0.52
15 0.34 0.41 0.45 0.48 0.49 0.5 0.52 0.53
17.5 0.35 0.43 0.47 0.49 0.5 0.51 0.53 0.54
20 0.34 0.42 0.46 0.49 0.5 0.51 0.52 0.54
2.5 5 7.5 10 12.5 15 17.5 20
D
Between subject
Minutes (Y3)
NMI
2.5 0.18 0.22 0.23 0.24 0.25 0.25 0.25 0.26
5 0.21 0.26 0.28 0.29 0.3 0.31 0.31 0.31
7.5 0.23 0.28 0.31 0.32 0.33 0.33 0.33 0.34
10 0.24 0.29 0.32 0.33 0.34 0.35 0.35 0.35
12.5 0.25 0.3 0.33 0.34 0.35 0.36 0.36 0.36
15 0.25 0.31 0.33 0.35 0.36 0.36 0.37 0.37
17.5 0.26 0.31 0.34 0.36 0.37 0.37 0.37 0.38
20 0.26 0.32 0.35 0.36 0.37 0.38 0.38 0.38
2.5 5 7.5 10 12.5 15 17.5 20
E
Within
Between
NMI
Minutes

**Supplementary Figure 23**. *Effect of data amount on longitudinal functional network stability.* A) Number of subjects with at least X minutes of data. B) The distribution of NMI with increasing Y3 data while keeping the Y2 data the same. C) Mean within-subject NMI for different combinations of data amounts. D) Mean between-subject NMI for different combinations of data amounts. E) Mean within- and between-subject NMI when minutes in Y2 and Y3 are the same (i.e., diagonals in C and D). The shaded areas show 83.4% CI (commonly used for comparing two means, see Supplementary Text).

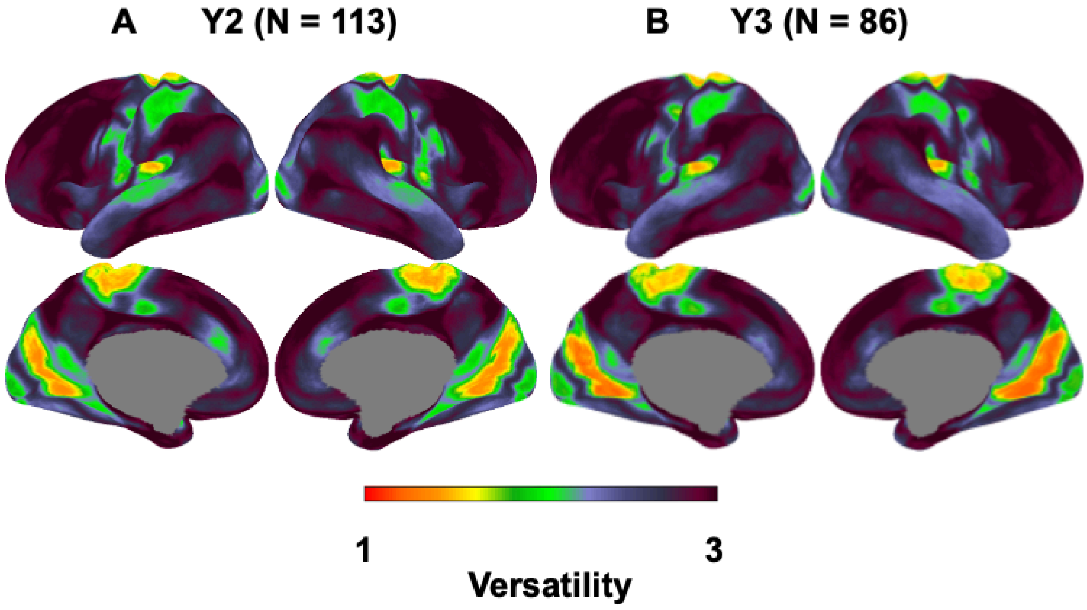


**Supplementary Figure 24**. *Nodal versatility across subjects in Dataset 1.* A) Y2. B) Y3.

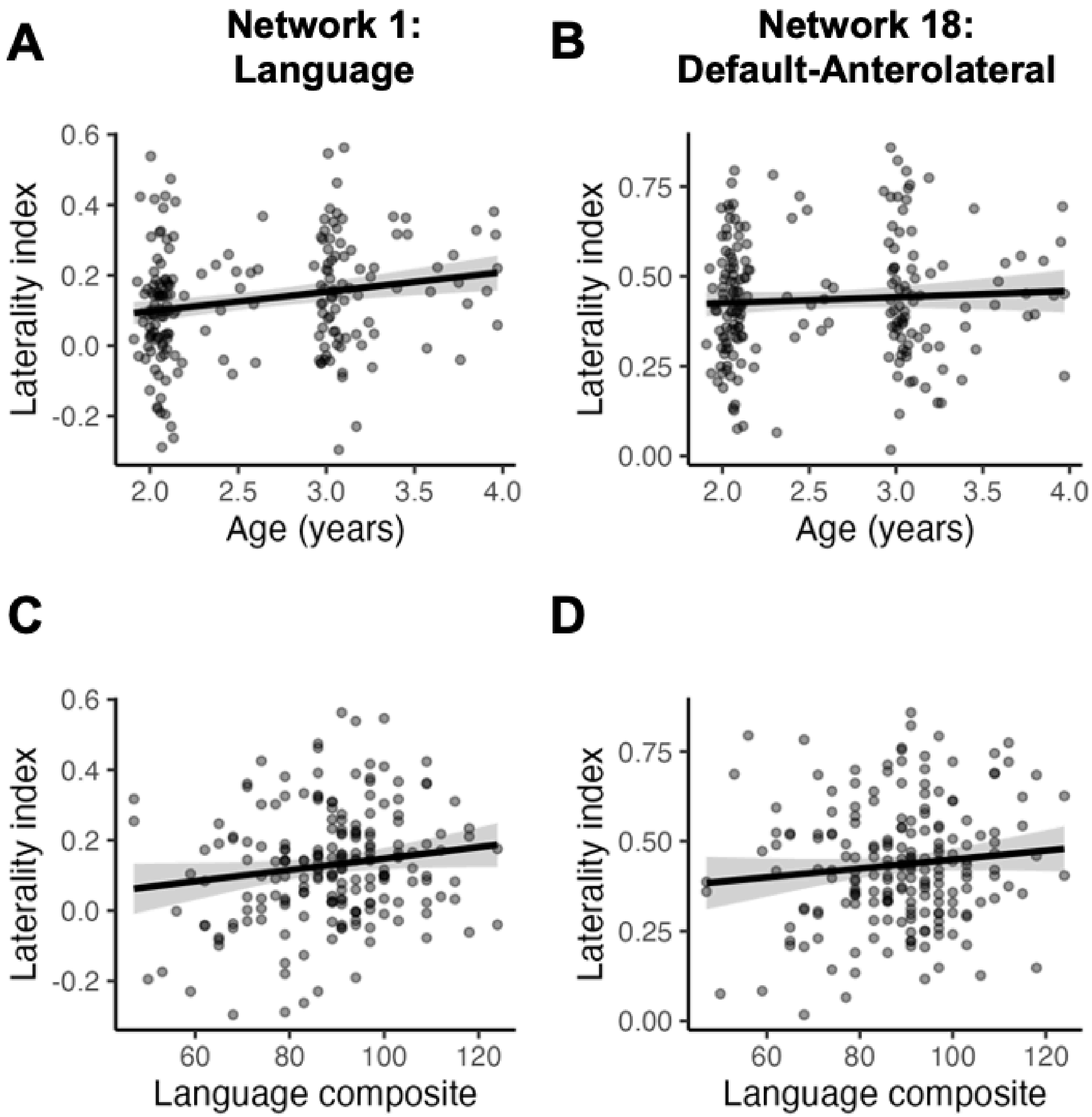


**Supplementary Figure 25**. *The relationship between functional network laterality and age or Bayley Scales of Infant and Toddler Development age-normalized composite scores.* A-B) Age, C-D) Language composite scores. Shaded band = 95% CI of fixed effect (age was held constant in C-D).

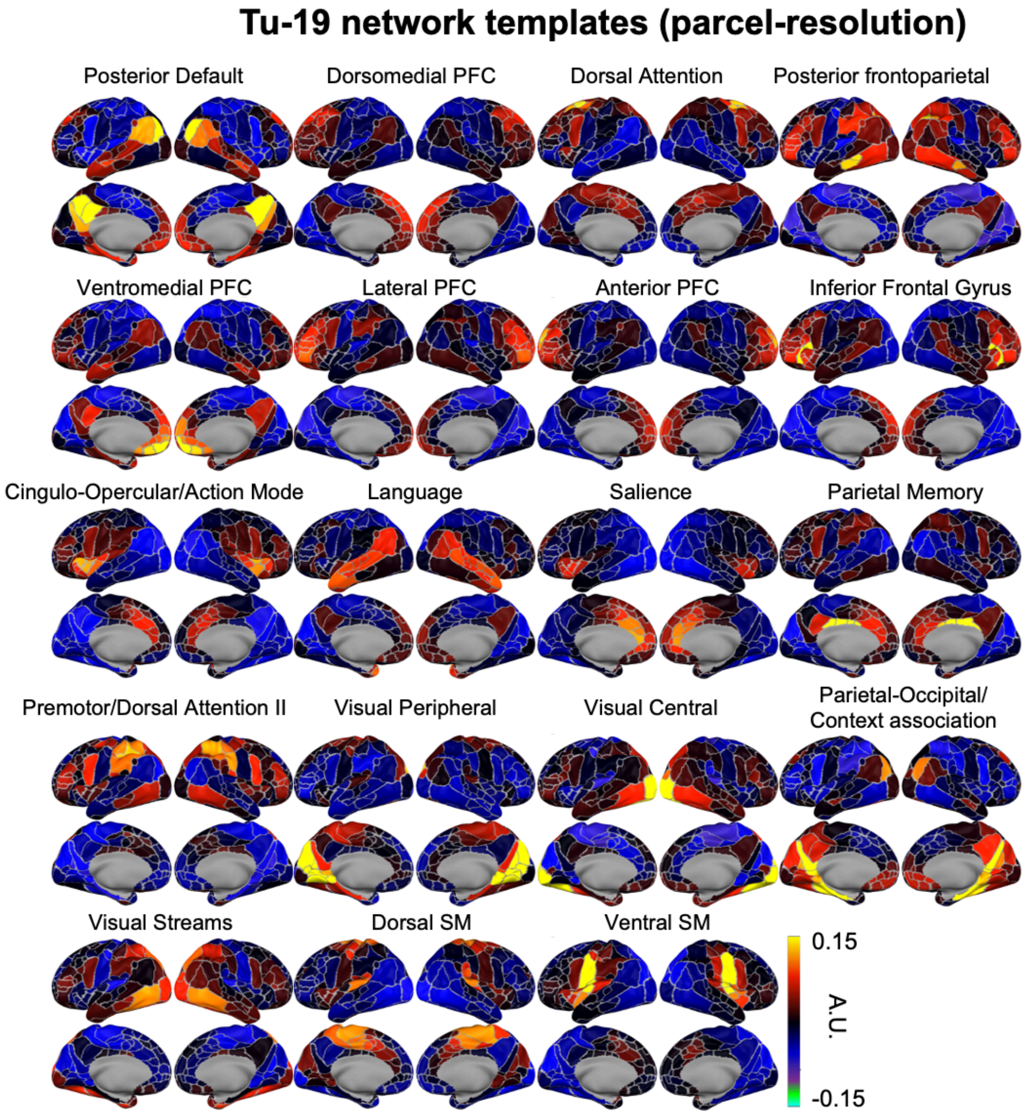


**Supplementary Figure 26**. *Tu-19 network templates at the resolution of parcels. PFC, prefrontal cortex. SM, Somatomotor.* The values represent the mean of normalized vectors (a.k.a. cluster centroids using correlational distance) in arbitrary units (A.U.).

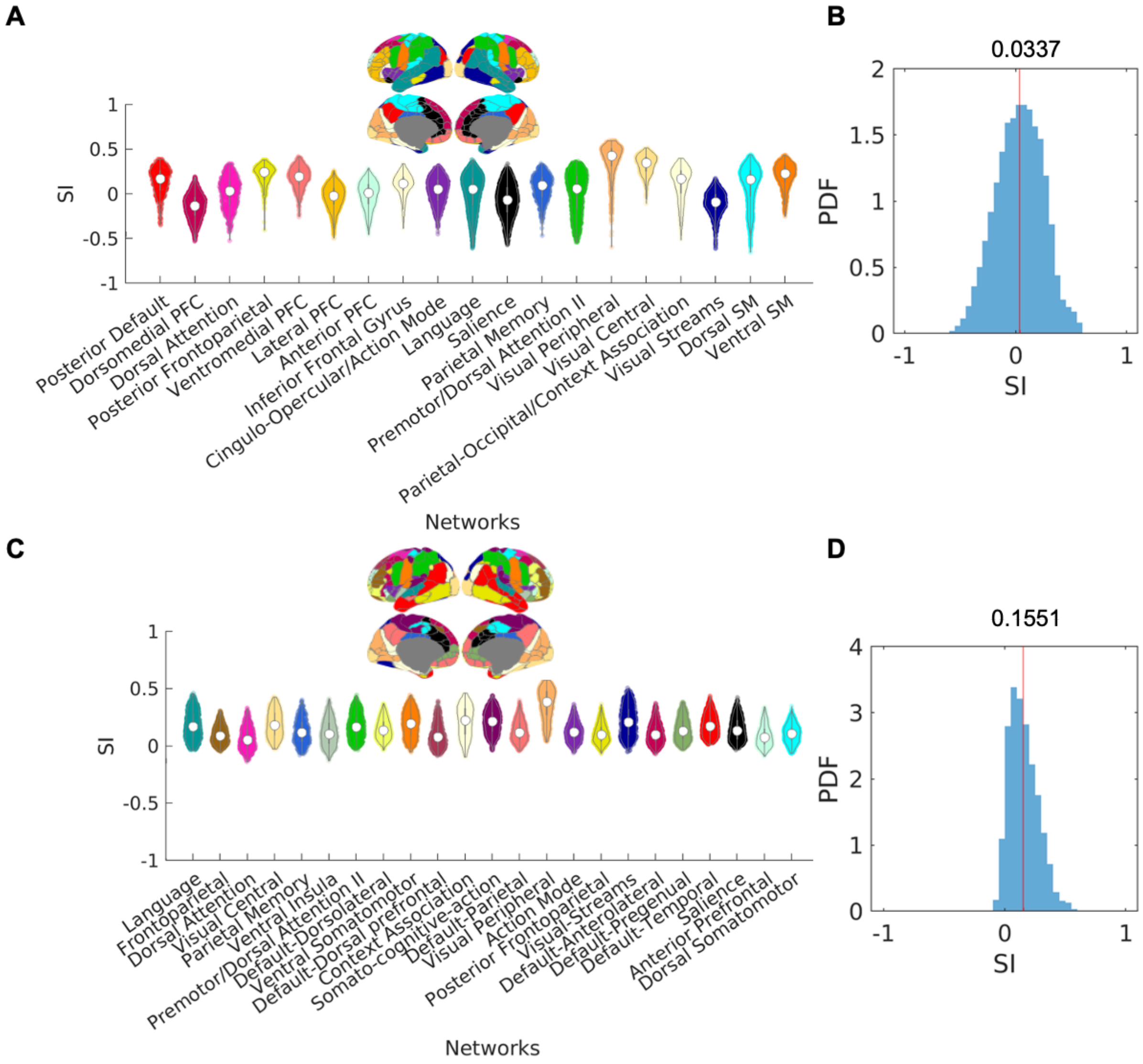
A
B
C
D
SI
PDF
Networks
0.0337
0.1551
Posterior Default
Dorsomedial PFC
Dorsal Attention
Posterior Frontoparietal
Ventromedial PFC
Lateral PFC
Anterior PFC
Inferior Frontal Gyrus
Cingulo-Opercular/Action Mode
Language
Salience
Parietal Memory
Premotor/Dorsal Attention II
Visual Peripheral
Visual Central
Parietal-Occipital/Context Association
Visual Streams
Dorsal SM
Ventral SM
Frontoparietal
Ventral Insula
Default-Dorsolateral
Ventral Somatomotor
Default-Dorsal prefrontal
Context Association
Somato-cognitive-action
Default-Parietal
Action Mode
Posterior Frontoparietal
Visual-Streams
Default-Anterolateral
Default-Pregenual
Default-Temporal
Anterior Prefrontal
Dorsal Somatomotor

**Supplementary Figure 27**. *Comparing the quality of separation between the FC profiles using the Tu-19 networks and the 23 networks from clustering concatenated FC matrices across sessions. N.B. Each data sample represents one parcel from a single individual.* A) The silhouette index (SI) for parcels in each of the Tu-19 networks. B) The SI for all parcels across all 19 networks. C-D) Same as A-B, but for the 23 networks. N.B. The population consensus was plotted for anatomical reference, but the actual network assignments differ across sessions, unlike the Tu-19 networks.

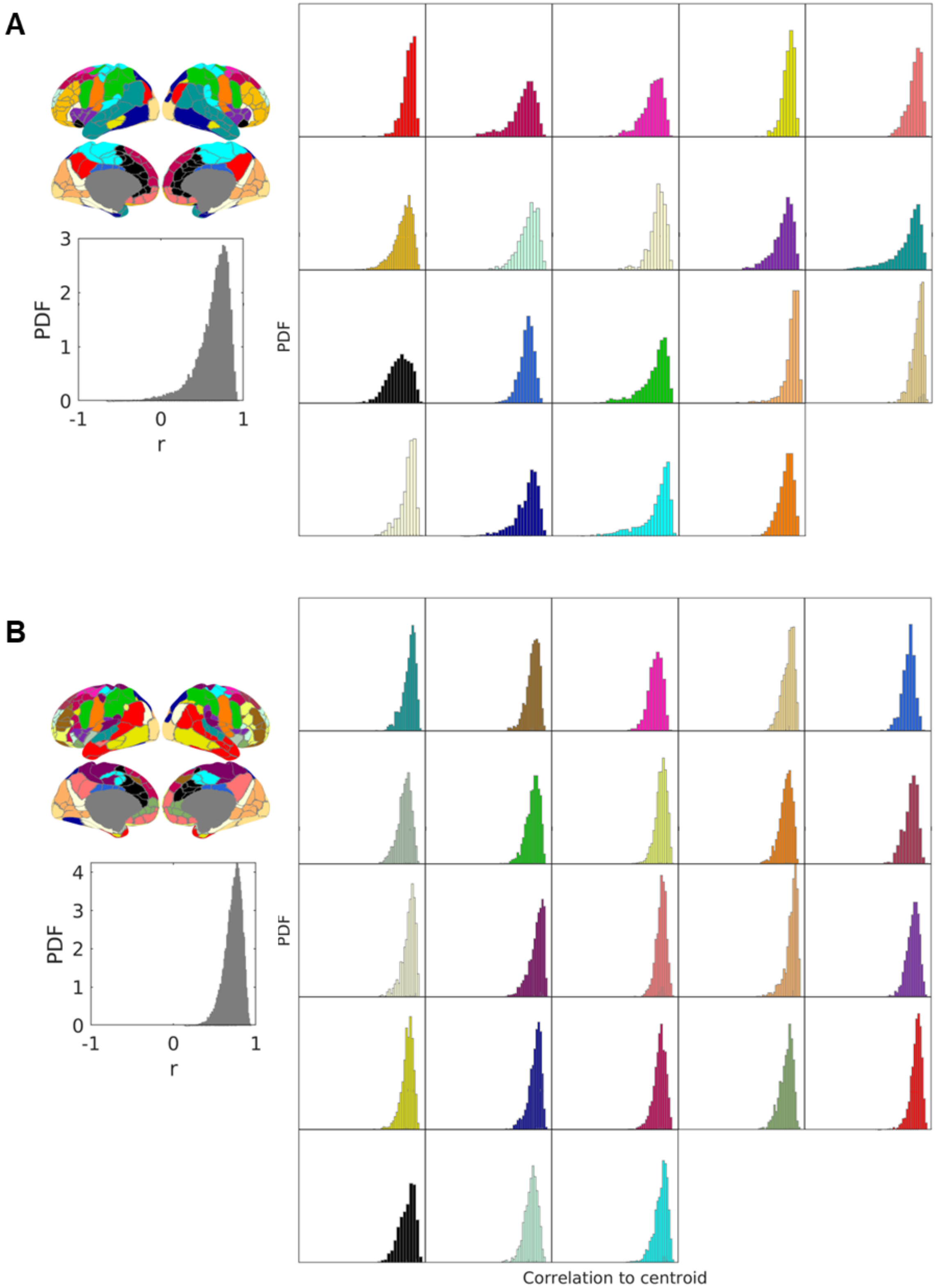
A
3
2
1
0
PDF
-1
0
1
r
PDF
B
4
3
2
1
0
PDF
-1
0
1
r
PDF
Correlation to centroid

**Supplementary Figure 28**. *Pearson's correlation between the network templates (centroids) to FC profiles from all parcels assigned to that network.* A) Tu-19 networks. B) 23 networks. N.B. The population consensus was plotted for visualization only; the actual network assignments differ across sessions, unlike the Tu-19 networks. The x-axis ranges from -1 to 1.

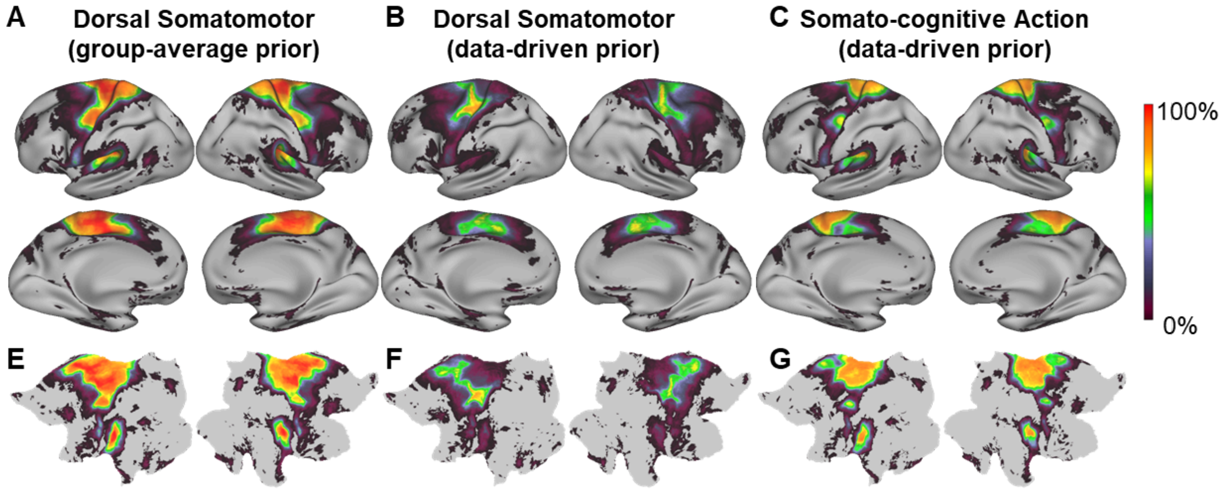


**Supplementary Figure 29.** *Network spatial topography of similar networks using group-average prior versus data-driven prior.* A) Dorsal Somatomotor (group-average prior). B) Dorsal Somatomotor (data-driven prior). C) Somato-cognitive Action (data-driven prior). D-F) Same as A-C but viewed in flat maps.

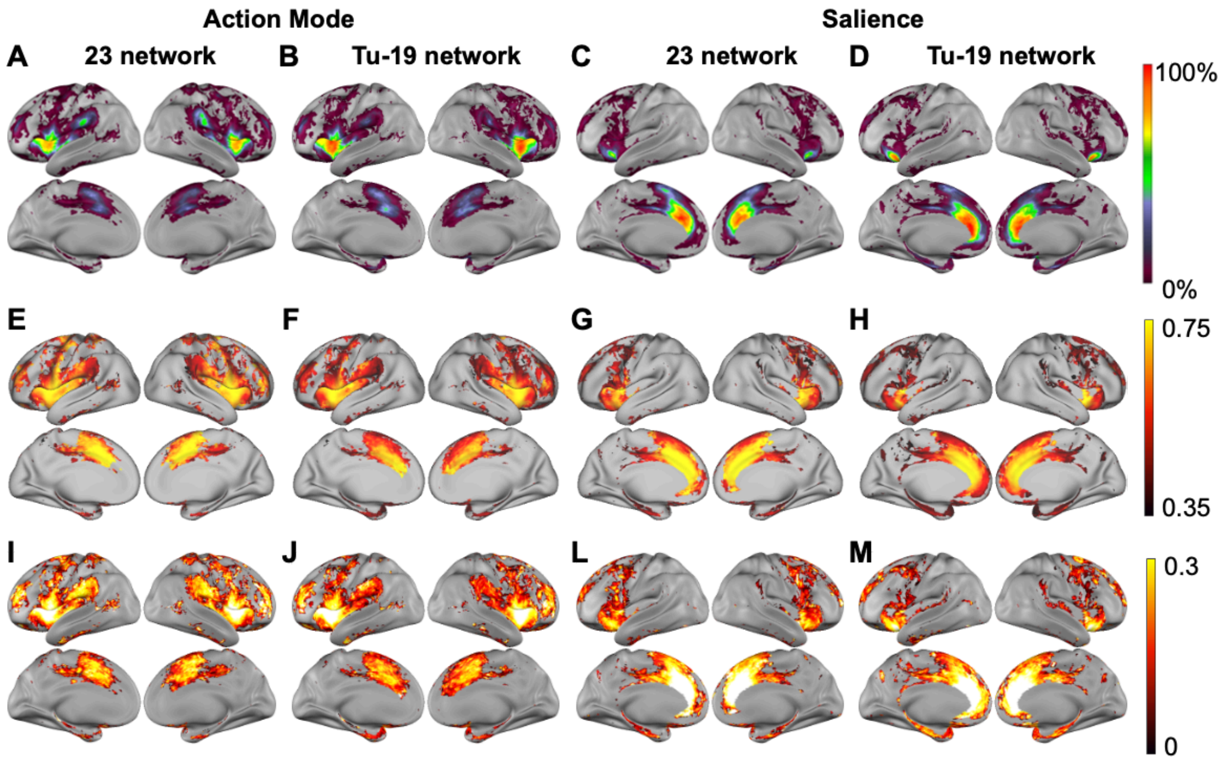


**Supplementary Figure 30.** *Comparing the network topography of the same network using different network templates*. A-B) Spatial probability E-H) Mean Pearson's correlation to the network template. I-M) Silhouette Index (Confidence) with correlational distance.

Gordon, E.M., Chauvin, R.J., Van, A.N., Rajesh, A., Nielsen, A., Newbold, D.J., Lynch, C.J., Seider, N.A., Krimmel, S.R., Scheidter, K.M., Monk, J., Miller, R.L., Metoki, A., Montez, D.F., Zheng, A., Elbau, I., Madison, T., Nishino, T., Myers, M.J., Kaplan, S., Badke D'Andrea, C., Demeter, D.V., Feigelis, M., Ramirez, J.S.B., Xu, T., Barch, D.M., Smyser, C.D., Rogers, C.E., Zimmermann, J., Botteron, K.N., Pruett, J.R., Willie, J.T., Brunner, P., Shimony, J.S., Kay, B.P., Marek, S., Norris, S.A., Gratton, C., Sylvester, C.M., Power, J.D., Liston, C., Greene, D.J., Roland, J.L., Petersen, S.E., Raichle, M.E., Laumann,

T.O., Fair, D.A., Dosenbach, N.U.F., 2023. A somato-cognitive action network alternates with effector regions in motor cortex. Nature 617. https://doi.org/10.1038/s41586-023-05964-2
Gordon, E.M., Laumann, T.O., Gilmore, A.W., Newbold, D.J., Greene, D.J., Berg, J.J., Ortega, M., Hoyt-Drazen, C., Gratton, C., Sun, H., Hampton, J.M., Coalson, R.S., Nguyen, A.L., McDermott, K.B., Shimony, J.S., Snyder, A.Z., Schlaggar, B.L., Petersen, S.E., Nelson, S.M., Dosenbach, N.U.F., 2017. Precision Functional Mapping of Individual Human Brains. Neuron 95. https://doi.org/10.1016/j.neuron.2017.07.011
Hermosillo, R.J.M., Moore, L.A., Feczko, E., Miranda-Domínguez, Ó., Pines, A., Dworetsky, A., Conan, G., Mooney, M.A., Randolph, A., Graham, A., Adeyemo, B., Earl, E., Perrone, A., Carrasco, C.M., Uriarte-Lopez, J., Snider, K., Doyle, O., Cordova, M., Koirala, S., Grimsrud, G.J., Byington, N., Nelson, S.M., Gratton, C., Petersen, S., Feldstein Ewing, S.W., Nagel, B.J., Dosenbach, N.U.F., Satterthwaite, T.D., Fair, D.A., 2024. A precision functional atlas of personalized network topography and probabilities. Nat Neurosci 27, 1000–1013. https://doi.org/10.1038/s41593-024-01596-5
Ji, J.L., Spronk, M., Kulkarni, K., Repovš, G., Anticevic, A., Cole, M.W., 2019. Mapping the human brain's cortical-subcortical functional network organization. NeuroImage 185, 35–57. https://doi.org/10.1016/j.neuroimage.2018.10.006
Kardan, O., Kaplan, S., Wheelock, M.D., Feczko, E., Day, T.K.M., Miranda-Domínguez, Ó., Meyer, D., Eggebrecht, A.T., Moore, L.A., Sung, S., Chamberlain, T.A., Earl, E., Snider, K., Graham, A., Berman, M.G., Uğurbil, K., Yacoub, E., Elison, J.T., Smyser, C.D., Fair, D.A., Rosenberg, M.D., 2022. Resting-state functional connectivity identifies individuals and predicts age in 8-to-26-month-olds. Developmental Cognitive Neuroscience 56, 101123. https://doi.org/10.1016/j.dcn.2022.101123
Knol, M.J., Pestman, W.R., Grobbee, D.E., 2011. The (mis)use of overlap of confidence intervals to assess effect modification. Eur J Epidemiol 26, 253–254. https://doi.org/10.1007/s10654-011-9563-8
Kong, R., Li, J., Orban, C., Sabuncu, M.R., Liu, H., Schaefer, A., Sun, N., Zuo, X.-N., Holmes, A.J., Eickhoff, S.B., Yeo, B.T.T., 2019. Spatial Topography of Individual-Specific Cortical Networks Predicts Human Cognition, Personality, and Emotion. Cerebral Cortex 29, 2533–2551. https://doi.org/10.1093/cercor/bhy123
Kong, R., Spreng, R.N., Xue, A., Betzel, R.F., Cohen, J.R., Damoiseaux, J.S., De Brigard, F., Eickhoff, S.B., Fornito, A., Gratton, C., Gordon, E.M., Holmes, A.J., Laird, A.R., Larson-Prior, L., Nickerson, L.D., Pinho, A.L., Razi, A., Sadaghiani, S., Shine, J.M., Yendiki, A., Yeo, B.T.T., Uddin, L.Q., 2025. A network correspondence toolbox for quantitative evaluation of novel neuroimaging results. Nat Commun 16, 2930. https://doi.org/10.1038/s41467-025-58176-9
Laumann, T.O., Gordon, E.M., Adeyemo, B., Snyder, A.Z., Joo, S.J., Chen, M.-Y., Gilmore, A.W., McDermott, K.B., Nelson, S.M., Dosenbach, N.U.F., Schlaggar, B.L., Mumford, J.A., Poldrack, R.A., Petersen, S.E., 2015. Functional System and Areal Organization of a Highly Sampled Individual Human Brain. Neuron 87, 657–670. https://doi.org/10.1016/j.neuron.2015.06.037
Payton, M.E., Greenstone, M.H., Schenker, N., 2003. Overlapping confidence intervals or standard error intervals:  What do they mean in terms of statistical significance? Journal of Insect Science 3, 34. https://doi.org/10.1093/jis/3.1.34
Percival, C., Woodward, T., 2025. Cognitive modes detectable with task-based fMRI. https://doi.org/10.5281/zenodo.4274397
Salo, T., Yarkoni, T., Nichols, T.E., Poline, J.-B., Bilgel, M., Bottenhorn, K.L., Jarecka, D., Kent, J.D., Kimbler, A., Nielson, D.M., Oudyk, K.M., Peraza, J.A., Pérez, A., Reeders, P.C., Yanes, J.A., Laird, A.R., 2023. NiMARE: Neuroimaging Meta-Analysis Research Environment. Aperture Neuro 3, 1–32. https://doi.org/10.52294/001c.87681

Shen, X., Tokoglu, F., Papademetris, X., Constable, R.T., 2013. Groupwise whole-brain parcellation from resting-state fMRI data for network node identification. NeuroImage 82, 403–415. https://doi.org/10.1016/j.neuroimage.2013.05.081

Tu, J.C., Myers, M.J., Li, W., Li, J., Wang, X., Dierker, D., Day, T.K.M., Snyder, A., Latham, A., Kenley, J.K., Sobolewski, C.M., Wang, Y., Labonte, A.K., Feczko, E., Kardan, O., Moore, L.A., Sylvester, C.M., Fair, D.A., Elison, J.T., Warner, B.B., Barch, D.M., Rogers, C.E., Luby, J.L., Smyser, C.D., Gordon, E.M., Laumann, T.O., Eggebrecht, A.T., Wheelock, M.D., 2025. The generalizability of cortical area parcellations across early childhood. Cerebral Cortex 35, bhaf116. https://doi.org/10.1093/cercor/bhaf116

Winkler, A.M., Ridgway, G.R., Webster, M.A., Smith, S.M., Nichols, T.E., 2014. Permutation inference for the general linear model. Neuroimage 92, 381–397. https://doi.org/10.1016/j.neuroimage.2014.01.060

Yan, X., Kong, R., Xue, A., Yang, Q., Orban, C., An, L., Holmes, A.J., Qian, X., Chen, J., Zuo, X.-N., Zhou, J.H., Fortier, M.V., Tan, A.P., Gluckman, P., Chong, Y.S., Meaney, M.J., Bzdok, D., Eickhoff, S.B., Yeo, B.T.T., 2023. Homotopic local-global parcellation of the human cerebral cortex from resting-state functional connectivity. NeuroImage 273, 120010. https://doi.org/10.1016/j.neuroimage.2023.120010

Yarkoni, T., Poldrack, R.A., Nichols, T.E., Van Essen, D.C., Wager, T.D., 2011. Large-scale automated synthesis of human functional neuroimaging data. Nat Methods 8, 665–670. https://doi.org/10.1038/nmeth.1635
